# Plankton energy flows using a global size-structured and trait-based model

**DOI:** 10.1101/2022.02.01.478546

**Authors:** Gabriela Negrete-García, Jessica Y. Luo, Matthew C. Long, Keith Lindsay, Michael Levy, Andrew D. Barton

**Affiliations:** Scripps Institute of Oceanography, University of California San Diego, La Jolla, CA; Climate & Global Dynamics Laboratory, National Center for Atmospheric Research, Boulder, CO; Geophysical Fluid Dynamics Laboratory, National Oceanic and Atmospheric Administration, Princeton NJ; Department of Ecology, Behavior and Evolution, University of California, San Diego, CA, USA

**Keywords:** plankton communities, trait-based models, phytoplankton, zooplankton, Earth system modeling

## Abstract

Plankton community models are critical tools for understanding the processes that shape marine plankton communities, how plankton communities impact biogeochemical cycles, and the feedbacks between community structure and function. Here, using the flexible Marine Biogeochemistry Library (MARBL), we present the Size-based Plankton Ecological TRAits (MARBL-SPECTRA) model, which is designed to represent a diverse plankton community while remaining computationally tractable. MARBL-SPECTRA is composed of nine phytoplankton and six zooplankton size classes represented using allometric scaling relationships for physiological traits and interactions within multiple functional types. MARBL-SPECTRA is embedded within the global ocean component of the Community Earth System Model (CESM) and simulates large-scale, emergent patterns in phytoplankton growth limitation, plankton phenology, plankton generation time, and trophic transfer efficiency. The model qualitatively reproduces observed global patterns of surface nutrients, chlorophyll biomass, net primary production, and the biogeographies of a range of plankton size classes. In addition, the model simulates how predator:prey dynamics and trophic efficiency vary across gradients in total ecosystem productivity. Shorter food chains that export proportionally more carbon from the surface to the ocean interior occur in productive, eutrophic regions, whereas in oligotrophic regions, the food chains are relatively long and export less organic matter from the surface. The union of functional type modeling with size-resolved, trait-based modeling approaches allows MARBL-SPECTRA to capture both large-scale elemental cycles and the structure of planktonic food webs affecting trophic transfer efficiency.

## 1. Introduction

Plankton communities, and consequently their ecosystem and biogeochemical functions, exhibit pronounced spatial and temporal variations. Relatively large phytoplankton and their mesozooplankton consumers are most abundant in eutrophic regions of the ocean, such as coastal upwelling zones and subpolar gyres, but are relatively scarce in oligotrophic regions, such as the subtropical gyres (Hirata et al., 2011; James et al., 2022). Eutrophic regions have shorter, more efficient food chains compared with oligotrophic regions (Ryther, 1969), and exhibit relatively high flux of carbon and other elements from the ocean surface to depth (Henson et al., 2012; Falkowski et al., 1998; Guidi et al., 2009). The average trophic position of planktonic consumers is consequently higher in oligotrophic compared with eutrophic conditions (Décima, 2022). The size structure of the plankton community, often summarized by the slope of the log-log relationship between body size and abundance (Cermeño et al., 2006), summarizes these key ecological differences across environmental gradients, with typically steeper negative slopes indicative of relatively few larger plankton found in oligotrophic compared to eutrophic conditions (Marañón, 2015; Ward et al., 2012). Moreover, plankton communities exhibit characteristic and often pronounced phenology driven by seasonal changes in environment, organism traits, individual species’ generation times (Cole, 1954), and biotic interactions (Margalef, 1978), changes which have consequences for temporal variations in biogeochemical (e.g., Rynearson et al. (2013)) and ecosystem function (e.g., Cushing (1990)). Therefore, the mechanistic representations of the spatial distributions of plankton of different sizes, phenology, and food web length are essential for understanding and predicting the function of oceanic ecosystems and global biogeochemical cycles.

Coupled physical-biogeochemical models have long been used to better understand the processes that shape phytoplankton dynamics and distributions in the ocean. Historically, many marine plankton community models were constructed from a common nutrient-phytoplankton-zooplankton-detritus (NPZD) structure (Evans and Parslow, 1985; Fasham et al., 1990; Franks, 2002). Although these models simplify substantial biological complexity, when coupled to ocean circulation models, they provided large-scale estimates of total phytoplankton and zooplankton biomass and biologically-mediated carbon fluxes (Six and Maier-Reimer, 1996). More recent “intermediate complexity” marine ecosystem models (Stock et al., 2014a; Moore et al., 2004, 2013b; Aumont et al., 2015; Yool et al., 2013) coupled into Earth system models have been successful at simulating large-scale biogeographical variation in the efficiency of the biological pump and climate effects on ocean ecosystems through a number of mechanisms, including shifts in primary productivity, ocean acidification, and deoxygenation. These marine ecosystem models typically include a few key plankton functional types and/or species adequate for simulating broad plankton biogeography and biogeochemical interactions such as variations in export efficiency. However, these models typically do not resolve many plankton functional types and sizes within types, and are consequently limited for studying the complex dynamics and interactions of plankton communities and linkages to biogeochemical cycles and ecosystems.

Trait-based models are a mechanistic approach for increasing model diversity and ecological realism (Ward et al., 2012; Bruggeman and Kooijman, 2007; Follows et al., 2007; Dutkiewicz et al., 2019). Instead of simulating a few species or generic types of plankton, trait-based models resolve a higher diversity of organisms with distinct physiological and interaction traits, as well as trade-offs between these traits (Litchman et al., 2007). Trait-based models have been used to study the mechanisms shaping plankton biogeography, size structure, and diversity (Barton et al., 2010; Ward et al., 2012; Banas, 2011; Acevedo-Trejos et al., 2015; Monteiro et al., 2011; Follows et al., 2007). Allometric scaling relationships, which describe variations in traits across body size (e.g. Tang, 1995; Hansen et al., 1997; Finkel et al., 2010; Marañón, 2015), are useful for reducing the number of model parameters (e.g. Fuchs and Franks, 2010; Ward et al., 2012; Taniguchi et al., 2014; Dutkiewicz et al., 2015a; Heneghan et al., 2020; Stock et al., 2020). Such scaling relationships allow the parameterization of models that predict how optimal size, or community size structure, should vary with nutrient affinities (Edwards et al., 2012), light (Taguchi, 1976; Edwards et al., 2015b), maximum biosynthesis rates and respiration rates (Kiørboe and Hirst, 2014), clearance rates (Kiørboe and Hirst, 2014), predator-prey mass ratios (Hansen et al., 1994), and predation risk from larger organisms (Hirst and Kiørboe, 2002). While organism traits vary across cell size, they also vary across functional group. For example, nitrogen fixation by diazotrophs is a costly, but important source of new nitrogen in low nutrient, oligotrophic gyres that sustains key biogeochemical processes such as carbon export (Karl et al., 1997). Diatoms produce silica frustules that serves as a defense against predation (Finkel and Kotrc, 2010), represent the majority of the biogenic silica production, and serve as a significant control on the ocean silica cycle. Moreover, key organism traits, such as growth and prey ingestion rates, vary across functional groups (Ward et al., 2012; Kiørboe and Hirst, 2014). Therefore, it is critical for coupled plankton community and biogeochemical models to resolve multiple functional groups and organism sizes within groups (e.g., Dutkiewicz et al. (2015b); Mangolte et al. (2022); Dutkiewicz et al. (2021)).

Here, we describe a new size-structured plankton community and biogeochemical model, called MARBL-SPECTRA, that leverages advances in trait-based ecological modeling while remaining computationally tractable within coupled climate simulations. We leverage the Marine Biogeochemistry Library (MARBL) (Long et al., 2021), a configurable ocean biogeochemical model that has been coupled to the Community Earth System Model (CESM), in order to implement the Size-based Plankton ECological TRAits (SPECTRA) model. SPECTRA builds on the MARBL-CESM version 2.1 default ecosystem by expanding the total number of phytoplankton and zooplankton types and functional groups, using allometric scaling relationships to tie organismal traits and interactions to body size and functional groups. MARBL-SPECTRA includes nine phytoplankton groups belonging to four different plankton functional types (picoplankton, mixed phytoplankton, diatoms, and diazotrophs). It also includes six zooplankton groups including two microzooplankton (<200 *μ*m ESD) and four mesozooplankton size classes (between 0.2 mm and 20 mm). In the Methods, we describe the features and assumptions of MARBL-SPECTRA, and then use the model to explore large-scale, emergent patterns in plankton biogeography, plankton size structure, phytoplankton growth limitation, plankton phenology, plankton generation time, and trophic dynamics (in Results). We describe model sensitivity analyses conducted to understand the impact on plankton biomass of key phytoplankton physiological and zooplankton grazing parameters (Methods). We compared the model results to a suite of biogeochemical and plankton distribution observations to assess MARBL-SPECTRA’s ability to capture global-scale patterns in macronutrients, plankton biogeography, chlorophyll, and mesozooplankton biomass. We used the model to examine spatial gradients in the linkages between carbon export from the ocean surface and size structure and trophic dynamics. SPECTRA builds upon considerable, recent developments in plankton community modeling and leverages the flexible MARBL environment to create a coupled plankton community and biogeochemical model that can be used to study complex ecological dynamics and interactions with ocean biogeochemical cycles.

## 2. Methods

### 2.1. Size-based Plankton ECological TRAits (SPECTRA) Model

The Size-based Plankton ECological TRAits (SPECTRA) planktonic community model is implemented using the Marine Biogeochemistry Library (MARBL) (Long et al., 2021), which is the ocean biogeochemical component within the Community Earth System Model (CESM). MARBL is designed to allow for a flexible number of plankton functional types, and in its default configuration, invokes an updated version of the marine ecosystem of its predecessor, the Biogeochemistry Elemental Cycle (BEC) model (Moore et al., 2001, 2004, 2013b). MARBL-SPECTRA is a new configuration of MARBL that resolves nine phytoplankton (Fig. 1) ranging in size from 0.47 *μ*m to 300 *μ*m in equivalent spherical diameter (ESD; (Fig. 1)). The nine model phytoplankton include one picoplankton, one diazotroph, three sizes of diatoms, and four sizes of mixed phytoplankton. Size classes were chosen such that: 1) within each phytoplankton group, characteristic size (geometric mean of the size range) was evenly spaced on a *log*_10_ scale, and 2) size classes across functional types were overlapping but not identical. The picoplankton group is analogous to *Prochlorococcus* and *Synechococcus* with a characteristic size of 0.89 *μ*m ESD. Diazotrophs fix nitrogen and have a characteristic size of 6.2 *μ*m ESD. Diatoms, the silicifiers in the community, range in size between 20 *μ*m to 200 *μ*m ESD. The mixed phytoplankton size ranges from 1.7 *μ*m to 300 *μ*m ESD, and represent solitary protists not included in the other functional groups, such as picoeukaryotes and autotrophic dinoflagellates. Within the mixed phytoplankton group, implicit calcifiers (including coccolithophores) are represented by size classes between 3 *μ*m and 25 *μ*m to encompass the main species of coccolithophores (i.e., *Emiliania huxleyi*) (Aloisi, 2015). Phytoplankton ESD was converted to carbon biomass according to carbon:biovolume (C:BV) relationships, for picoplankton (Bertilsson et al., 2003), small nanoplankton (Reynolds, 2006), diatoms, and other non-diatom phytoplankton (Menden-Deuer and Lessard, 2000). The traits and parameters for each model phytoplankton are determined by their body size and functional group, which we describe in greater detail below.

**Figure 1:**
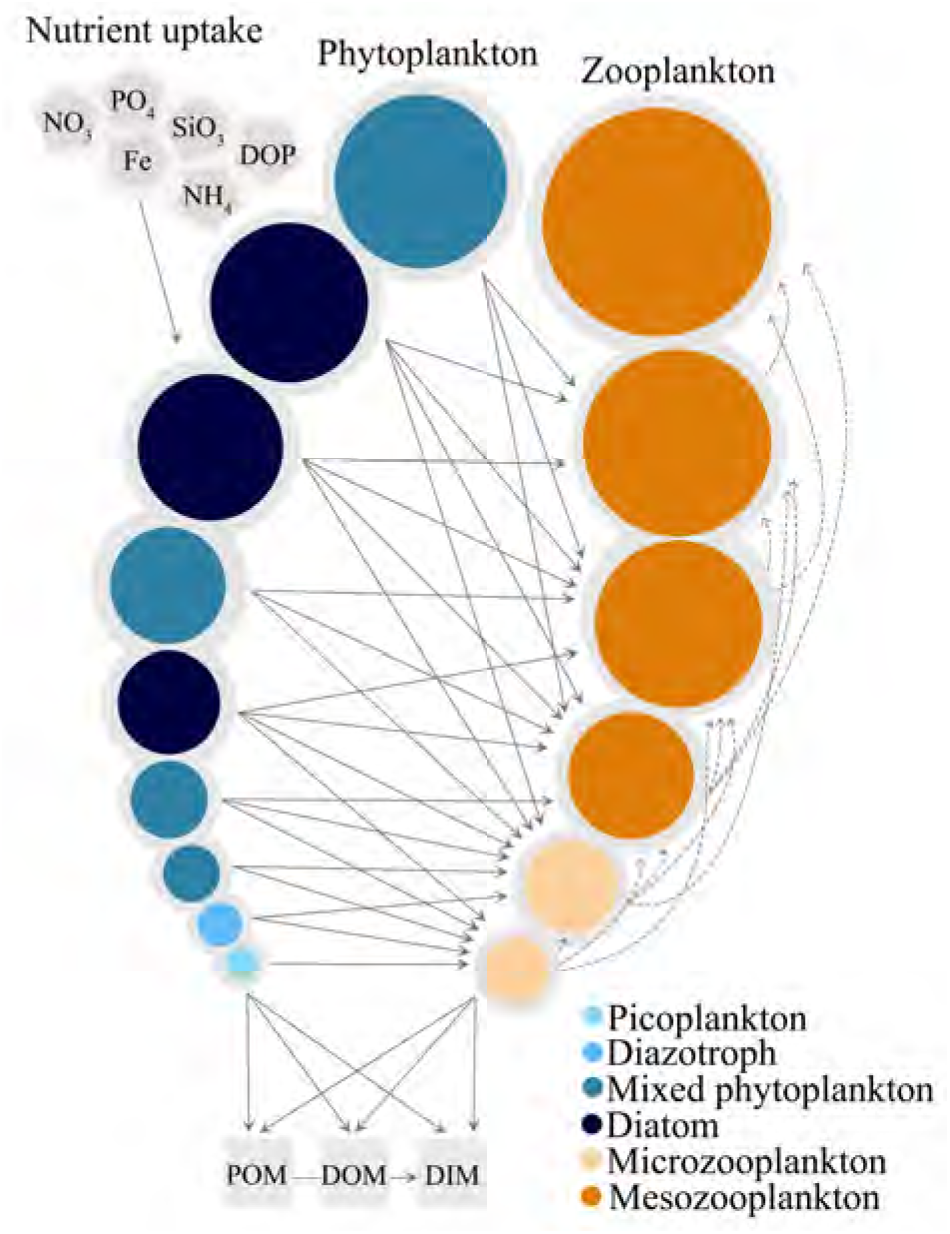
Schematic representation of MARBL-SPECTRA Model. The plankton community is composed of nine phytoplankton groups belonging to four different functional types; picoplankton (light blue), diazotrophs (sky blue), mixed phytoplankton (medium blue), and diatoms (dark blue), and six zooplankton groups composed of microzooplankton (light orange), and mesozooplankton (bright orange). Inorganic nutrients are taken up by phytoplankton (SiO_3_ is only taken up by diatoms) who are grazed by zooplankton. Larger circles indicate larger organisms, but the circles are not to scale. Straight arrows indicate phytoplankton consumption by zooplankton, while dotted arrows indicate zooplankton consumption by zooplankton. Mortality and aggregation transfer living organic material into sinking particulate and dissolved organic detritus. The fluxes to particulate organic matter (POM), dissolved organic matter (DOM), and dissolved inorganic matter (DIM) pools are shown as arrows from phytoplankton and zooplankton groups.

MARBL-SPECTRA includes six zooplankton ranging in size from 20 *μ*m to 20 mm ESD (Fig. 1), with each zooplankton consuming multiple phytoplankton and zooplankton prey types. The smallest two zooplankton (<200 *μ*m ESD) are heterotrophic organisms commonly referred to as microzoo-plankton. The smallest microzooplankton consumes only small phytoplankton, whereas the larger microzooplankton consumes both, small phytoplankton groups and the smallest microzooplankton. Mesozooplankton (between 0.2 mm and 20 mm) correspond to the largest four zooplankton size classes (zoo3-zoo6) and include a range of organisms such as copepods, krill, chaetognaths, and some gelatinous zooplankton. The first three mesozooplankton size classes are omnivorous, able to consume a range of phytoplankton and zooplankton prey, while the largest mesozooplankton is carnivorous, ranging from 6.3 to 20 mm in size. These feeding relationships between predators and prey are depicted by the feeding preference coefficient, composed of a predator-to-prey optimal size ratio of 12.5:1 and maximum and minimum of 50:1 and 1:1, respectively (Section 2.3.2) (Law et al., 2009; Fuchs and Franks, 2010; Taniguchi et al., 2014; Heneghan et al., 2020). Notably excluded from the mesozooplankton are gelatinous zooplankton like salps and pyrosomes with extremely wide predator-to-prey size ratios (e.g., between 10,000:1 and 50:1) (Conley et al., 2018). Zooplankton ESD was converted to carbon biomass using microzooplankton values from Menden-Deuer and Lessard (2000) and general non-gelatinous mesozooplankton values from Pitt et al. (2013).

MARBL-SPECTRA leverages MARBL’s flexible ecosystem configuration, which represents phytoplankton types *P_i_* and grazers *Z_j_*, where *P_i_* (mmol C m^−3^) is the phytoplankton biomass of the of the *i^th^* phytoplankton type, and *Z_j_* (mmol C m^−3^) is the zooplankton biomass of the *j^th^* zooplankton type. The rate of change of biomass for the *i^th^* phytoplankton is a balance of growth and losses to grazing (predation), mortality, and aggregation, in addition to physical transport processes not shown here. Key model symbols and units are summarized in Tables 1, 2, and 3. See Long et al. (2021) for a comprehensive presentation of the plankton community in CESM, version 2.

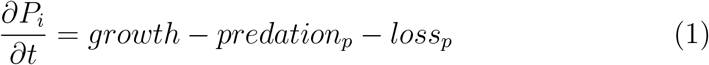

**Table 1:** Size-independent biological parameters.

| Parameter | Symbol | Value | Units |
| --- | --- | --- | --- |
| Phytoplankton activation energy | $Ea$ | 0.32* | $eV$ |
| Picoplankton activation energy | $Ea^{PP}$ | 0.42* | $eV$ |
| Zooplankton activation energy | $Ea^Z$ | 0.55<sup>†</sup> | $eV$ |
| Phytoplankton “quadratic mortality” rate | $a_i$ | 0.035 | $\text{mmol C}^{-1} \text{ m}^3 \text{ d}^{-1}$ |
| Phytoplankton linear mortality scaling | $m_i$ | 0.03 | $\text{d}^{-1}$ |
| Diatom linear mortality scaling | $m_{diat}$ | 0.02 | $\text{d}^{-1}$ |
| Grazing half-saturation coefficient | $K^P$ | 1.1 | $\text{mmol m}^{-3}$ |

\* Kremer et al. (2017).
<sup>†</sup> Lpez-Urrutia et al. (2006).

**Table 2:** Size-dependent phytoplankton biological parameters and scaling coefficients (*αV^β^*), where V is volume of each phytoplankton, *α* is a scaling constant, and *β* is an exponent describing the size dependence.

| Parameter | Symbol | Picoplankton |  | Mixed phytoplankton |  | Diatoms |  | Diazotroph |  | Units |
| --- | --- | --- | --- | --- | --- | --- | --- | --- | --- | --- |
| | | $\alpha$ | $\beta$ | $\alpha$ | $\beta$ | $\alpha$ | $\beta$ | $\alpha$ | $\beta$ | |
| C-specific rate of photosynthesis | $PC_i^{ref}$ | 1.7 | | 2.5 | -0.14<sup>†</sup> | 5.4 | -0.14<sup>†</sup> | 1.8 | | d<sup>-1</sup> |
| Initial slope of the photosynthesis-irradiance curve | $\alpha_i^{chl}$ | 0.56 | -0.15 | 0.56 | -0.15 | 0.67<sup>‡</sup> | -0.12<sup>‡</sup> | 0.56 | -0.15 | mmol C m<sup>2</sup> (mg Chl W d)<sup>-1</sup> |
| Chlorophyll to C ratio | $\theta_i^C$ | 0.12* | | 0.28* | 0.026* | 0.28* | 0.026* | 0.28* | 0.026* | mg Chl mmol C<sup>-1</sup> |
| Half-saturation concentration NO<sub>3</sub> | $K_{NO_3}$ | 0.22 | 0.30<sup>◇</sup> | 0.22 | 0.30<sup>◇</sup> | 0.20 | 0.30<sup>◇</sup> | 2 | | mmol NO<sub>3</sub> m<sup>-3</sup> |
| Half-saturation concentration NH<sub>4</sub> | $K_{NH_4}$ | 0.020 | 0.30 | 0.020 | 0.30 | 0.020 | 0.30 | 0.20 | | mmol N m<sup>-3</sup> |
| Half-saturation concentration PO<sub>4</sub> | $K_{PO_4}$ | 0.0060 | 0.30 | 0.0060 | 0.30 | 0.0060 | 0.30 | 0.006 | 0.30 | mmol PO<sub>4</sub> m<sup>-3</sup> |
| Half-saturation concentration Fe | $K_{Fe}$ | 0.60e<sup>-5</sup> | 0.30 | 0.60e<sup>-5</sup> | 0.3 | 0.6e<sup>-5</sup> | 0.30 | 0.60e<sup>-5</sup> | 0.30 | mmol Fe m<sup>-3</sup> |
| Half-saturation concentration DOP | $K_{DOP}$ | 0.080 | 0.30 | 0.080 | 0.30 | 0.080 | 0.30 | 0.080 | 0.30 | mmol DOP m<sup>-3</sup> |
| Half-saturation concentration SiO<sub>3</sub> | $K_{SiO_3}$ | | | | | 0.035 | 0.30 | | | mmol SiO<sub>3</sub> m<sup>-3</sup> |

\* matched with observed values from Hartmann et al. (2014); Li et al. (2010); Geiderlo et al. (1986); Geider et al. (1998).
<sup>†</sup> Marañón et al. (2013).
<sup>‡</sup> Edwards et al. (2015b).
◇ Edwards et al. (2012).

**Table 3:**
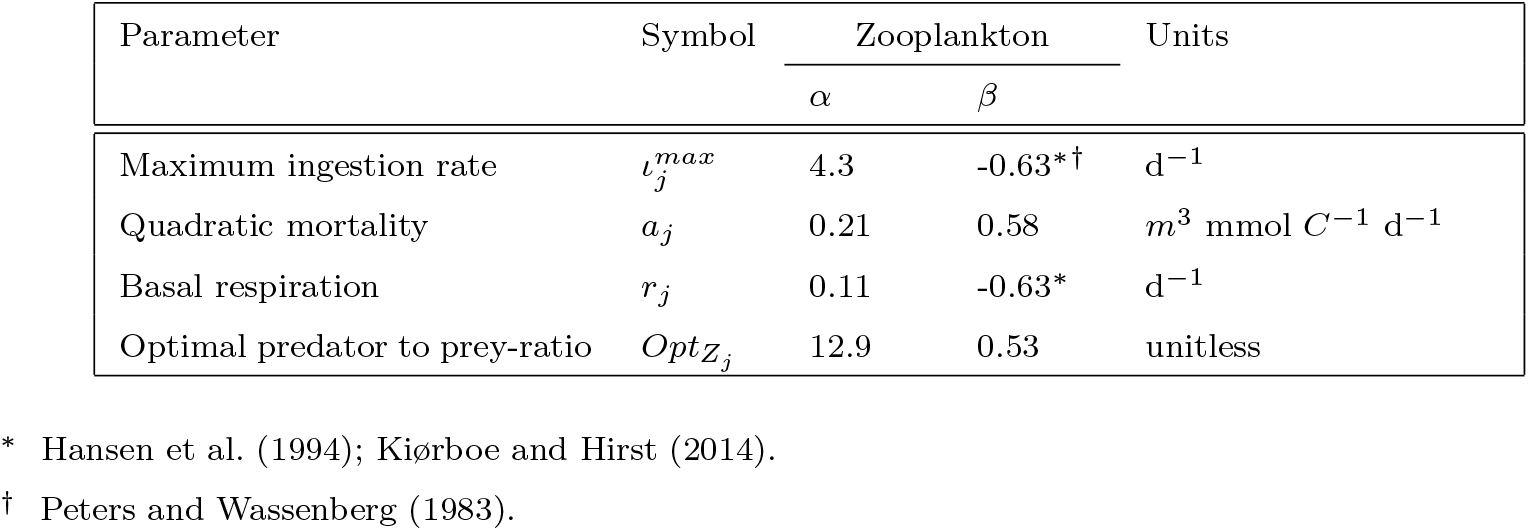
Size-dependent zooplankton biological parameters and scaling coefficients (t = *aV^β^*).

Phytoplankton growth (mmol C m^−3^ d^−1^) is determined by a carbonspecific, light-saturated photosynthesis rate 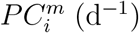 for each phytoplankton group, modulated by a non-dimensional factor which reflects sensitivities to light 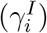:

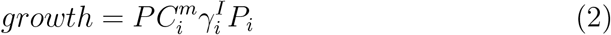

The light sensitivity of growth rate 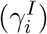 is parameterized using a modified form of the Geider et al. (1997, 1998) dynamic growth model (Eq. 3), where 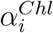 (mmol C m^2^ (mg Chl W d)^−1^) is the Chl-specific initial slope of the photosynthesis irradiance curve, *I* (W m^−2^) is the instantaneous irradiance, and 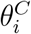 is the *Chl*:C ratio (mg Chl mmol C^−1^), as follows:

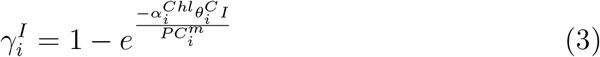

MARBL also uses a multi-column subgrid scale treatment for light, following Long et al. (2015), which reduces biases when light fields are heterogeneous, such as high latitude spring bloom conditions. The above equation describes the biomass-specific rate of photosynthesis as a saturating function of irradiance. 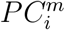 is expressed as a function of the reference carbon-specific photosynthesis rate 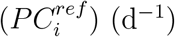 (the maximum achievable carbon-specific photosynthesis rate at the reference temperature) for each phytoplankton group, the temperature dependence function 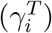, and the nutrient limitation function 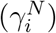 for each phytoplankton type. 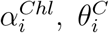 and 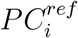 are set according to allometric relationships defined by Edwards et al. (2015b), Geider et al. (1997) and (Marañón et al., 2013) explained in more detail in Section 2.2.

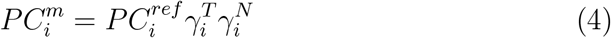

Nutrient limitation of growth 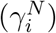 is determined by the most limiting nutrient resource (mmol m^−3^) for that phytoplankton, computed using Liebig’s Law of the Minimum:

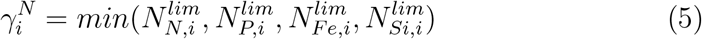

where the nutrients considered are nitrogen, iron, silicate, and phosphorous, yet not all nutrients are required for each phytoplankton group. Diatoms require nitrogen, phosphorous, silicate, and iron. Picoplankton, the mixed phytoplankton group and diazotrophs do not assimilate silicate, and diazotrophs are not limited by nitrogen due to their nitrogen fixing abilities. Simultaneous limitation by multiple nitrogen forms, i.e., nitrate (NO_3_) and ammonium (NH_4_), is represented following the substitutable model of O’Neill et al. (1989); see Long et al. (2021) for more details. A similar approach is used to compute limitation terms for phosphate (PO_4_) and semiliable dissolved organic phosphate (DOP). The effect on growth rate of each of these nutrients for each phytoplankton is represented according to Michaelis-Menten kinetics:

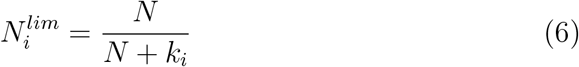

where, *k_i_* (mmol N m^−3^) represents the half-saturation nutrient concentration for each phytoplankton type *i* set according to allometric relationships defined by Edwards et al. (2012) explained in more detail in Section 2.2.4.

In contrast to the default MARBL configuration, which uses the Eppley (Eppley, 1972) temperature scaling with the *Q*_10_ factor, here, the temperature modulation of growth for each phytoplankton 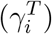 is represented by the Arrhenius-Van’t Hoff equation (Arrhenius, 1915). Kremer et al. (2017) found that the Arrhenius-Van’t Hoff temperature scaling function more closely matched observations of how phytoplankton growth rates scale with temperature. Here, the temperature modulation of phytoplankton rates is expressed relative to the metabolic rate at a reference temperature.

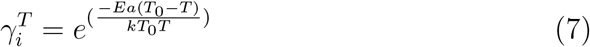

where, *Ea* is the activation energy (eV), *k* is the Boltzmann’s constant (*k* = 8.617 × 10^−5^ eV K^−1^), *T* is temperature (°K), and *T*_0_ represents the reference temperature in the model (293.15°K). *Ea* for all phytoplankton is set to 0.32 eV (Kremer et al., 2017), except for picoplankton, where *Ea^pp^* is set to 0.42 eV, a value derived from an analysis of the Kremer et al. (2017) dataset. Multiple studies have shown that picoplankton have a higher temperature sensitivity compared to phytoplankton of larger sizes (Chen et al., 2014; Stawiarski et al., 2016; Anderson et al., 2021), and model experimentation showed that higher *Ea* is key for excluding picoplankton from polar regions, compared to lower temperature sensitivity of larger sizes.

Predation on phytoplankton (predation_*p*_; mmol C m^−3^ d^−1^) is modeled using a Holling type II function, where predation pressure increases approximately linearly as prey increases, before saturating to a maximum rate at high prey concentrations:

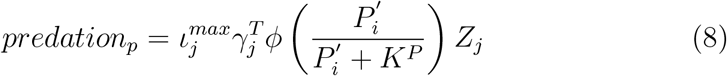

Here, 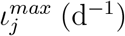 is the zooplankton maximum ingestion rate at a reference temperature, and scales with zooplankton size (Section 2.3.1). The temperature modulation of ingestion for each zooplankton 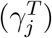 is similar to that of phytoplankton, but differs by having a greater zooplankton activation energy (*Ea^Z^*) compared to autotrophs (Allen et al., 2005), as Rubisco carboxylation (rate limiting for photosynthesis) has a lower *Ea* than ATP synthesis (Allen et al., 2005; López-Urrutia et al., 2006). Thus, for zooplankton, *Ea^Z^* are set to 0.55 eV, a value similar to López-Urrutia et al. (2006) observations. This is in contrast with the default version of MARBL, which uses the same temperature sensitivity for both phytoplankton and zooplankton processes. Among global ocean biogeochemical models, very few models use a higher temperature sensitivity for zooplankton vs. phytoplankton (e.g., PISCES; Aumont et al., 2015); the majority of models use either the same scaling for all plankton (e.g., COBALT; Stock et al., 2014b, 2020), or no temperature scaling of zooplankton rates (e.g., MEDUSA; Yool et al., 2013). Using a higher temperature sensitivity in zooplankton vs. phytoplankton may have implications for phytoplankton-zooplankton coupling and trophic transfer, particularly under climate change, however, a systematic study has not yet been done.

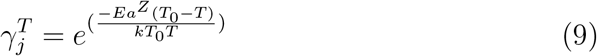

*K^P^* (mmol C m^−3^) is the half-saturation prey concentration which regulates ingestion efficiency at low prey concentrations, and is set as a constant value for all zooplankton (Section 2.3.1). *ϕ* (unitless) is the feeding preference coefficient, which describes the probability of a given predator ingesting prey of a particular size. The feeding preference coefficient will be discussed in greater detail in Section 2.3.2. 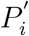 is the phytoplankton concentration in excess of a temperature-and depth-dependent refuge, and is used to limit autotroph mortality at low biomass (mmol C m^−3^) (Long et al., 2021).

Phytoplankton loss (loss_*p*_; mmol C m^−3^ d^−1^) is represented by a linear loss term (*m_i_*) (d^−1^) that includes non-predation mortality and a collection of density-independent processes such as dissolved organic matter (DOM) exudation, viral lysis, and cell death. In MARBL-SPECTRA, instead of a single allometric scaling, linear mortality is set as a fraction of 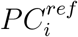, with a factor of 0.02 for diatoms and 0.03 for all other phytoplankton (see Section 2.4).

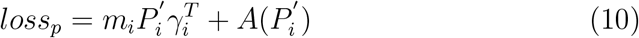

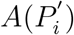 (mmol C m^−3^ d^−1^) represents loss of phytoplankton due to aggregation and unresolved predation, and this loss goes directly to particulate organic matter (POC).

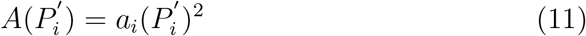

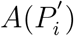 is parameterized by a “quadratic mortality” rate, *a_i_* (d^−1^ mmol C^−1^ m^3^) for all phytoplankton that falls between imposed minimum 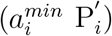 and maximum aggregation 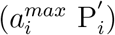 rates.

As with phytoplankton, the time rate of change in zooplankton is a balance between growth and losses to predation and non-predation mortality:

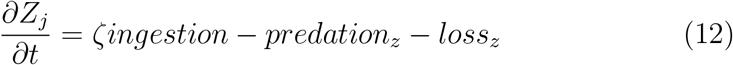

Zooplankton ingestion (mmol C m^−3^ d^−1^) represents the predation gains by zooplankton from their prey, partitioned into gross growth efficiency, res-piration and egestion. Where, *ζ* (*unitless*) represents the maximum gross growth efficiency coefficient (i.e., the maximum fraction of ingestion that goes to growth; Straile, 1997), and is set to be 30% for all zooplankton. Zooplankton (*Z_j_*) are able to feed on both phytoplankton (*P_i_*) and other zooplankton (*Z_k_*, this excludes the largest zooplankton), modulated by a feeding preference coefficient (*ϕ*). Ingestion is thus the total consumption for a zooplankton (*Z_j_*):

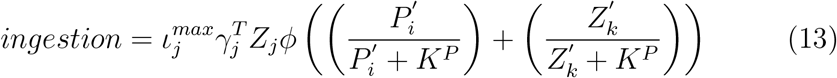

where 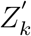 is the zooplankton concentration in excess of a temperature- and depth-dependent threshold, used to limit zooplankton mortality at low biomass (mmol C m^−3^). 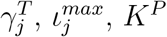, and *ϕ* are described above.

Of the total zooplankton ingestion, 35% is egested, yielding an assimi-lation efficiency (AE) of 65%, which is within the general range of 60-80% used for zooplankton (Carlotti et al., 2000). Partitioning of the egestion into the POC, DOC, and DIC fractions depends on zooplankton size, and is discussed further in Section 2.3.3. Active respiration is 35% of ingestion, with basal respiration a product of the biomass-specific basal respiration rate and zooplankton biomass (mmol C m^−3^ d^−1^). Thus, zooplankton production becomes:

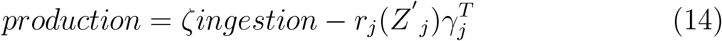

where *r_j_* is the biomass-specific basal respiration rate (d^−1^), and is set following allometric relationships, as discussed in Section 2.3.3 (see also Table 3).

Except for the largest mesozooplankton, all other zooplankton are also predated upon by larger zooplankton. These predator-prey relationships are displayed in Fig. 1. The predation term (predation_*z*_; mmol C m^−3^ d^−1^) thus represents the predation losses from one zooplankton (*Z_k_*) to another (*Z_j_*):

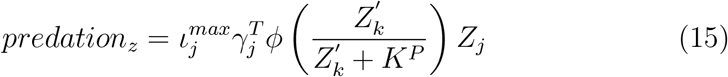

Zooplankton losses (loss_*z*_; mmol C m^−3^ d^−1^) consist of a linear loss term representing zooplankton basal respiration, as well as unresolved losses to higher tropic levels (Steele and Henderson, 1992), which are represented by a biomass- and temperature-dependent quadratic mortality term a*_j_* (*m*^3^ mmol *C*^−1^ d^−1^). The largest mesozooplankton size class has a higher quadratic loss mortality to compensate for higher trophic grazing not directly represented by grazing from the modeled ecosystem. Total non-predation losses include basal respiration and quadratic losses:

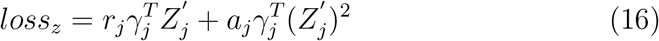

### 2.2. Allometric scaling of phytoplankton traits

Many phytoplankton traits, such as metabolic rate and nutrient affinity, are related to cell size (Chisholm, 1992; Litchman et al., 2007; Edwards et al., 2012). There are also ecologically meaningful differences in key traits across phytoplankton functional groups. For example, large diatoms tend to grow more slowly than do smaller diatoms, but diatoms as a whole tend to grow more rapidly than other competing functional groups such as dinoflagellates (Litchman et al., 2007; Edwards et al., 2012). We use key trade-offs among functional traits to model community composition of marine phytoplankton along environmental gradients. For example, major functional traits in phytoplankton parameters such as nutrient-dependent growth and uptake have physiological trade-offs in the ability to acquire and utilize resources (Litchman et al., 2007). Incorporating these traits and trait trade-offs into a model allows it to represent the fundamental and realized ecological niche of a species and facilitates its representation across a range of environmental and biotic conditions (Ward et al., 2012; Follows et al., 2007). This approach has been used in a range of plankton community and biogeochemical models (e.g. Fuchs and Franks, 2010; Ward et al., 2012; Taniguchi et al., 2014; Dutkiewicz et al., 2015a; Heneghan et al., 2020; Stock et al., 2020). MARBL-SPECTRA adopts this approach, and ties organismal traits and interactions to body size and functional group by employing allometric rules to distinguish within plankton groups instead of individually tuning each plankton functional type. The use of these allometric relationships substantially reduces the number of free parameters.

The effect of size variation on phytoplankton traits is often idealized using a series of power-law scaling function with the typical form:

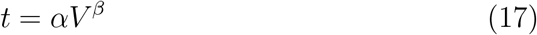

where *t* is the physiological trait, *V* is the cell volume across plankton in the model, *α* is a scaling constant, and *β* is an exponent describing the size dependence. Below, we describe important allometric traits in the model, and discuss how our choices of *α* and *β* across functional groups were informed by empirical studies across many phytoplankton sizes and functional groups (Litchman et al., 2007; Edwards et al., 2012; Marañón et al., 2013). Model traits are summarized in Figs. 2 & 3 and Tables 1 & 2.

**Figure 2:**
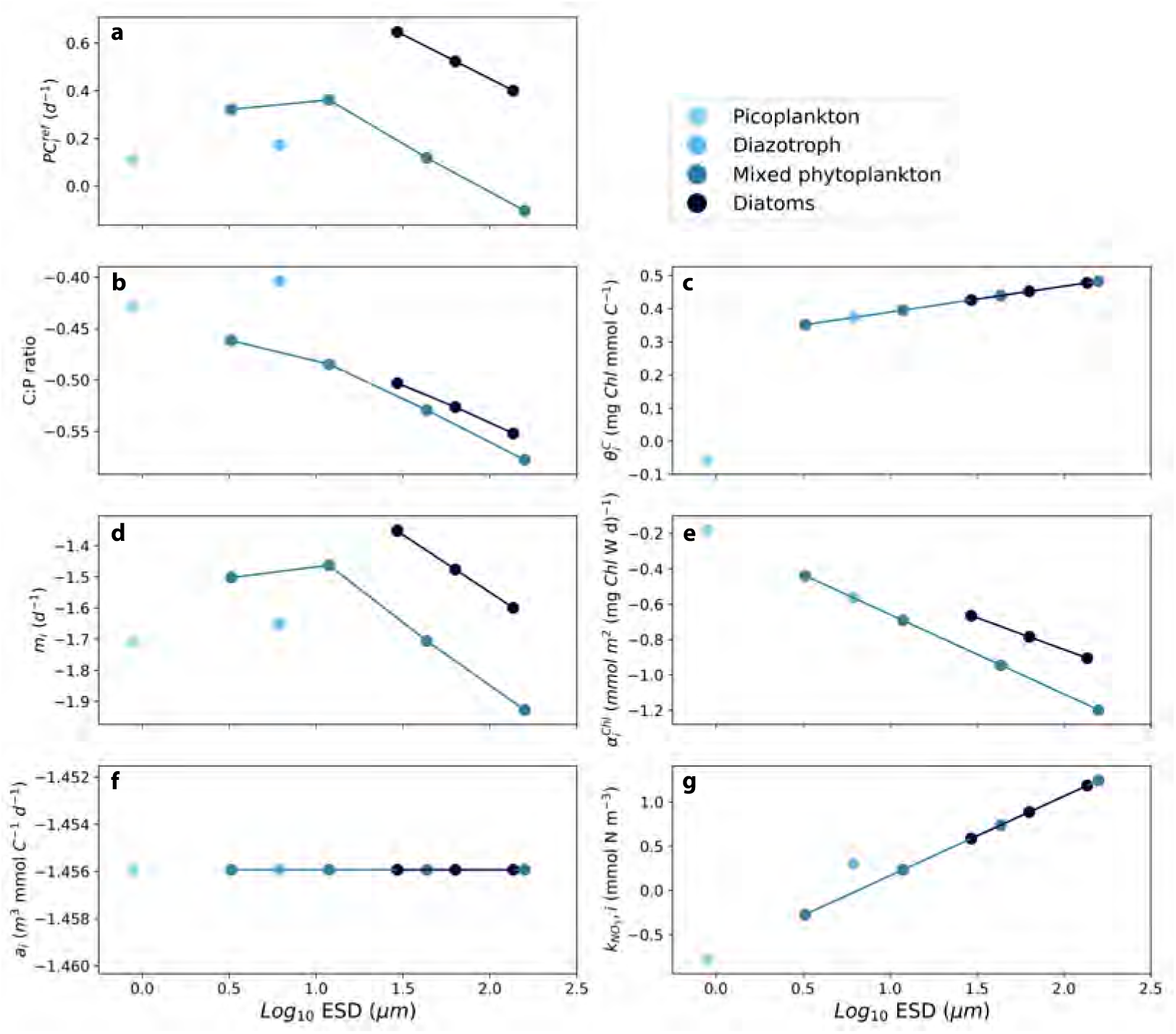
Model phytoplankton traits and parameters. Relationships for picoplankton (light blue), diazotrophs (sky blue), mixed phytoplankton (medium blue), and diatoms (dark blue), between equivalent spherical diameter (ESD) and (a) daily C-specific rate of photosynthesis 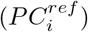 at a reference temperature (20°C), (b) P:C ratio, (c) Linear mortality (m_*i*_), (d) aggregation loss ((*a_i_*) representing aggregation and unresolved predation, (e) maximum value of the Chl to phytoplankton carbon ratio 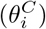, (f) initial slope of photosynthesis-irradiance curve 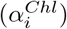, and half saturation nitrate concentration (*k*_*NO*_3__).

**Figure 3:**
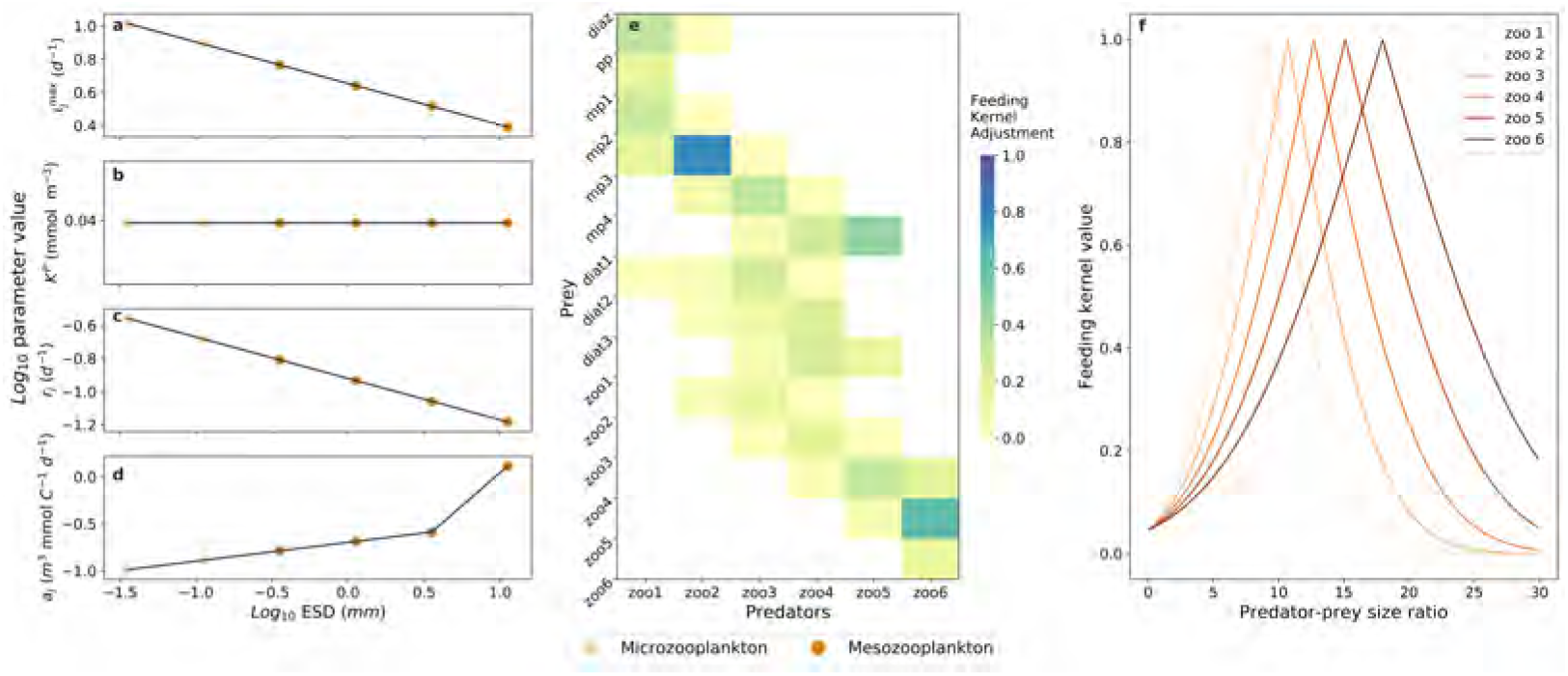
Zooplankton parameters. Relationships for microzooplankton (<0.2 mm ESD) (light orange) and mesozooplankton (>0.2 mm ESD) (dark orange) between equivalent spherical diameter (ESD) and a) maximum zooplankton ingestion rate 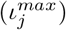, b) zooplankton grazing half saturation constant for grazing (*K^P^*), c) basal respiration (*r_j_*), and (d) quadratic mortality (*a_j_*) (representing predation by higher trophic levels). e) Maximum grazing rates between predator and prey pairs and f) Value of the feeding kernel, which is then modified to give the feeding preference coefficient. The mean and width of the feeding kernel increases as zooplankton sizes increase.

#### 2.2.1. Phytoplankton growth and photosynthesis

Within phytoplankton functional types and for cells larger than approximately 5 *μm* ESD, phytoplankton 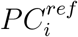 rates generally decrease with increasing cell size (Marañón et al., 2013; Edwards et al., 2012; Tang, 1995). For phytoplankton smaller than 5 *μm* ESD, larger cells grow faster than do smaller ones, such that the overall relationship between 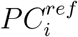 and cell size is unimodal, with the fastest growth rates achieved for cells around 5 *μm* ESD (Marañón et al., 2013; Edwards et al., 2012; López-Sandoval et al., 2014). In addition, different functional groups tend to deviate from this overall pattern. Diatoms, for example, tend to grow faster than other groups (Edwards et al., 2012).

Consistent with this overall paradigm, 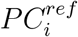 rates in MARBL-SPECTRA scale with cell volume and functional group with a scaling slope of −0.14 (Marañón et al., 2013) within functional types (Fig. 2a). Diatoms have higher 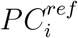 rates than other groups, but within diatoms, 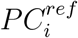 decreases with body size ranging from 4.4 *d^−1^* for the smallest diatoms, and 2.5 *d^−1^* for the largest diatom, consistent with observations from Marañón et al. (2013) and López-Sandoval et al. (2014). The high 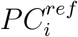 rates of diatoms facilitate their high biomass in nutrient-rich habitats and during bloom conditions (Margalef, 1978). Picoplankton have a low 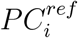 rate of 1.3 *d*^−1^ compared to quickly growing but somewhat larger cells. We have found that incorporating lower 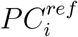 rates for picoplankton and mixed phytoplankton smaller than 5 *μ*m ESD was essential for controlling small phytoplankton growth. Otherwise, the picoplankton and smallest mixed phytoplankton dominate, particularly in the higher latitude seasonal seas. 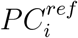 rates for the second smallest mixed phytoplankton through the largest mixed phytoplankton ranged between 2.3 *d*^−1^ and 0.8 *d*^−1^. Diazotrophs have 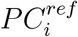 rates of 1.5 *d*^−1^, roughly half compared with other phytoplankton of their size, due to the high energetic demands of nitrogen fixation, which reduces growth rates (Margalef, 1978; Fu et al., 2005; Falcón et al., 2005; Breitbarth et al., 2008).

#### 2.2.2. Chlorophyll-carbon ratios

The chlorophyll to carbon ratio 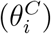 reflects photoacclimation and nutrient stress and has been shown to track phytoplankton physiology both in the laboratory and in the field (Behrenfeld et al., 2005; Behrenfeld and Boss, 2003). Under the dynamic growth parameterization (Geider et al., 1997), the carbon-specific photosynthesis rate is a function of irradiance as well as 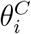. Chl synthesis is regulated by the balance between light absorption and photosynthetic carbon fixation (Geider et al., 1998). Depending on this ratio, a fraction of newly assimilated nitrogen is diverted to the synthesis of Chl. 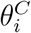 values vary greatly among species and are affected nonlinearly by ambient nutrients, light, and temperature (Geider et al., 1997; Behrenfeld et al., 2002). 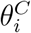 is maximal at high temperatures and low irradiances under nutrient-replete conditions and declines at high irradiances, especially at low temperature and under nutrient limiting conditions (Geider et al., 1997). The maximum chlorophyll to carbon ratio 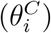 is used as an input parameter in the model but is weakly constrained by empirical studies, with generally higher 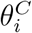 values for large diatoms and lower values for picoplankton such as *Prochlorococcus* (Geider et al., 1997; Sathyendranath et al., 2009). We therefore used a single allometric scaling relationship for most of the phytoplankton, where 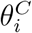 ranges from 0.025 − 0.035 [mg Chl mg *C*^−1^], except for picoplankton which have a 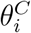 of 0.01 [mg Chl mg *C*^−1^] (Fig. 2c), to match with observed values (Hartmann et al., 2014; Li et al., 2010; Geider et al., 1998, 1986).

#### 2.2.3. Initial slope of the photosynthesis-irradiance curve

Phytoplankton growth rates generally increase under increasing light up to an irradiance optima, at which point growth rates peak before declining due to photoinhibition at higher irradiance levels (Falkowski et al., 1985). These patterns can be illustrated by the photosynthesis-irradiance (P-I) curve, described by the initial slope of the P-I curve 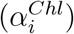 and the biomass-specific rate of photosynthesis 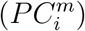 under optimal irradiance (Eq. 3). Variations in 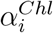 across phytoplankton can be explained in part by consistent differences between major taxonomic groups (Richardson et al., 1983; Cullen and MacIntyre, 1998; MacIntyre, 1998; Boyd et al., 2010) as well as cells of differing size (Geider et al., 1986; Finkel, 2001). Where, a decrease in 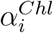 with cell size represents the ability of smaller cells to outperform larger cells under low-light conditions (Edwards et al., 2015b). This is consistent with self-shading of intercellular photosynthetic pigments, also referred to as the “Package effect” (Kirk, 1976), where as cell size increases, the same concentration of pigment, cellular volume, or unit of biomass will adsorb less light due to self-shading of pigment molecules (Kirk, 1994).

Discrepancies across functional types exist, with higher 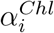 in diatoms compared to other phytoplankton of similar sizes, due to the ability of diatoms to perform relatively well under both limiting light and excessive light (Richardson et al., 1983), or fluctuating light (Litchman, 1998) environments. Based upon the dataset of Edwards et al. (2015b), we set *α*, and *β* to be 0.67 mmol C m^2^ (mg Chl W d)^−1^ and −0.12 for diatoms. For other groups, we used 0.56 mmol C m^2^ (mg Chl W d)^−1^ and −0.15 respectively (Table 2 & Fig. 2e).

#### 2.2.4. Nutrient acquisition

Phytoplankton growth in MARBL is a multiplicative factor of temperature, light, and nutrient limitation, with the nutrient limitation defined using Liebig’s law of the minimum (Moore et al., 2004; Long et al., 2021). Experimental data and theoretical evidence demonstrates that smaller cells have higher rates of nutrient uptake per unit biomass and lower half-saturation constants (Eppley et al., 1969; Aksnes and Egge, 1991) compared to larger cells. The observed *β* between *k* and cell volume falls between 0.24 and 0.45 for NO_3_, and 0.29 to 0.56 for PO_4_ (Edwards et al., 2012). Since our model includes multiple limiting nutrients, we used a single size-scaling exponent of 0.3 for all nutrients. This means that within groups, picoplankton have more efficient nutrient uptake (smallest *k_N_*) compared to the large diatoms and large mixed phytoplankton. Diazotrophs (e.g. *Trichodesmium* spp.) are the only exception from this allometric scaling, since they are less efficient at inorganic nutrient uptake (McCarthy and Carpenter, 1979) and they often occur as large colonies, where their surface to volume considerations imply higher half-saturation constants relative to the small phytoplankton and diatom groups (Letelier and Karl, 1998). However, higher half saturation constants for diazotrophs were only set for nitrogen and iron. See Table 2 for all nutrient half-saturation constants and scaling coefficients.

The C:N:P elemental stoichiometry of phytoplankton is highly variable across ocean biomes, but exhibits general patterns of dependence on temperature, nutrients, and species composition (Finkel et al., 2010; Martiny et al., 2013; DeVries and Deutsch, 2014; Moreno and Martiny, 2018). While increased temperatures and nutrient limitation, particularly phosphorus limitation, can increase C:P and N:P levels (Thingstad, 1998; Yvon-Durocher et al., 2015), there still exists broad stoichiometric characteristics due to taxonomy and cell size (Finkel et al., 2010). For example, picoplankton such as *Prochlorococcus* and *Synechococcus* generally exhibit higher C:P and N:P ratios than other phytoplankton, even in P-replete conditions (Bertilsson et al., 2003). In contrast, diatoms and other large-sized phytoplankton that typically dominate the high latitudes exhibit low C:P and N:P ratios (Martiny et al., 2013).

Here, we impose C:P stoichiometric ratios based on phytoplankton size (Finkel et al., 2010), such that the smallest mixed phytoplankton (mp1) has a C:P ratio of 147:1 and the largest mixed phytoplankton (mp4) has a C:P ratio of 54:1. Diazotrophs have a C:P of 300:1 and picoplankton have a C:P of 215:1 (Bertilsson et al., 2003; White et al., 2006). Imposing fixed, non-Redfield C:P stoichiometry enables the elemental ratios within MARBL-SPECTRA to respond to changes in plankton community composition, though it does not include mechanisms such as frugality, where the frugal plankton living in phosphorus-deficient environments uses less phosphorus than its counterpart in phosphorus-replete environments (Martiny et al., 2013; Galbraith and Martiny, 2015). While a dynamic model of C:P stoichiometry based on the frugality hypothesis is provided within MARBL (Galbraith and Martiny, 2015; Long et al., 2021), enabling it with MARBL-SPECTRA would have added 15 additional tracers to the model, making computation extremely expensive.

Photoadaptation is calculated as a variable phytoplankton ratio of chlorophyll to nitrogen based on Geider et al. (1998). The model allows for variable Fe/C and Si/C ratios with an optimum and minimum value prescribed. As ambient Fe (or Si for diatoms) concentrations decline the phytoplankton lower their cellular quotas. Phytoplankton N/P ratios are fixed at the Redfield value of 16, but the diazotroph group has a higher N/P atomic ratio of 50 (see a detailed description of the model in Moore et al. (2001, 2004)). Thus, community N/P uptake varies with the phytoplankton community composition.

### 2.3. Zooplankton allometric scaling terms

#### 2.3.1. Zooplankton Ingestion

The vital rates of organisms depend on their size: ingestion, metabolism, and growth rates all increase with body size to a power of approximately 0.75, typically such that the mass-specific rates decline with body mass to a power of near −0.25 (Peters and Wassenberg, 1983; Kiørboe and Hirst, 2014; Hansen et al., 1997). In MARBL-SPECTRA, zooplankton are defined as heterotrophs that can consume phytoplankton, other zooplankton, or a combination of both. Zooplankton ingestion rates are calculated as a function of prey carbon concentrations using the Hollings type II (Michaelis-Menten) function. There are two free parameters, maximum ingestion rate 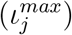 and the half-saturation constant for grazing (*K^P^*).

Size-based variations in maximum specific ingestion rates were calculated as an allometric function of the predator biomass, with biomass-specific rates decreasing as biomass increases (Hansen et al., 1997). 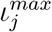 are also modified by the feeding preference coefficient, which is a function of the ratio between the predator size and the prey size. *K^P^* are highly variable and typically hard to constrain. Across all the zooplankton classes, *K^P^* has been found to be independent of body size (Hansen et al., 1997). Therefore, the effective *K^P^* is set to 1.1 (mmol C m^−3^) across zooplankton types (Table 3).

#### 2.3.2. Predator-prey relationships

In addition to physiological rates, predation in marine ecosystems is size-specific, with larger prey eating a characteristic size range of smaller prey (Sheldon et al., 1977; Hansen et al., 1994; Cohen et al., 1993; Barnes et al., 2008). We model these trophic links using a feeding kernel (FK) that is further modified to give the feeding preference coefficient (Eq. 8, *ϕ*). Feeding kernels constitute the probability of a given predator ingesting prey of a particular size and can be highly variable, reflecting a great deal of measurement uncertainty and biological variability, with various studies employing gaussian, Laplace, and log-normal distributions (Law et al., 2009; Fuchs and Franks, 2010; Taniguchi et al., 2014; Heneghan et al., 2020). Here, we use the feeding kernel as a starting point to determine predator-prey feeding relationships, which are subject to additional tuning to achieve plankton distributions consistent with large-scale observational constraints. The feeding kernel *FK_Z_j__* is represented as a complementary error function:

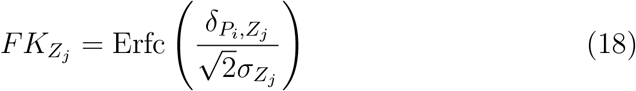

where Erfc(*x*) = 1 - Erf(*x*), the standard error function:

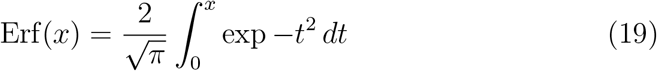

and is closely related to the cumulative distribution function of the standard normal distribution.

Here, the numerator of the Erfc function is *δ_P_i_,Z_j__*, which is the absolute value of the difference between the predator-prey size ratio and the optimal predator-prey size ratio for any given predator, *Z_j_*:

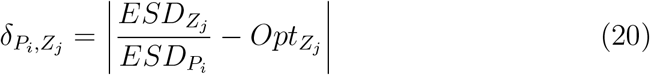

where *ESD_Z_j__* and *ESD_P_i__* refer to the ESD (in mm) of the predator *Z_j_* and its prey, respectively, and *Opt_Z_j__* is the predator specific optimal predator-to-prey size ratio. Note that we have used the *P_i_* subscript here for simplicity, but the prey of *Z_j_* encompasses both phytoplankton and zooplankton prey. *Opt_Z_j__* varies from approximately 7.5 to 18 from the smallest to the largest zooplankton, and represents the phenomenon that the mean predator-to-prey size ratio will often increase as predator size increases (Hansen et al., 1994). The parameters that define *Opt_Z_j__* are given in Table 3.

Similarly, the width of the feeding kernel also increases as predator size increases, reflecting both the wider organism size ranges, varied prey capture strategies, and multiphagy of larger zooplankton (Hansen et al., 1994; Kiørboe, 2011). Here it is represented by *σ_Z_j__*, which is in the denominator of Eq. 18, and is defined as *σ_Z_j__* = 0.5 * *Opt_Z_j__*.

The feeding kernel as defined by the complementary error function has the property of being exactly 1 when *δ_P_i_,Z_j__*. =0, and then declines in a sigmoidal manner as the predator-to-prey ratio increasingly differs from the optimum. Put together over the whole range of predator-to-prey size ratios, the resultant curve increases to a point at the center (the optimum), and declines on either side (see Fig. 3f), resembling the Laplace (double exponential) distribution, which was used in Fuchs and Franks (2010). It is important to note here that the exact shape of the feeding kernel is secondary relative to the following adjustments to the predator-prey feeding preference coefficient, as they allowed us to tune this highly sensitive but poorly constrained grazing term to achieve plankton distribution patterns consistent with large-scale observational constraints.

Building upon this basic kernel formulation, we made several adjustments to model predator prey interactions to improve the representation of the model plankton community. First, we increased microzooplankton grazing on picoplankton relative to the value in the feeding kernel. The increased grazing accounts for unresolved grazing by heterotrophic nanoflagellates, and allows for a higher stability in picoplankton populations. Second, we increased grazing on small diatoms, kept medium diatoms the same, and decreased the grazing on large diatoms. The increase in grazing on small diatoms was necessary to provide a strong top-down control on the abundance of fast growing small diatoms. The reduced grazing on larger diatoms accounts for their ability to form colonies and/or frustules to reduce losses to predation (Oostende et al., 2018). Third, zooplankton production rates were lower than estimated values (Landry and Calbet, 2004), so we decreased zooplankton grazing on other zooplankton to increase zooplankton production. Fourth, we also increased grazing on the small implicit calcifying mixed phytoplankton group to increase zooplankton production and at the same time reduce their high abundance in subpolar regions. Fifth, to increase mesozooplankton production, we decreased microzooplankton grazing on small diatoms, to allow mesozooplankton to take advantage of diatom blooms especially in very productive regions of the ocean.

To ensure ingestion does not exceed maximum ingestion for a particular predator, the feeding kernel values were normalized by predator, such that the sum of all feeding kernel values per predator equalled one. The individual feeding kernel values per predator-prey pair then modified predator ingestion rates (Eq. 8).

#### 2.3.3. Zooplankton egestion, metabolism, and mortality

In MARBL-SPECTRA, 65% of the ingested prey carbon is assimilated with the remainder (35%) egested via sloppy feeding and fecal pellet production. Egestion can be partitioned into sinking particulate organic carbon (POC), semi-labile dissolved organic carbon (DOC), and highly labile DOC (which is instantly transformed to dissolved inorganic carbon, or DIC). To simulate the effect of zooplankton size on sinking fecal pellet carbon (Stamieszkin et al., 2015), the fraction that goes into the POC pools (*ρ^POC^*) differs by zooplankton size and phytoplankton prey type, ranging from 21% of egestion from small microzooplankton (zoo1) predation to 100% of egestion from the largest mesozooplankton (zoo6) predation. Small microzooplankton grazing on picoplankton contributes 0% to POC, and for zooplankton grazing on diatoms, the fraction of egestion to POC increases by 20%. All detritus remineralization length scales are from the CESM 2.0 configuration (and do not vary by zooplankton type; reproduced in Table S1), with longer length scales for ballast materials (silicate, biogenic calcium carbonate, and mineral dust), which simulates faster sinking speeds (Armstrong et al., 2002). For more information on detritus remineralization, see Long et al. (2021). Finally, the routing of egestion to DOC (*ρ^DOC^*) vs. DIC (*ρ^DIC^*) is split from the remainder of the non-POC fraction: 25% goes to DOC and 75% to DIC. Since MARBL-SPECTRA does not have an explicit heterotrophic bacteria and does not simulate labile DOC generated by the microbial loop (Azam et al., 1983), this 75% fraction refers to the fast remineralization (hours to days) of labile DOC from sloppy feeding.

One of the new additions to the zooplankton formulation in MARBL-SPECTRA is the separation between zooplankton ingestion-based (active) respiration and biomass-based (basal) respiration. Active respiration is a fixed fraction of total ingestion (35%), but basal respiration is a function of zooplankton biomass and decreases with size (Kiørboe and Hirst, 2014). Similar to the specific ingestion rate (Section 2.3.1), the biomass-specific basal respiration (*r_j_*) is temperature-dependent and decreases with body size with a *β* of −0.25 (Hansen et al., 1997; Kiørboe and Hirst, 2014) and *α* of 0.12 d^−1^. These scaling relationships were converted to be scaled with volume with a *β* of −0.63 and *α* of 0.11 d^−1^.

Zooplankton mortality due to predation by other zooplankton is resolved in the model in the lower size classes, but it becomes progressively unresolved with increasing size classes. To account for the unresolved predation by higher trophic levels (fish, carnivorous jellies, marine mammals), the zooplankton quadratic mortality (*a_j_*) increases with biovolume with a *β* of 0.21 and *α* of 0.58 m^3^ mmol C–1 d^−1^. We increased the quadratic mortality for largest mesozooplankton by a factor of four to account for higher level grazing (Section 2.4). The fraction of zooplankton quadratic mortality fluxing into particulate and dissolved organic matter pools depends on diet and organisms size (see Long et al. (2021) for more details). With a greater proportion of large zooplankton mortality being transferred to particulate organic matter pools compared to smaller zooplankton.

### 2.4. Model Calibration

Many of the parameters required to simulate planktonic foods webs are difficult to measure directly, yet are highly important to simulate carbon and energy flow patterns (Stock and Dunne, 2010). In order to produce a balanced ecosystem, two main calibrations were done. First, the zooplankton loss terms (basal respiration and quadratic mortality) were calibrated to preserve global totals of zooplankton production while largely maintaining allometric trait relationships across size classes (Kiørboe and Hirst, 2014; Hansen et al., 1997). We increased the zooplankton quadratic mortality for the largest mesozooplankton by a factor of four to account for unresolved predation by higher trophic levels. Phytoplankton linear mortality and aggregation loss were also calibrated because these parameters are poorly constrained by observations. Instead of a single allometric scaling, linear mortality was set as a fraction of 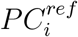, with a factor of 0.02 for diatoms and 0.03 for all other phytoplankton. The lower linear mortality for diatoms provide a slight advantage over other phytoplankton, particularly in nutrient rich (upwelling and polar) regions and an increase in their global production.

Second, the grazing half-saturation prey concentration for zooplankton were calibrated to allow higher global total zooplankton production. These parameters are poorly constrained by observations (Hansen et al., 1997), but the values used (Table 3) still fall within the observed ranges in Hansen et al. (1997). Because grazing half-saturation constants have been shown to be independent of body size (Hansen et al., 1997), only one K^P^ had to be calibrated, because it was used for every zooplankton.

### 2.5. Sensitivity Analysis

We conducted model sensitivity analyses in order to understand the impact on model ecological outcomes of changes in key model parameters in MARBL-SPECTRA (Figs. S11–S20). We carefully examined changes to key allometric scaling exponents for both phytoplankton (i.e., nutrient uptake, light affinity) and zooplankton (ingestion and mortality rates), as well as a size-independent scaling of the grazing half-saturation constant. All parameter sensitivity runs were conducted following the same physical forcing as the MARBL-SPECTRA base simulation, and were run for 20 years to stabilize the ecosystem. We compared the biomass of small and large phytoplankton and microzooplankton and mesozooplankton in the last 10 years of these model sensitivity simulations against the same variables in the base MARBL-SPECTRA run. We conducted the sensitivity analyses for these five key parameters, and discuss them in more detail in Section 3.10:

1. The allometric scaling exponent for the initial slope of the photosynthesis-irradiance curve 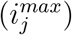 was adjusted to vary ± 25% compared to the scaling in the base run (Fig. S11). These trait variations fall within estimates from Edwards et al. (2015b).
2. The allometric scaling exponent for all nutrient half-saturation constants (*k_N_*) was adjusted to showcase the upper (0.45) and lower (0.24) limits found in Edwards et al. (2012) (Fig. S13).
3. The allometric scaling exponent for zooplankton basal respiration (*r_j_*) was adjusted to vary ± 50% compared to the scaling in the base run (Fig. S15). These variations fall within estimates from Hansen et al. (1997).
4. The allometric scaling exponent for the maximum ingestion rate for zooplankton 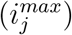 was adjusted to vary ± 50% compared to the scaling in the base run (Fig. S17). These variations fall within estimates from Hansen et al. (1997).
5. The size independent scaling for the grazing half-saturation constant (*K^P^*) was adjusted to vary ± 50% compared to the scaling in the base run (Fig. S19). These variations fall within estimates from Hansen et al. (1997).

### 2.6. Physical Framework

MARBL-SPECTRA builds from the default MARBL configuration in CESM2.1 (MARBL-CESM2.1) in terms of biogeochemistry, plankton interaction and transmission of light as described by tables and equations in Long et al. (2021). However, we have increased the number of plankton functional types and size classes to include greater diversity. Here we briefly provide an overview of MARBL-SPECTRA, and some more detailed description of the more complex ecosystem. More details and the full set of equations and parameters can be found in Long et al. (2021). MARBL runs within the ocean component of the Community Earth System Model version 2 (CESM 2.1) (Moore et al., 2013b; Gent et al., 2011), which is the Parallel Ocean Program, version 2 (Smith et al., 2010). The physical configuration used here is very similar to that in CESM1, and a detailed description and evaluation of the ocean general circulation model in previous versions of CESM is given by Danabasoglu et al. (2012). The model has a nominal horizontal resolution of 1°, with 60 vertical depth levels ranging in thickness from 10 m in the upper 150m to 250 m in the deep ocean (Moore et al., 2013b). The sea-ice component (CICE) is described by Hunke et al. (2017).

MARBL-SPECTRA simulates 55 tracers, including 17 non-living tracers and 38 tracers associated with the plankton community. This includes 27 tracers associated with the nine phytoplankton size classes, with each phytoplankton C, Chl, and Fe tracked separated (Moore et al., 2001, 2004). In addition, there are three phytoplankton Si tracers associated with the three diatom classes, as well as two phytoplankton CaCO_3_ tracers associated with the two implicit calcifiers that are part of the mixed phytoplankton classes. Constant stoichiometry was assumed for zooplankton, therefore only six zooplankton carbon tracers were included. The model simulates six dissolved organic matter pools, including semi-labile and refractory dissolved organic carbon, nitrogen, and phosphorus (Letscher and Moore, 2015; Letscher et al., 2015). It also includes sinking particulate pools and an explicit simulation of the biogeochemical cycling of key elements (C, N, P, Fe, Si, O, plus alkalinity) (Moore et al., 2004). Riverine nutrients (N, P, Si, Fe), dissolved inorganic carbon, alkalinity, and DOM fluxes are supplied to the CESM2 ocean model via the nutrient loading estimates from GlobalNEWS (Mayorga et al., 2010) and the Integrated Model to Assess the Global Environmental-Global Nutrient Model (IMAGE-GNM) (Beusen et al., 2015, 2016). The plankton community component is coupled with a carbonate chemistry module based on the Ocean Carbon Model Intercomparison Project (OCMIP)(Najjar et al., 1999), allowing dynamic computation of surface ocean pCO_2_ and air-sea CO_2_ flux.

MARBL-SPECTRA simulations are forced with the Common Ocean-Ice Reference Experiment (CORE-II) data set (Large and Yeager, 2009). The forcing period from 1948 to 2009 (62 years) underwent two repeating cycles. This differs from CORE-II protocol where forcing undergoes five repeating cycles (Griffies et al., 2009). A shorter integration does not provide a fully-equilibrated model solution in the deep ocean, but has been used for studying surface ocean dynamics (Stock et al., 2014b). Thus, by the end of the 62 year spin up time, surface biomass distributions are nearing an equilibrium state, even if deep ocean tracers may not be. We focus our analyses on the final 20 years of the simulation (1990-2009). Code for generating the namelist parameters for MARBL-SPECTRA are available at: https://github.com/marbl-ecosys/spectra-config.

## 3. Results

### 3.1. Biogeochemical comparisons

MARBL-SPECTRA qualitatively captures the large-scale global biogeochemical and ecological patterns evident in available observations. Simulated global annual mean marine NPP, and nitrogen fixation, averaged over the top 150 m, and sinking POC flux below 150m are shown in comparison to empirical estimates in Table 4. For these metrics, the model falls within range of empirical estimates. The global total marine diatom production was 14 Pg C yr^−1^, about 28% of total NPP and falls within estimated values (20-40% of total NPP) (Nelson et al., 1995; Aumont et al., 2003). Global total zooplankton production was 12 Pg C yr^−1^ (23% of NPP), falling within empirical estimates (21-25% of NPP) (Landry and Calbet, 2004), with microzooplankton contributing 8.7 Pg C yr^−1^ to overall zooplankton production, and the remainder 3.3 Pg C yr^−1^ coming from mesozooplankton production.

**Table 4:** Global annual averages of marine net primary production (NPP), and nitrogen fixation averaged over the top 150m and sinking POC flux below 150m.

| Biogeochemical field | Global average | Empirical estimate | Reference |
| --- | --- | --- | --- |
| NPP | 52 Pg C yr<sup>-1</sup> | 35-70 Pg C yr<sup>-1</sup> | (Carr et al., 2006) |
| Nitrogen Fixation | 107 Tg N yr<sup>-1</sup> | 51 - 196 Tg N yr<sup>-1</sup> | (Luo et al., 2014; Tang et al., 2019; Wang et al., 2019; Großkopf et al., 2012) |
| POC flux | 6.8 Pg C yr<sup>-1</sup> | 4-12 Pg C yr<sup>-1</sup> | (Dunne et al., 2007; DeVries and Weber, 2017) |

The model captures the large-scale surface (top 10 m) NO_3_ (Fig. 4a-c), PO_4_ (Fig. 4d-f), and SiO_3_ (Fig. 4g-i) distributions, with low nutrient concentrations in the subtropical gyres and higher nutrient concentrations in subpolar and upwelling regions. Compared with the 2018 World Ocean Atlas (WOA18) macronutrient observations (Garcia et al., 2019) (Fig. 4b,e & h), near-surface NO_3_, PO_4_, and SiO_3_ concentrations were slightly different in the model, with a bias of +0.50, +0.090, and −1.4 mmol m^−3^ respectively. For NO_3_ and PO_4_, the bias was greatest in the tropical Pacific Ocean. This could be due to a slightly lower export flux from the upper oceans, due to higher nutrient recycling in this region coming from the dominance of smaller phytoplankton (Fig. 8). SiO_3_ biases were highest in the Southern Ocean, potentially due to insufficiently vigorous diatom production depressing SiO_3_ consumption. In the subpolar North Pacific Ocean, the model showed lower NO_3_, PO_4_, and SiO_3_, compared to the WOA18 observations (Fig. 4 c, f, i). This underestimation of macronutrients in the North Pacific is likely due to insufficient vertical mixing in this region, with phytoplankton production utilizing surface nutrients faster than they can be replenished. Simultaneously, overproduction of diatoms, due to insufficient *Fe* limitation stimulates the utilization of nutrients, leading to under-representation of the high nitrate, low chlorophyll (HNLC) region of the sub-Arctic North Pacific.

**Figure 4:**
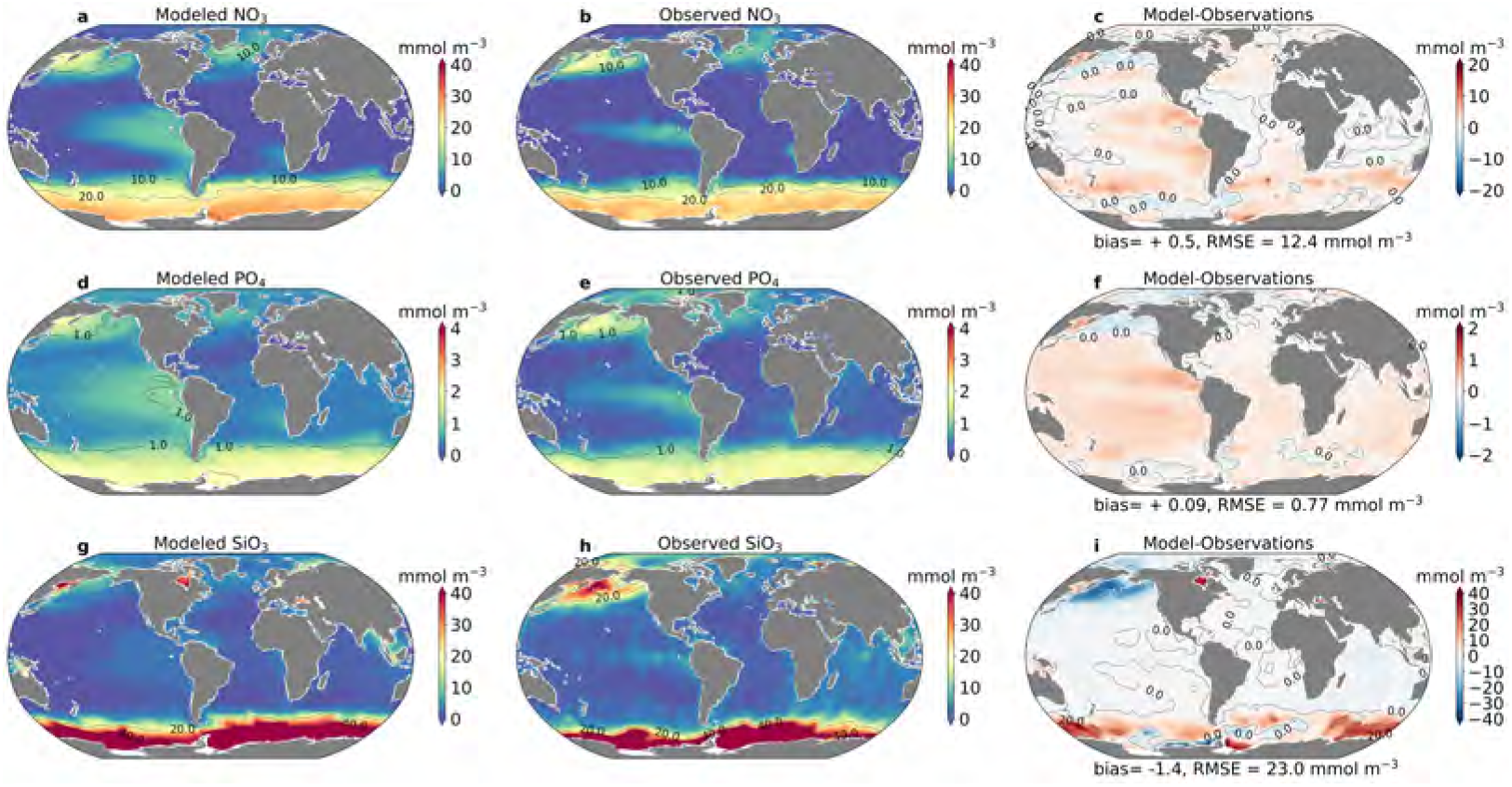
Macronutrients. Annual average modeled (a,d,g) and observed (b,e,h) surface (top 5m) concentrations of NO_3_, PO_4_ and SiO_3_ and their differences (Model-Observations; c,f,i). Observations are from the 2018 World Ocean Atlas release. (Garcia et al., 2019)

### 3.2. Limitation of model phytoplankton growth

Using biomass-weighted means of nutrient limitation, we show the nutrients most limiting growth for each phytoplankton in the model (Fig. 5). Phytoplankton growth was limited primarily by the availability of nitrate (NO_3_) or Fe and regionally by PO_4_ (diazotrophs) and SiO_3_ (diatoms) (Fig. 5), consistent with previous modeling studies using CESM (Moore et al., 2013a; Long et al., 2021). The degree of growth limitation by nutrients becomes stronger as body size increases (Fig. 5). This occurs because smaller phytoplankton have a greater capacity to acquire nutrients via diffusion relative to their nutrient demands (Edwards et al., 2012). Nutrient replete areas (white areas in Fig. 5) were characterized by where the concentration of nutrients was high enough to support growth rates *>* 90% of the maximum potential growth rate. For picoplankton, these occurred in the equatorial upwelling region and the subpolar and polar regions (Fig. 5a). For diazotrophs, these occurred in equatorial regions of the Atlantic and Indian Oceans, as well as the western Pacific (Fig. 5b). Diatoms and mixed phytoplankton were rarely nutrient replete due to their high nutrient requirements (Fig. 5c,d). In addition to these overall patterns, diazotrophs underwent stronger PO_4_ limitation in the North Atlantic due to enhanced N_2_ fixation simulated by Fe associated with dust deposition (Wu et al., 2000).

**Figure 5:**
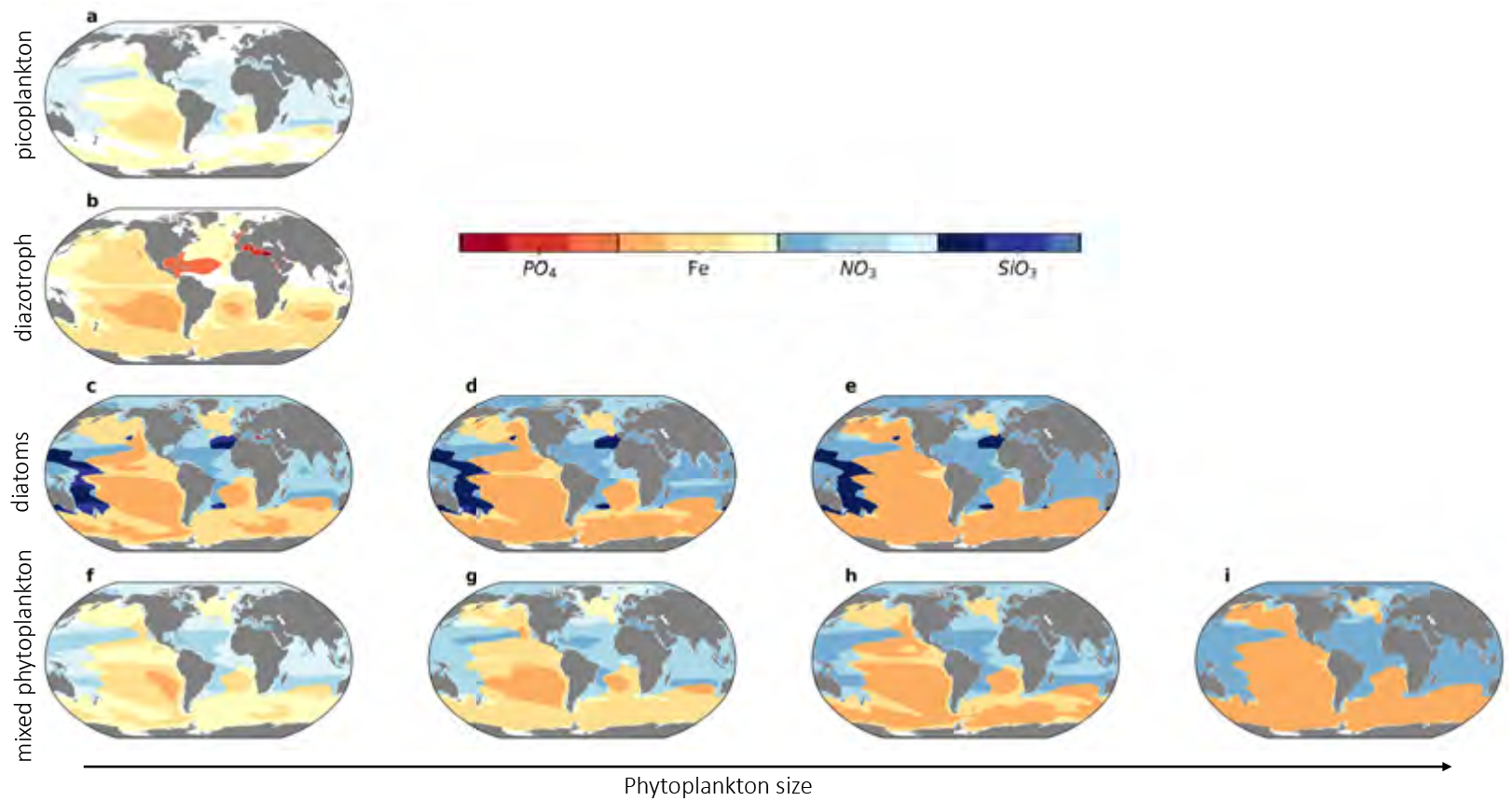
Phytoplankton nutrient limitation in top 100m. Figure shows the most limiting nutrient and strength of limitation, as calculated by a biomass-weighted minimum of the nutrients needed for growth for each phytoplankton type. For example, diatoms are the only group that are limited by SiO_3_. The darker shading within each limitation term corresponds to stronger limitation (lower 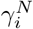 values, see Eq. 5). Areas where nutrients are replete (defined as 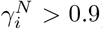) are marked in white. Rows categorize phytoplankton by their functional group: (a) picoplankton, (b) diazotroph, (c) diatoms, and (d) mixed phytoplankton. Columns indicate relative phytoplankton size within each group, increasing from left to right.

### 3.3. Chlorophyll

Model annual-mean surface (top 10 m) chlorophyll exhibits plausible spatial gradients tied to provision of nutrients to the ocean surface and good overall agreement with observations (Fig. 6). Surface chlorophyll observations were obtained from the Sea-viewing Wide Field-of-view Sensor (SeaWiFS) climatology from 1998-2009, which corresponds to the last twelve years of the CORE-II forcing dataset (Large and Yeager, 2009). Model chlorophyll was low in subtropical gyres due to wind-driven downwelling and low surface nutrient availability (Fig. 4a). In higher latitudes and upwelling areas, higher nutrient concentrations allowed for higher chlorophyll concentrations (Fig. 6a). Model annual average chlorophyll generally exceeds observations in subtropical and temperate locations, while the model underestimates chlorophyll in the Arctic, Antarctic, and coastal upwelling regions (Fig. 6c).

**Figure 6:**
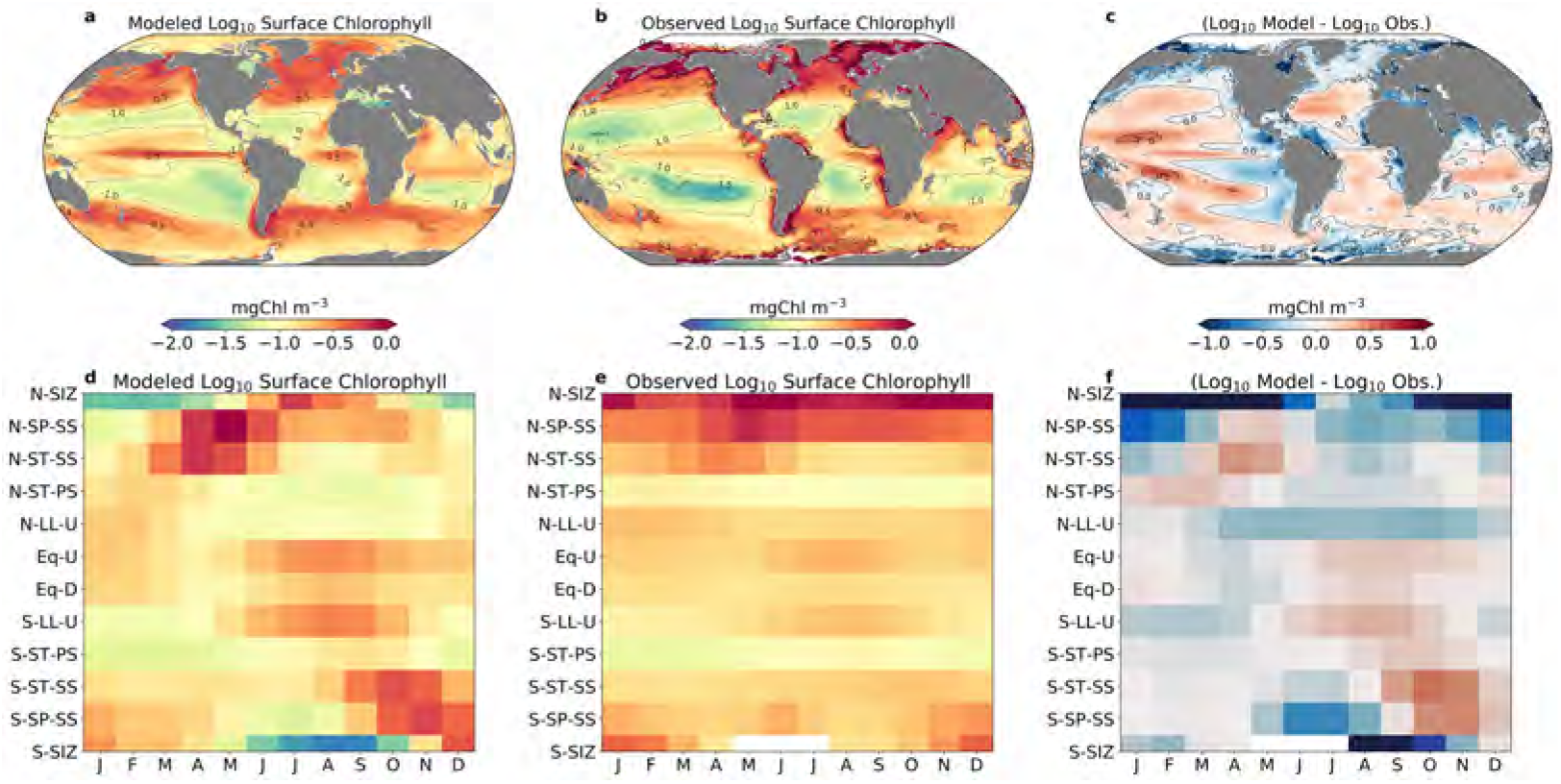
Surface (top 10 m) chlorophyll concentration (mg Chl m^−3^). (a) Simulated annual-mean surface chlorophyll, (b) satellite-derived (SeaWiFS) estimate of annual-mean surface chlorophyll, (c) model – SeaWiFS, (d) mean monthly modeled surface chlorophyll by biomes; (e) mean monthly SeaWiFS surface chlorophyll by biomes; and (f) difference between model and observations on a monthly, per-biome basis. Biomes were designated using Sarmiento et al. (2004) (Fig. S1)

The positive chlorophyll bias in subtropical and subpolar seasonally stratified biomes of the northern hemisphere was do to an earlier than observed phytoplankton bloom that started in March and ended around June, about a month earlier than the observed bloom in April through June (Fig. 6f). This was perhaps due to lower mesozooplankton biomass in the spring from lack of diaupasing zooplankton, yielding insufficient top-down control on phytoplankton and leading to an earlier spring bloom (Behrenfeld, 2014). The higher chlorophyll concentrations in the central Equatorial Pacific between July to November (Fig. 6f) were associated with higher nutrient delivery to the surface from the Equatorial upwelling zone. In the subtropical and subpolar Southern Hemisphere, a stronger bloom initiated sooner (September) than observational estimates (November/December) (Fig. 6f), leading to a positive bias in subtropical and subpolar seasonally stratified biomes of the Southern Hemisphere.

Vertical profiles of model chlorophyll show important biases compared with observations. Comparing vertical chlorophyll profiles of The Bermuda Atlantic Time-series Study (BATS), and the Hawaii Ocean Time-series (HOT) stations (Fig. S2), MARBL-SPECTRA simulated a shallower deep chlorophyll maximum (DCM) layer for BATS (60-80m) compared to HOT (70-90m). However, compared with observed values, MARBL-SPECTRA simulated DCMs that were too shallow for both of these regions. For instance, for the period of comparison (1990-2009) annual averages from HOT and BATS indicated a DCM layer falling between 80-110m for BATS (http://batsftp.bios.edu/BATS/bottle/bats and 70-130m for the HOT station (https://hahana.soest.hawaii.edu/FTP/hot/primary_production/). The tendency of these deep chlorophyll maximum to be shallower than observations may be due do a variety of reasons, such as the lack of representation of low-light adapted ecotypes of picoplankton which are generally restricted to the deep euphotic zone (Moore et al., 2002; Johnson et al., 2006; Moore and Chisholm, 1999) contributing to the deep chlorophyll maximum. The under-representation of mixotrophy in the model could also contribute to this bias, as it has been found that the incorporation of mixotrophy in models has helped represent DCMs more accurately (Moeller et al., 2019).

### 3.4. Phytoplankton biogeography

The distribution of small, medium, and large phytoplankton in the model is consistent with satellite-derived size class estimates from Hirata et al. (2011). The small group included the picoplankton (pp) and the smallest mixed phytoplankton (mp1), the medium group included the smallest diatom (diat1), the diazotroph (diaz), and the medium mixed phytoplankton (mp2), and the large group included the largest two diatoms (diat2, diat3) and the largest two mixed phytoplankton (mp3, mp4). The satellite algorithm used by Hirata et al. (2011) estimated the biomass of three phytoplankton size classes as microphytoplankton (>20 *μm*), nanophytoplankton (2-20 *μm*) and picophytoplankton (<2 *μm*).

Small phytoplankton dominated the subtropical gyres with over 70% of the total Chl_*a*_ (Fig. 7c). These regions are characterized by strong vertical stratification and weak nutrient delivery to the surface. Here, small phytoplankton can outcompete larger phytoplankton due to their higher scaled nutrient and light affinities allowing them to maintain positive net growth at low nutrient concentrations compared to larger competitors (Edwards et al., 2012). Medium phytoplankton dominated in subpolar gyres, coastal upwelling zones, and equatorial upwelling regions where nutrient delivery is greater. In these regions, grazing pressure on small phytoplankton prevented small cells from consuming all resources and allowed the medium phytoplankton to become established. The largest phytoplankton were found mainly in polar regions in the Arctic and Southern Oceans, where the balance between growth and predation on small and medium phytoplankton, together with lower light affinities, allowed these larger phytoplankton to survive.

**Figure 7:**
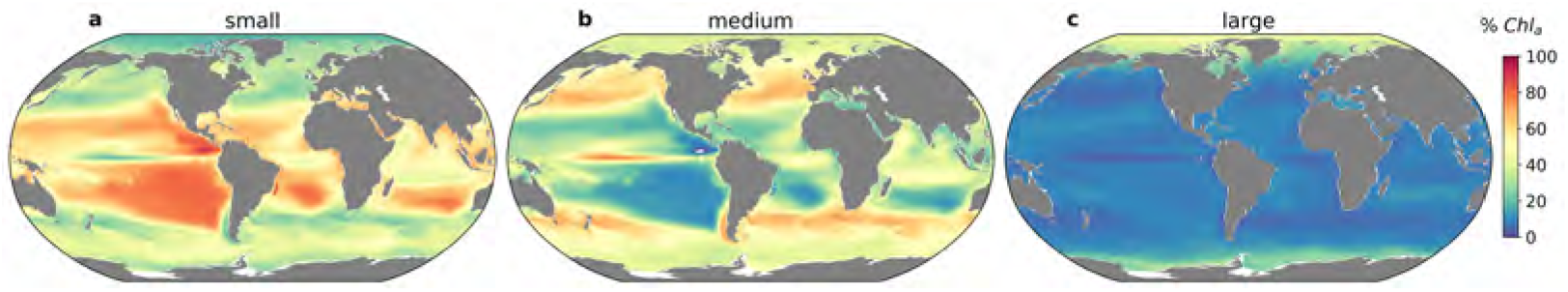
Phytoplankton size class biogeography. Percent of total chlorophyll in each size class: (a) Small phytoplankton (includes picoplankton and smallest mixed phytoplankton, (b) Medium phytoplankton (includes the smallest diatom, the second smallest mixed phytoplankton, and the diazotroph), and (c) Large phytoplankton (includes the two largest mixed phytoplankton, and the largest two diatoms on the model).

Diatoms illustrate the importance of modeling different size classes within each phytoplankton functional type. Diatoms were found from the tropics to the poles, but were most abundant in polar to temperate, nutrient-rich regions, where silicic acid and other nutrients were not limiting. However, the distribution of modeled diatoms varies by size and associated organism traits (Fig. S3g-i). Compared with other diatoms, the smallest diatom has a higher specific growth rate, lower nutrient half-saturation constants, and higher affinity for light (Fig. 2), but also proportionally higher losses to mortality and grazing (Fig. 2; 3). Small diatoms were most abundant in coastal, equatorial upwelling, and subpolar regions (Fig. S3g). Larger diatoms have somewhat lower growth rates, weaker nutrient uptake abilities, and lower light affinity 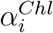, but lower mortality and losses to grazing (Fig. 2; 3). Large diatoms, therefore, were most abundant in subpolar and polar regions (Fig. S3i). The ability to model different size classes within each functional group allowed us to observe these patterns that would otherwise not be resolved.

The phytoplankton size abundance relationship is a general descriptor of phytoplankton community size structure, and plays a fundamental role in pelagic ecosystems as it determines the trophic organization of plankton communities and, hence, the biogeochemical functioning of the ecosystem (Legendre and Rassoulzadegan, 1996; Kiørboe, 1993). The relationship between phytoplankton abundance and cell volume (*V*) follows a power law, *N* = *αV^β^*, where *N* is the cell density and *α* is the intercept of the resulting linear regression. The size-scaling exponent, *β*, is a descriptor of community size structure (Marquet et al., 2005) and generally takes values between −1.4 and −0.5 (Huete-Ortega et al., 2010, 2012; Reul et al., 2005; Marañón et al., 2007; Cermeño and Figueiras, 2008; Cavender-Bares et al., 2001). Here, the slope and intercept of the size abundance relationship was calculated using a linear least-squares regression between the logarithmic abundances of all phytoplankton size classes (depth integrated biomass over cell mass) and their corresponding cell biovolume.

Overall, MARBL-SPECTRA captures the spatial gradients in phytoplankton size structure driven by the provision of nutrients to the ocean surface (Barton et al., 2013). Locations with more negative slopes tend to have relatively few large phytoplankton present, whereas a less negative slope indicates the presence of proportionally more large phytoplankton(Cermeño et al., 2006). In MARBL-SPECTRA (Fig. 8b), the most negative slopes (between −1.2 and −0.9) occurred in the permanently stratified oligotrophic subtropical gyres where small phytoplankton dominated and large cells were scarce (Fig. 7). The highest contribution of small cells was especially seen during the boreal and austral summer of permanently stratified subtropical gyres, and lower latitude upwelling regions (Fig. 8b). The least negative slopes (>-0.9) were found in more productive regions of the ocean, like the subpolar and polar regions where larger phytoplankton had a higher contribution to total phytoplankton biomass (Fig. S3). Seasonally, less negative slopes were found during the boreal and austral Winter of the seasonal ice zone, and the northern seasonally stratified subpolar gyre (Fig. 8b).

**Figure 8:**
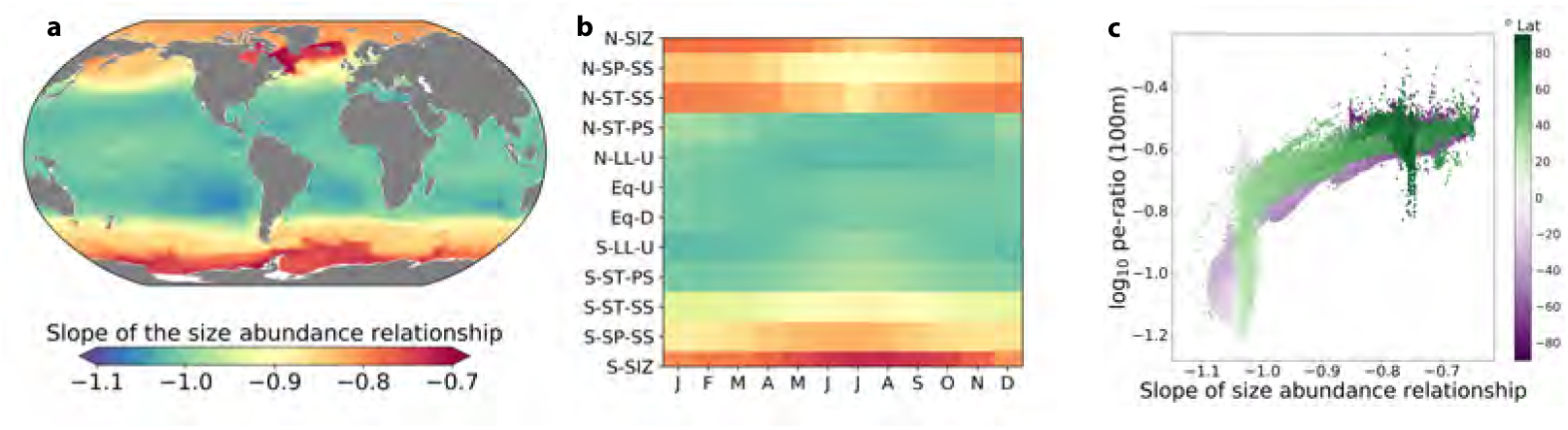
Slope of the size abundance relationship. (a) The annual depth-integrated slope of the size abundance relationship over the top 150m, and (b) mean monthly depth-integrated slope of the size abundance relationship over the top 150m at each biome (Fig. S1). The slope of the size abundance relationship were calculated using a linear leastsquares regression between phytoplankton abundance (depth integrated biomass over cell mass) in all size classes and their corresponding cell biovolume, on a log-log scale. More negative slopes are seen in the stratified waters of low-latitude, open-ocean environments, where small cells account for most of the biomass, and less negative slopes appear in more nutrient-rich, productive regions, where larger cells generally constitute a greater fraction of total biomass than in lower nutrient regions. (c) The relationship between annually averaged log_10_ pe-ratio (defined as the fraction of depth-integrated NPP exported as sinking particles at a 100m) against the depth integrated slope of the size abundance relationship over the top 150m.

### 3.5. Zooplankton production

MARBL-SPECTRA’s global zooplankton production was mostly composed of microzooplankton. Approximately 73% of the total zooplankton production comes from microzooplankton (<200 *μ*m ESD), represented by the two smallest zooplankton groups. These zooplankton dominate the grazing on picoplankton and small mixed phytoplankton. As a result, the mi-crozooplankton are broadly distributed and are most abundant in the oligotrophic and subpolar regions (Fig. S4a,b). MARBL-SPECTRA simulates cross-biome patterns in mesozooplankton biomass, with the highest values in the North Pacific, the equatorial Pacific, and coastal upwelling regions (Fig. 9a).

**Figure 9:**
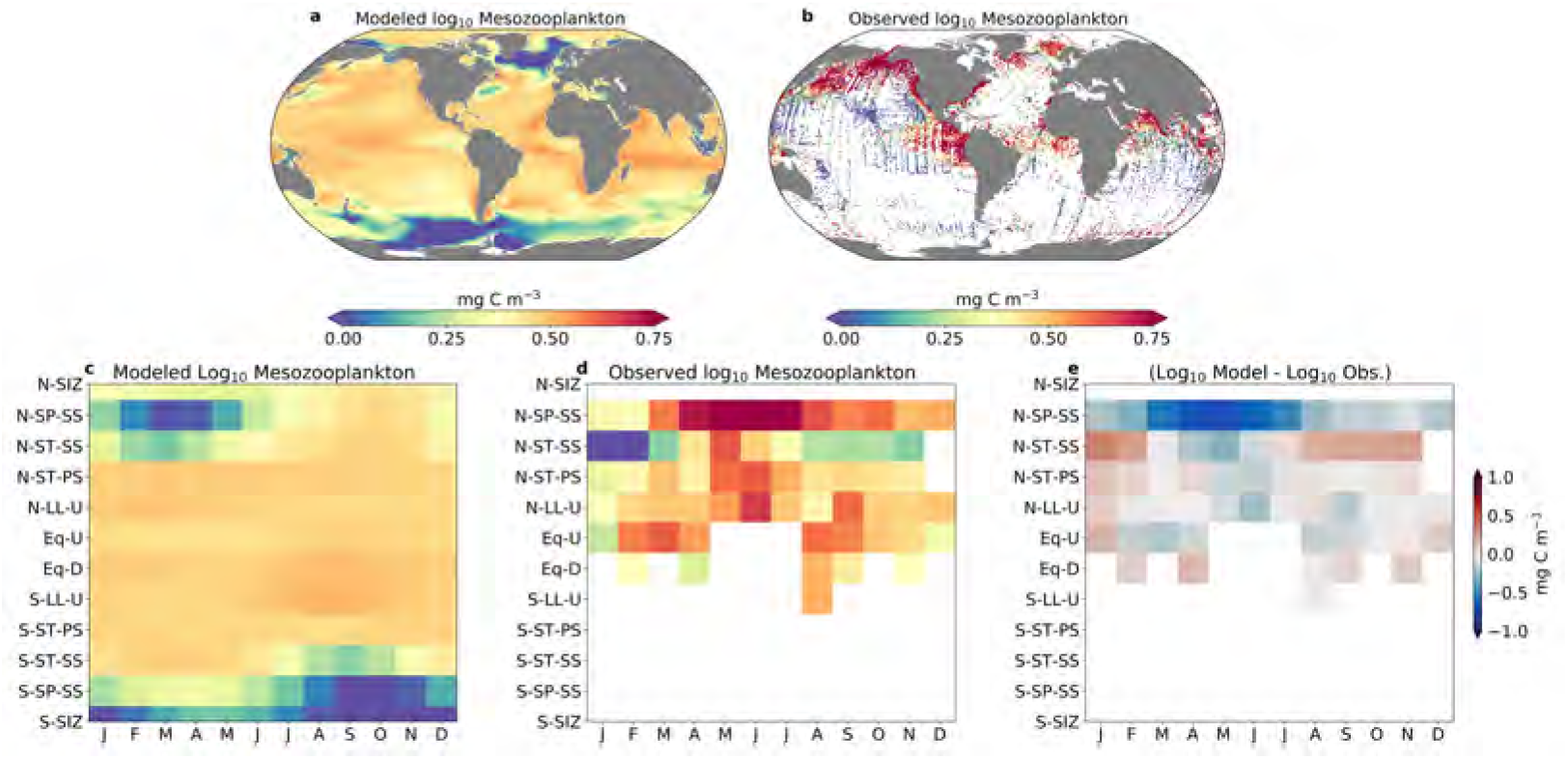
Mesozooplankton biomass. (a) Modeled annual mean mesozooplankton biomass (mg C m^−3^) over the top 150 m (only including largest three mesozooplankton (zoo4 - zoo6), compared with (b) observed annual average mesozooplankton biomass using the COPEPOD database (mg C m^−3^) (Moriarty and O’Brien, 2013). (c) Mean monthly modeled surface mesozooplankton biomass by biomes; (d) mean monthly COPEPOD mesozooplankton biomass by biomes (only showing biomes that have more than 25% of data at each month); and (e) difference between model and observations.

We compared model zooplankton biomass to observations from the NOAA COPEPOD global zooplankton database (https://www.st.nmfs.noaa.gov/copepod/), of which the global mesozooplankton carbon biomass dataset is the most relevant and accessible (Moriarty and O’Brien, 2013) for model output comparison. However, because the COPEPOD database compiles measurements collected by net-tows of epipelagic mesozooplankton captured primarily using large 300 *μ*m nets that under-sample small mesozooplankton (Moriarty and O’Brien, 2013; Landry et al., 2001; O’Brien, 2005)), we only used the three largest mesozooplankton (zoo4-zoo6) for our model-data comparison. Additionally, because the COPEPOD database includes more samples during summer months, we only compared with summer months of each hemisphere (Fig. 9a,b). When comparing modeled and observed mesozooplankton across biomes (Fig. 9c-e), we excluded biomes containing less than 25% of observations at each month. MARBL-SPECTRA’s annual average mesozooplankton biomass (only accounting for grid cells with observations) was 2.7 mg C m^−3^, compared with COPEPOD’s annual average of 4.7 mg C m^−3^. The discrepancy between the model and observations was caused by coastal upwelling regions in the model having lower biomass than observed in the COPEPOD database. Mesozooplankton biomass was lowest in the Southern Ocean and the sub-Arctic North Atlantic (Fig. 9a). Compared with the COPEPOD database observations, MARBL-SPECTRA overestimated mesozooplankton biomass in subtropical regions (Fig. 9e). This can be seen especially in gradients between coastal and offshore regions (e.g. near California Current), where high mesozooplankton production near the coast did not decrease to considerably lower values in oligotrophic regions.

Additionally, we compared mesozooplankton biomass with the Generalized Linear Mixed Model (GLMM) of Heneghan et al. (2020), which is a statistical model product that models mesozooplankton biomass based on a range of point observations, including those from the COPEPOD database (see also Petrik et al., for further details). The comparison with Heneghan et al. (2020) showed a better match in the subtropical gyres and lower latitudes, but the higher latitudes were still biased low in the spring and summer (Fig. S10). This is admittedly one of the largest deficiencies in MARBL-SPECTRA, where we were able to capture the global mean biomass of mesozooplankton but were limited in our ability to simulate their dynamic range, both seasonally and across biomes. We conducted extensive tuning runs to try and improve this dynamical range, but found that the mesozooplankton biomass in MARBL-SPECTRA was stubbornly insensitive to many parameter modifications that were utilized in other models (e.g., basal respiration rate; Stock et al., 2014b). Some of the difficulties in tuning are highlighted in the sensitivity cases, where small modifications in one zooplankton parameter had cascading impacts on the broader food web (Section 3.10 & Figs. S15–S20). This may indicate that the allometrically-constrained zooplankton parameters had insufficient degrees of freedom to simulate the desired zooplankton dynamical range, or that the feeding kernel was too wide. Other discussion of potential model limitations are in Section 4.2.

Model mesozooplankton biomass in MARBL-SPECTRA display a weaker spatial dynamic range compared to observations. The strong negative bias of modeled mesozooplankton (Fig. 9e) in the subpolar and subtropical seasonally stratified biomes of the Northern Hemisphere came from the underestimation of mesozooplankton biomass in the sub-Arctic North Atlantic, along with a three-four month delay in the zooplankton bloom (Fig. 9e). MARBL-SPECTRA does not resolve zooplankton life histories, including dormancy or diapause, which may contribute to these discrepancies. Due to limited observations, we were unable to diagnose seasonal zooplankton biomass patterns in poorly-sampled regions of the ocean (Fig. 9d). However, MARBL-SPECTRA simulated a Southern Hemisphere subpolar zooplankton bloom from December to March and the subtropical seasonally stratified bloom in the Southern Hemisphere from October to June (Fig. 9c). The low latitude upwelling region in the Southern Hemisphere showed a model mesozooplankton bloom from June to October, similar to a shorter one observed in the equatorial upwelling region from June to September.

### 3.6. Generation time

MARBL-SPECTRA simulates plankton generation times and allowed us to analyze their influence by organism size, temperature, and latitude. Plankton generation time was calculated as the ratio of top 150m depth-integrated biomass (mmol m^−2^ C) over production (mmolC m^−2^ d^−1^). Plankton production, as defined in Eq. 14, is the assimilated ingestion minus total respiration (basal and active). We found that the global average generation time for phytoplankton increased with body size, ranging from a few days for picoplankton to a few months for the largest mixed phytoplankton (Fig. 10a). Global average zooplankton generation times ranged from about a week for the smallest microzooplankton to a few months for the largest mesozooplankton. However, there were considerable regional variations in generation time. The longest generation times reached almost a year for the largest mixed phytoplankton and almost two years for mesozooplankton near the poles, with the shortest generation times found in the tropics (Fig. SI, S7, S8). These variations in generation time reflect body size and temperature effects (Gillooly et al., 2001; Gillooly, 2000; Gillooly et al., 2002), although, consumption, respiration, predation, and mortality are also of influence. Generation times for some copepod species have been observed to reach up to 3-4 years (Hirche, 1997). However, MARBL-SPECTRA does not resolve zooplankton life histories such as diapause, which limits generation times for some mesozooplankton especially in polar regions.

**Figure 10:**
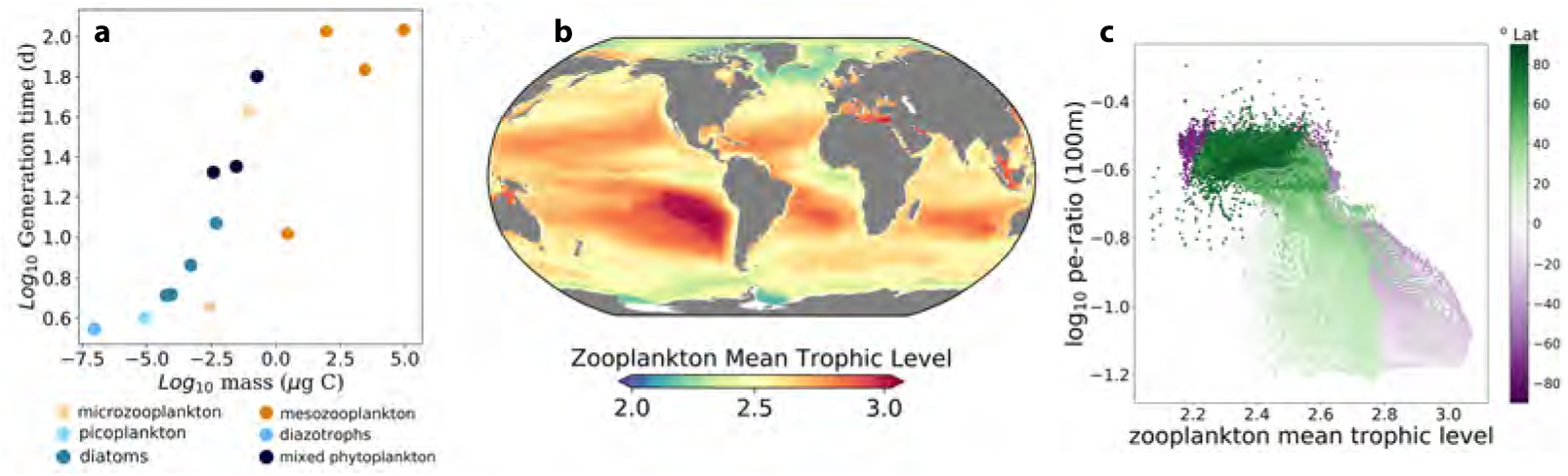
Predator-prey generation time and trophic dynamics. (a) Annual global average generation time averaged over the top 150m in days for each phytoplankton (blue), and zooplankton (orange) size class, as a function of organisms body mass (units). (b) Zooplankton annual mean trophic level over the top 150 m. A trophic level of 2 indicates an entirely herbivorous zooplankton feeding on primary producers. A trophic level 3 indicates secondary consumers, which are carnivorous zooplankton that eat herbivores. (c) Relationship between annually averaged log_10_ pe-ratio at 100m against the zooplankton annual mean trophic level over the top 150m.

### 3.7. Trophic Scaling

The model simulates not only zooplankton biomass and generation time, but also food chain length and zooplankton trophic level. Any zooplankton grazing on phytoplankton were placed in trophic level 2. Because of our complex food web, we calculated each additional trophic level using an ingestion-weighted biomass. For example, if a mesozooplankton size class (e.g., zoo3) obtained half of their food from phytoplankton and half of their food from zooplankton in the 2*^nd^* trophic level, they were placed in trophic level 2.5. Given that we simulated fixed size classes in the zooplankton, the zooplankton mean trophic level was used as a proxy for food chain length.

Our model simulations showed average zooplankton trophic levels to be highest in the oligotrophic subtropical gyres and lowest in polar regions around the Southern Ocean and the Arctic Ocean (Fig. 10b & Fig. S6). We explored the relationship between food-chain length and particulate export efficiency (pe-ratio) (Fig. 10c), defined as the fraction of depth-integrated NPP exported as sinking particles at 100m. We found that higher zooplankton mean trophic levels correlate with lower export production (lower pe-ratio). The higher mean trophic levels occurred primarily in oligotrophic regions of the ocean (Fig. 10b), where there tends to be greater recycling of organic matter in the microbial loop. Lower mean zooplankton trophic levels occurred with higher export production (higher pe-ratio) and were mainly found in coastal and upwelling regions, as well as subpolar and polar regions (Fig. 10b). These regions tend to have proportionally more larger phytoplankton (Fig. 8), which strongly contribute to export production.

### 3.8. Zooplankton to phytoplankton biomass ratio

MARBL-SPECTRA resolved spatial and temporal variations in the biomass pyramid in lower trophic levels of marine ecosystems, and consequently can provide mechanistic insights on factors regulating this structure. Regions of high phytoplankton and zooplankton biomass were concentrated in subpolar and coastal regions, whereas the oligotrophic gyres supported much lower total biomass (Fig. S5 a,b). The zooplankton to phytoplankton biomass ratio was at or below 1 in most of the ocean, consistent with observations from Irigoien et al. (2004); Gasol et al. (1997), and modeling results in marine (Vallina et al., 2014) and lake systems (Yuan and Pollard, 2018). Z:P biomass ratios were also shown to vary seasonally, in this case focusing on data from Northern Hemisphere subpolar and polar regions (35°N-90°N). Here, the highest Z:P ratios occurred in winter months, driven by declines in phytoplankton biomass (Fig. 11c).

**Figure 11:**
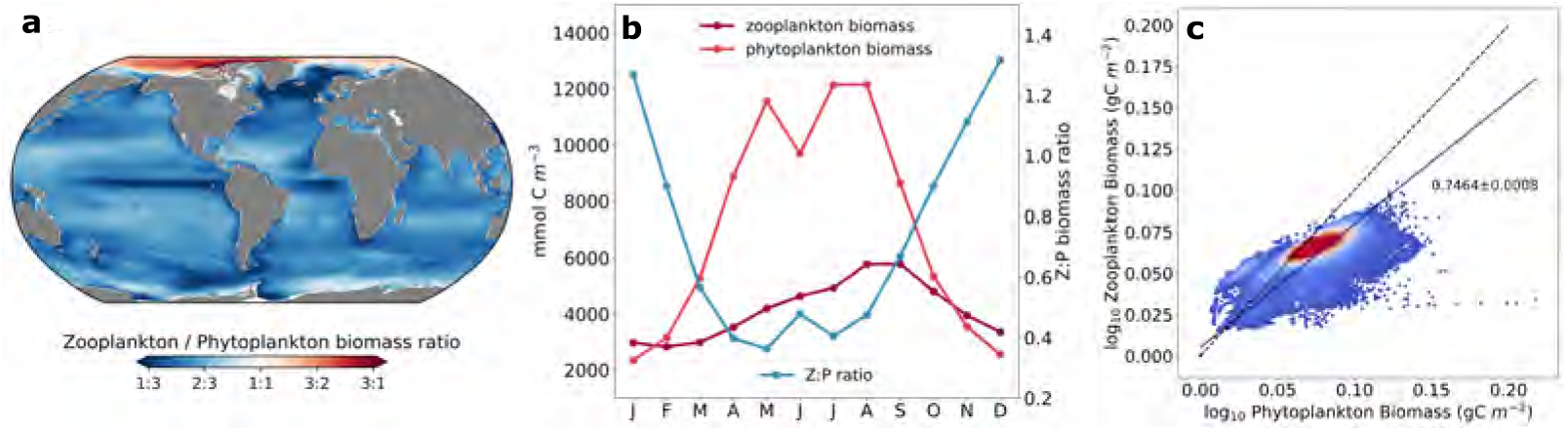
Predator-prey biomass ratios. (a) Global map of the zooplankton to phytoplankton biomass ratio, showing the depth integrated annual mean over the top 150 m. (b) Seasonal zooplankton biomass (dark pink), phytoplankton biomass (light pink), and zooplankton to phytoplankton biomass ratio (blue) of polar and subpolar regions in the Northern Hemisphere (35°N-90°N). (c) log_10_ zooplankton and phytoplankton biomass relationship integrated over the top 150 m. The dashed black line represents the 1:1 line, and the solid black line represents the least squares line of best fit, which has an exponent of 0.7464 ± 0.0008 in bold (with 95% CI in blue).

The model indicates that as phytoplankton biomass increases, so too does zooplankton biomass. However, the rate of increase in zooplankton biomass was less than for phytoplankton biomass, such that the slope of the log_10_-log_10_ relationship between model phytoplankton and zooplankton biomass was approximately 0.75 (Fig. 11c). In regions of low phytoplankton biomass, such as the oligotrophic gyres, Z:P ratios were close to 1:1, suggesting a tight and efficient coupling between small phytoplankton and their microzooplankton consumers. In regions of higher phytoplankton biomass, Z:P ratios were lower, suggesting a greater degree of decoupling between predators and prey.

One exception from this overall relationship was the Arctic Ocean, which had much higher Z:P ratios, in some cases approaching 3:1. Here, the balance between strong seasonal bottom-up (light and temperature) controls and intense grazing pressure in these regions might explain the high Z:P biomass ratios.

### 3.9. Plankton phenology

The enhanced plankton community in MARBL-SPECTRA provides a mechanistic representation of the function and dynamics of plankton phenology, where the phenology of model plankton is tied to their body size, traits, and interactions. Here, we show the seasonal cycle of biomass for seven 5° by 5° locations in the global ocean (Fig. 12). While the details of a given site may differ between the model and observations for a range of reasons, the model simulates a seasonal succession at each location tied to nutrient delivery, temperature, light availability, and grazing pressure.

**Figure 12:**
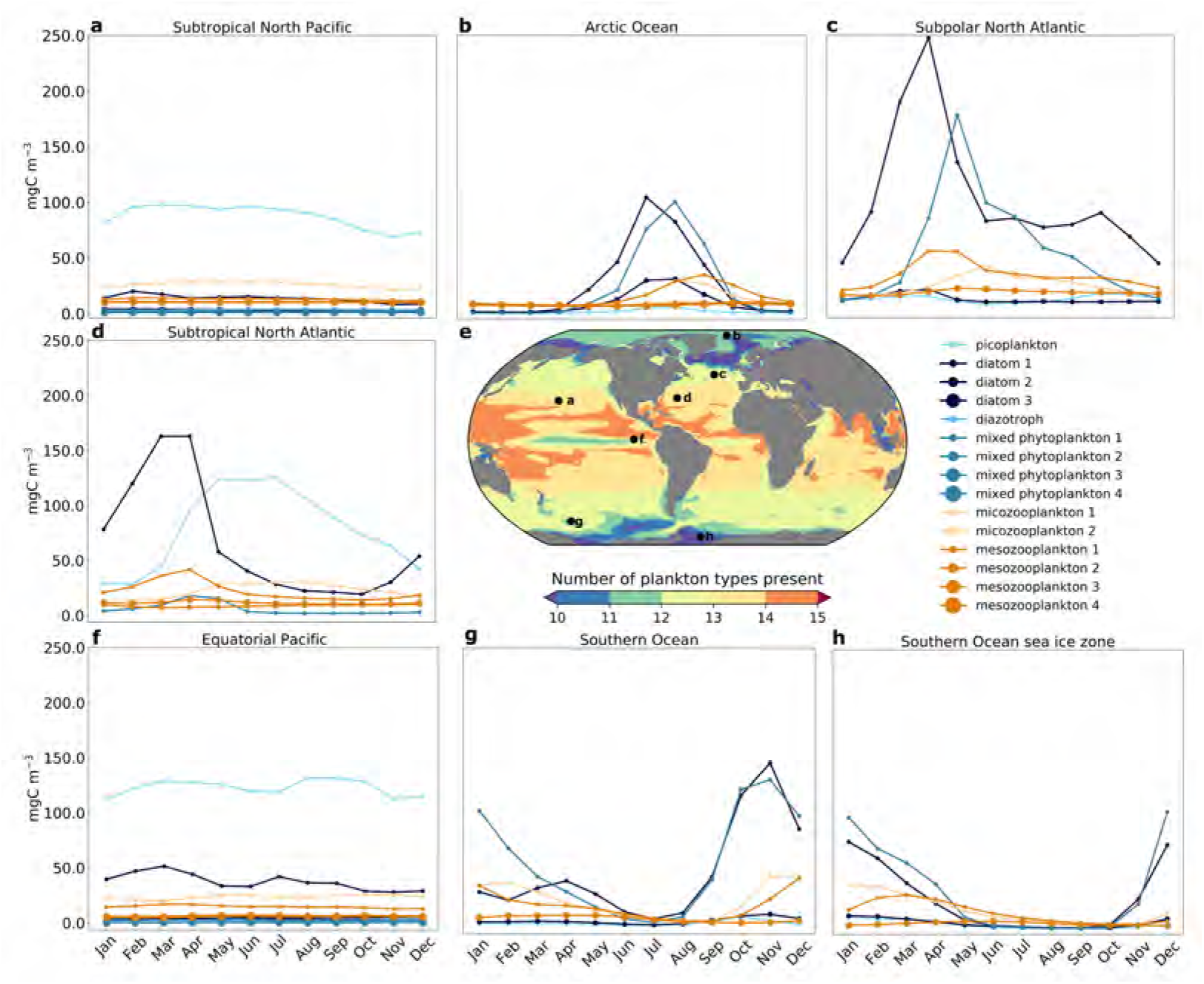
Plankton phenology. Average number of plankton present at a level more than 1% of the total plankton biomass for a given grid cell at any time of the year (e). Seasonal cycles for phytoplankton (blue) and zooplankton (orange) in a 5° by 5° region in the (a) Subtropical North Pacific (47–51*°N*, 165–160*°W*), (b) Arctic Ocean (78–83*°N*, 1–6*°W*), (c) Subpolar North Atlantic (45 – 50*°N*, 27 – 32*°W*), (d) Subtropical North Atlantic (78 – 83*°N*, 1 – 6*°W*), (f) Equatorial Pacific (2*°N*-2*°S*, 97-101*°W*), (g) Southern Ocean (47 – 61*°S*, 171 – 175*°W*), and (h) Southern Ocean sea ice zone (66 – 70*°S*, 38 – 42*°W*).

For example, in the subtropical North Pacific (47 – 51*°N*, 165 – 160*°W*; Fig. 12a), low nutrient availability led to picoplankton dominance throughout the year. Strong grazing pressure from small microzooplankton together with low nutrient delivery allowed for the dominance of relatively small phytoplankton. The Equatorial Pacific (2*°N*-2*°S*, 97-101*°W*; Fig. 12f) was similarly dominated by picoplankton throughout the year with weak seasonality, however showed a higher contribution of small diatoms due to higher nutrient inputs from Equatorial upwelling (Fig. 4). Conversely, in the subpolar North Atlantic region (45 – 50*°N*, 27 – 32*°W*; Fig. 12c), MARBL-SPECTRA simulated a Spring bloom dominated by small diatoms and mixed phytoplankton. The bloom decreases with the emergence of small microzooplankton and mesozooplankton grazing, followed by the development of a fall bloom composed of small diatoms. The Subtropical North Atlantic (78 – 83*°N*, 1 – 6*°W*; Fig. 12d), showed a similar, but weaker spring bloom dominated by the small diatoms, decreasing with the emergence of mesozooplankton grazing. This bloom was followed by a longer fall and summer picoplankton bloom. In the Southern Ocean (47 – 61*°S*, 171 – 175*°W*; Fig. 12g), small diatoms and mixed phytoplankton drive a late Spring/early austral Summer bloom due to high nutrient supply. A similar but shorter bloom occurred in the Southern Ocean sea-ice zone (66 – 70*°S*, 38 – 42*°W*; Fig. 12h) driven by the small diatoms and mixed phytoplankton. In the Arctic Ocean (78 – 83*°N*, 1 – 6*°W*; Fig. 12b), small and medium diatoms and small mixed phytoplankton drive a boreal Summer bloom decreasing with the emergence of microzooplankton and mesozooplankton grazing.

Overall, MARBL-SPECTRA can simulate phenology and succession in a more diverse fashion than models with fewer taxa. We calculated the total number of phytoplankton and zooplankton taxa present at greater than 1% of total biomass of phytoplankton and zooplankton in each month of the year, and averaged this over all months, to find the averaged number of model species present at any time of year. The highest average number of plankton present contributing to more than 1% of plankton biomass are seen in the subtropical gyres, especially near coastal boundary currents (Fig. 12e). The weak seasonality, and high contribution of small phytoplankton and microzooplankton throughout the year might contribute to this greater number of plankton present in the subtropical gyres compared to other regions. Meanwhile, the higher nutrient concentration in the Equatorial upwelling region, drives the opportunist small diatoms to dominate most of the plankton biomass decreasing the number of plankton types present (Fig. 12e & Fig. S3). Polar regions displayed a lower average number of plankton types present throughout the year due to strong seasonality, and higher dominance of larger plankton types (Fig. 12e & Fig. 8).

### 3.10. Sensitivity Analyses

The sensitivity tests illustrated the impacts and feedbacks that variations in allometric scaling exponents have across model plankton. Variations in *α^Chl^* illustrated the strong constraint set by light affinity across phytoplankton biomass. The decreased *α^Chl^* in picoplankton resulted in decreased small phytoplankton biomass (Fig. S12b) in oligotrophic regions of the ocean, while increases in *α^Chl^* in the other phytoplankton contributed to increases in small phytoplankton biomass (Fig. S12b) in subpolar and polar regions, and overall increased large phytoplankton biomass ((Fig. S12e) throughout the globe except in the Western Pacific where decreases in large phytoplankton biomass (Fig. S12e) were due to nutrient limitation decreasing phytoplankton growth. In contrast, a decreased *α^Chl^* in picoplankton resulting from a steeper allometric scaling, led to increased biomass in the small phytoplankton (Fig. S12c). Similarly, increases in *α^Chl^* in all other phytoplankton due to a steeper allometric relationship led to decreased small phytoplankton biomass (Fig. S12c) in polar and subpolar regions, and overall decreases in large phytoplankton (Fig. S12f) except of the Western Pacific region, where higher nutrient availability stimulated growth in large phytoplankton (Fig. S12f). Average differences in phytoplankton biomass scaled similarly to microzooplankton (Fig. S12h,k) and mesozooplankton (Fig. S12i,l) biomass in both cases.

Variations in *k_N_* illustrated the strong dependency on nutrient uptake across phytoplankton biomass and the positive feedback on zooplankton biomass (Fig. S14). The shallower allometric scaling in *k_N_* led to decreased picoplankton biomass (Fig. S14a) especially in oligotrophic regions, due to less efficient nutrient uptake compared to the base model run. Meanwhile, the lower *k_N_* in all other phytoplankton led to biomass increases (Fig. S14d) in most regions of the ocean, except for areas in the Western Pacific, or where NO_3_ and SiO_3_ were most limiting phytoplankton growth. A steeper allometric scaling in *k_N_* with increasing body size showed the opposite trend, however was more sensitive to the increased efficiency in nutrient uptake for picoplankton, and decreased efficiency for all other phytoplankton leading to larger average differences in small (Fig. S14c) and large (Fig. S14f) phytoplankton biomass compared to the base run (Fig. S14a,b). Average differences in phytoplankton biomass scaled similarly to microzooplankton (Fig. S14h,k) and mesozooplankton (Fig. S14i,l) biomass in both cases.

Variations in *r_j_* illustrated the strong constraint zooplankton basal respiration had especially in the microzooplankton, and their feedbacks across plankton groups. The increased basal respiration rates in microzooplankton and small mesozooplankton (Fig. S15) led to decreased microzooplankton biomass throughout the globe (Fig. S16h), leading the decreased mesozooplankton biomass (Fig. S16k) due to lower food availability from decreased microzooplankton. This led to increased small phytoplankton biomass (Fig. S16b) due to decreased grazing pressure from microzooplankton, and decreased large phytoplankton biomass (Fig. S16e) from increased grazing pressure from mesozooplankton. The opposite occurs with a more positive *β_r_j__* (Fig. S16c,f,i,l), but here the decrease in small phytoplankton biomass (Fig. S16c) was caused by decreased basal respiration rates in microzooplankton.

Variations in 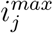 illustrated the negative feedback maximum ingestion rates in zooplankton had on plankton communities, where increases in microzooplankton and small mesozooplankton 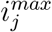 (Fig. S17) led to decreased microzooplankton and small phytoplankton biomass (Fig. S18h,b). Meanwhile, decreases in the larger mesozooplankton 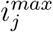 resulted in the opposite relationship (Fig. S16e,k). Increases in the maximum ingestion rate of microzooplankton initially would increase their biomass, but with time, decreases in small phytoplankton biomass (Fig. S18h) from higher grazing pressure led to decreases in microzooplankton biomass (Fig. S18h). Decreases in small phytoplankton alleviated nutrient limitation allowing larger phytoplankton biomass to increase and therefore increased mesozooplankton biomass (Fig. S18k). The opposite occurred with a more positive 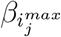 (Fig. S18c,f,i,l), where lower ingestion rates in microzooplankton increased small phytoplankton biomass (Fig. S18c), leading to decreases in large phytoplankton biomass (Fig. S18f) due to increased competition for resources and higher grazing pressure from mesozooplankton due to higher mesozooplankton ingestion rates. The decrease in large phytoplankton biomass (Fig. S18f) led to decreases in mesozooplankton biomass (Fig. S18l) due to decreased food availability.

Similar to variations in 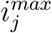, changes in *K^P^* illustrated the negative feedback grazing affinity has across plankton biomass. Changes in grazing affinity were similar to changes in maximum ingestion rates, where increases in grazing affinity (Fig. S20b,e,h,k) led to decreases in small phytoplankton and fed back to lower microzooplankton biomass (Fig. S20h). The alleviation of nutrient limitation from lower small phytoplankton biomass allowed larger phytoplankton to compete, increasing food availability for mesozooplankton (Fig. S20k). Lower grazing affinities (Fig. S20c,f,i,l) increased small phytoplankton biomass (Fig. S20c) by alleviating their grazing pressure. Larger phytoplankton biomass (Fig. S20f) decreased in oligotrophic regions of the ocean, where small phytoplankton can outcompete larger phytoplankton due to their higher nutrient affinities. Lower grazing pressures allowed large phytoplankton (Fig. S20f) to thrive in subpolar and polar regions where there tends to be higher nutrient availability. Higher phytoplankton biomass (Fig. S20c,f) led to increases in microzooplankton (Fig. S20i) in subpolar regions even with their decreased grazing affinity. Mesozooplankton biomass however did decrease in most of the globe due to higher sensitivity to decreased grazing affinity (Fig. S20l).

## 4. Discussion

### 4.1. Model advances

MARBL-SPECTRA extends the MARBL plankton food web to simulate nine phytoplankton and six zooplankton along two trait axes, size and biogeochemical function, such that we are now able to investigate key questions relating to plankton community dynamics that were previously not possible or limited, such as interactions between plankton size structure, phenology, and food web length with biogeochemical cycles. This is a fundamentally different model from the typical plankton functional type model utilized in most ESMs, where simple representations of plankton with different biogeochemical functions are used to resolve large-scale biogeochemical cycles. Here, we have identified a tractable approach for the union of plankton functional type modeling with size-resolved, trait-based models within an ESM, such that community dynamics within the plankton ecosystem can skillfully simulate key ecosystem and biogeochemical properties, such as surface chlorophyll, phytoplankton and zooplankton biomass, and surface nutrients. MARBL-SPECTRA builds upon considerable recent progress in plankton community modeling, including similar efforts that represent diversity across functional group and organism size (Dutkiewicz et al., 2021).

In MARBL-SPECTRA, the inclusion of more plankton functional types and size classes provides a more complex food-web structure (Fig. 1) capable of examining temporal and spatial patterns in plankton ecosystem size structure (Fig. 8), food web length (Fig. 10b), and phenology (Fig. 12), and how these influence the biological pump (Figs. 8c and 10c). Specifically, differentiating picoplankton from other small phytoplankton in MARBL-SPECTRA allowed us to better resolve phenological changes in different regions (Fig. 12). These advantages were particularly apparent in the subtropical vs. subpolar North Atlantic, where diatom blooms were followed by a summertime picoplankton bloom in the subtropics, compared to a late-spring mixed phytoplankton bloom in the subpolar regions (Fig.12c,d; see more in Section 4.1.2). Separating picoplankton from small mixed phytoplankton allowed us to better distinguish between the ecosystem impacts on remineralization and carbon export, particularly in the subtropical biomes.

Additionally, explicit representation of microzooplankton and mesozooplankton size classes allowed us to better resolve top-down controls on ecosystem productivity. The microzooplankton in MARBL-SPECTRA consumed 56% of total primary productivity (consistent with empirical estimates of 62%, (Schmoker et al., 2013), and exerted significant controls on the picoplankton biomass. These dynamics were particularly visible in the sensitivity studies that targeted maximum ingestion rates and ingestion half-saturation constants (Figs. S17–S20). The explicit representation of six size classes of zooplankton further allowed us to examine spatial variations in food chain length (as seen through the mean zooplankton trophic level variables; Fig.10b) and how they influenced the particle export ratio (Fig. 10c; see more in Section 4.1.2). Interestingly, while the subtropical gyres had similar export ratios, the iron-limited South Pacific subtropical gyre was distinct in that it had the longest food chains, with large mesozooplankton exceeding trophic level 4 (Fig. S6).

#### 4.1.1. Allometric Scaling and Feeding Kernel

While some ocean biogeochemical models use size to inform the structure and parameterizations of their plankton ecosystem (e.g. Stock et al., 2020), here we separated the plankton along two trait axes, size and biogeochemical function, with allometric relationships explicitly determining a range of parameters (e.g., metabolic rates, nutrient uptake, light affinity, ingestion rates, C:P ratios) (Figs. 2–3 & Tables 2–3). While it is well established that size is a “master trait” governing the physiology of marine organisms from numerous theoretical, empirical, and modeling studies (Andersen et al., 2016), there is still uncertainty in the scaling exponents for light affinity, nutrient uptake, ingestion rates, metabolic rates, and predator-to-prey size ratios (Litchman et al., 2007; Edwards et al., 2012, 2015b,a; Hansen et al., 1997, 1994). Further, in the marine system, the exceptions to the dominant rules are often highly ecologically important, yielding relationships such as the non-monotonic growth rate in phytoplankton (Marañón et al., 2013; Marañón, 2015).

In developing MARBL-SPECTRA, we applied the principles of allometric scaling for parameters that had a size-dependent relationship, but allowed for those scalings to vary by biogeochemical function. Thus, we ended up with three degrees of freedom for each model parameter (size scaling slope, intercept, and variations by biogeochemical type; Table 2). This allowed for some tuning within the constraints of known parameter uncertainties, which are discussed in more detail in the Methods (Section 2.4). However, due to the complexity of and interactions within the food web, the ecological consequences of tuning parameters for individual model species were not always intuitive. This was evident in the series of ten sensitivity analyses (Section 2.5), particularly the ones where zooplankton parameters varied. In particular, in the sensitivity experiments where the scaling slope of zooplankton maximum ingestion rate became steeper (increasing rates for microzooplankton), the microzooplankton biomass ultimately decreased due to a negative feedback from increasing grazing pressure on small phytoplankton (Fig. S17). A similar effect was also seen for the ingestion half-saturation constants (Fig. S19–S20). These sensitivity cases highlighted the complexities in tuning the ecosystem and how changes in one parameter may have unintended consequences on other model variables. Ultimately, we found that it was not possible to apply allometric scalings without substantial tuning of the ecosystem and still be able to achieve skillful simulations of marine biogeography and biogeochemical fields.

The structure and energy flows within a plankton food web were also inherently tied to predator-prey feeding relationships. Size-spectrum models often represent predator-prey trophic links with a discretization of a plankton feeding kernel, often a unimodal function of prey size (Fuchs and Franks, 2010; Wirtz, 2012; Law et al., 2009; Heneghan et al., 2020). Here, we applied the same principles to MARBL-SPECTRA, with the feeding kernel represented by a mirrored complementary error function that strongly resembled a Laplace distribution. The functional form of feeding kernels can vary, from Gaussian functions (Law et al., 2009; Heneghan et al., 2020) to a Laplace distribution (Fuchs and Franks, 2010), with the broader distributions resulting in more stable ecosystems in idealized simulations (Law et al., 2009). Because the functional form of the plankton feeding kernel is highly uncertain, we opted to use a feeding kernel similar to the Laplace distribution, with predator-to-prey size ratios increasing and the feeding kernel broadening as predator size increased (Hansen et al., 1997). However, similar to the plankton physiological rates, we also tuned the predator-prey feeding relationships to achieve realistic plankton biogeography and nutrient fields.

#### 4.1.2. Plankton community structure

The increased ecosystem complexity in MARBL-SPECTRA enabled improved simulation of plankton community structure, its drivers, and interactions with biogeochemistry. The community size structure, as evidenced by the size-abundance relationship, showed more negative slopes (more small plankton) in the lower latitudes and oligotrophic gyres and less negative slopes (more large plankton) in the high latitudes (Fig. 8). These results are broadly consistent with other models (Barton et al., 2013; Serra-Pompei et al., 2022) and observations (Kiko et al., 2022). Additionally, we found a positive relationship between the slope of the size-abundance relationship and export efficiency (pe-ratio), similar to Serra-Pompei et al. (2022). Model plankton communities with steeper (more negative) slopes were dominated by small phytoplankton with inefficient food webs, and thus had lower export efficiencies than plankton communities dominated by larger phytoplankton, consistent with observations (Henson et al., 2019).

Similarly, we found substantial spatial gradients in plankton food chain length, as evidenced by the zooplankton mean trophic level (Fig. 10b). Because MARBL-SPECTRA simulated fixed size classes, the mean trophic level for the modeled zooplankton indicated the number of trophic links within the range of simulated sizes. In low-productivity waters, picoplankton were the dominant phytoplankton type (Fig. 7), with microzooplankton as their main predators, consuming 75% of the primary production in oligotrophic regions. The remaining production was channeled directly through mesozooplankton or lost to sinking and other advective processes. The tight coupling between phytoplankton, microzooplankton, and mesozooplankton resulted in longer food chains in oligotrophic regions composed of more trophic levels (Fig. 10b) compared to other regions in the ocean. Oligotrophic regions in the model favored the recycling of organic matter rather than its efficient transfer upward toward higher trophic levels (Azam et al., 1983; Legendre and Le Fèvre, 1995). Meanwhile, productive regions were characterized by shorter trophic pathways (Fig. 10b), with a larger fraction of particulate organic carbon exported from the euphotic zone (Fig. S9). This was caused by the sinking of ungrazed cells or indirectly through the sedimentation of aggregated detritus and zooplankton fecal pellets, resulting in a biological pump more efficient in transporting biogenic carbon towards the ocean interior (Guidi et al., 2009; Boyd and Trull, 2007). Accordingly, we found a general negative relationship between plankton food chain length and export efficiency (Fig. 10c), such that shorter food chains (lower mean zooplankton trophic levels) were associated with higher export efficiency, with the opposite generally true for longer food chains. However, in the iron limited South Pacific oligotrophic gyre, export efficiencies < 0.1 were associated with longer food chains than in the other oligotrophic gyres. This was caused by iron limitation reducing the production of diatoms and other large phytoplankton that are grazed by mesozooplankton.

The increased model resolution of phytoplankton and zooplankton size classes enabled us to study the relative abundance of predators and prey across regions of contrasting productivity. Zooplankton to phytoplankton biomass ratios (Z:P) were consistent with a 3/4 scaling exponent observed by Hatton et al. (2015), with zooplankton biomass increasing at a lower rate than phytoplankton biomass. Coastal upwelling and other productive regions of the ocean displayed lower zooplankton to phytoplankton biomass fractions compared with oligotrophic regions of the ocean. The decrease in Z:P ratio with a eutrophication gradient was consistent with observations (Gasol et al., 1997; Yuan and Pollard, 2018; Hatton et al., 2015), but deviates from other modeling analyses that show an increase in Z:P ratio with a eutrophication gradient (Ward et al., 2014; Vallina et al., 2014). One reason for lower Z:P ratios in productive regions could be due to the longer generation times of mesozooplankton (weeks to months) compared to microzooplankton (days), which may impede them from thriving in upwelling regions where strong fluctuations in food supply and environmental conditions occur. Additionally, the use of the Holling Type II grazing function, which keeps predation pressure relatively high at low prey concentrations, may prevent mesozooplankton production from decreasing too much in oligotrophic regions of the ocean. Another reason for this deviation could be due to the high sensitivity of zooplankton biomass to basal respiration and quadratic mortality values in the model. Higher zooplankton quadratic mortalities for the mesozooplankton reflect higher trophic level grazing. The high mortality values can therefore decrease mesozooplankton biomass especially in upwelling regions of the ocean, contributing to a weaker dynamic range in mesozooplankton biomass between oligotrophic and eutrophic regions.

The more diverse plankton community in MARBL-SPECTRA simulated the seasonal succession of plankton communities tied to nutrient delivery, temperature, light availability, and grazing pressure. In temperate regions, model diatoms dominated the spring bloom with the onset of thermal stratification increasing light availability (Dandonneau et al., 2006; Knox, 2006; Siegel et al., 2002; Hirata et al., 2011). Sufficient light and nutrient supply aided the rapid growth of the smallest diatoms (Litchman et al., 2007; Edwards et al., 2015a). Mixed phytoplankton developed in late spring following strong microzooplankton and mesozooplankton grazing pressure on diatoms (Dandonneau et al., 2006; Knox, 2006; Calbet and Landry, 2004). In autumn, a weaker small diatom bloom occurred in many regions, driven by nutrient delivery to the surface due to enhanced mixing under favorable light conditions (Hirata et al., 2011). In polar regions, this small diatom bloom was shifted towards boreal and austral summer due to lower light availability and sea ice dynamics influencing phytoplankton growth (Knox, 2006). The small diatoms and mixed phytoplankton dominated the onset of the bloom, but larger diatoms still contributed to overall biomass due to high nutrient concentrations. This bloom declined with increased microzooplankton and mesozooplankton grazing and decreased in light availability towards the end of the summer. Tropical and subtropical regions displayed a weaker seasonality in phytoplankton blooms due to more stable light availability throughout the year and lower nutrient concentrations (Alvain et al., 2008; Hirata et al., 2011). Throughout the tropics and subtropics, but especially in oligotrophic regions of the ocean, picoplankton dominated throughout the year, with lower contributions of mixed phytoplankton and diatoms to overall biomass. These results were consistent with ocean color remote sensing (Alvain et al., 2008; Bracher et al., 2009; Hirata et al., 2011), field observations (Leblanc et al., 2012; Buitenhuis et al., 2013), as well as other modeling studies (Ward et al., 2012; Dutkiewicz et al., 2009).

### 4.2. Limitations & future improvement

All plankton community models, including MARBL-SPECTRA, are simplifications of natural plankton communities that seek to simulate phytoplankton physiology, predator-prey interactions, community structure, and biodiversity in a dynamic environment. MARBL-SPECTRA incorporates nine phytoplankton and six zooplankton types, where the traits of organisms and their interactions are determined by organism size and functional group, in the case of phytoplankton. While this approach is computationally tractable and allows for the study of lower tropic levels in the marine environment, it has several important limitations.

First, our model does not account for zooplankton life histories such as diapause. Diapause is a critical component of the life history of copepods, as it allows them to survive long periods of unfavorable environmental conditions (Hairston Jr and Munns Jr, 1984). Copepods accumulate lipid reserves prior to diapause, and are highly nutritious prey for a wide variety of predators in the oceans (Bauermeister and Sargent, 1979). Diapausing copepods are especially important in polar, subpolar, and temperate environments where *Calanoid* copepods are a key intermediary in the process of trophic energy transfer from phytoplankton to higher trophic levels (Baumgartner and Tarrant, 2017). The exclusion of zooplankton life histories can bias mesozooplankton biomass in polar, subpolar, and temperate regions, particularly in the spring (Fig. 9), when copepods are emerging from diapause. As a consequence, there may be insufficient top-down control on the spring phytoplankton bloom, thought to be one of the key mechanisms controlling bloom timing (Banse, 2013). Additionally, MARBL-SPECTRA does not differentiate zooplankton carbon content within zooplankton size classes. This limits the nutritional value and growth potential for higher trophic levels, because carbon content in zooplankton varies substantially. These variations in carbon content are especially seen between gelatinous groups with low carbon density to non-gelatinous zooplankton with high carbon density (Kiørboe, 2013; McConville et al., 2017). Including differentiation in zooplankton carbon content can impact the relative fitness of zooplankton groups, since critical physiological and competitive processes such as metabolism, search rate, and average growth efficiency scale with carbon across zooplankton groups (Acuña et al., 2011; Kiørboe, 2011; Kiørboe and Hirst, 2014; McConville et al., 2017; Heneghan et al., 2020).

Second, some key phytoplankton and zooplankton functional groups are absent from the model. Calcifying phytoplankton are a key functional group important in the carbon cycle, producing more than half of the marine carbonate flux (Schiebel, 2002). Although MARBL-SPECTRA accounts for this group implicitly, the inclusion of explicit calcifiers could improve the spatial and temporal representation of calcium carbonate production, as well as incorporate key carbon fertilization mechanisms thought to buffer coccolithophore responses to climate change (Krumhardt et al., 2017, 2019). In addition, phytoplankton dimethyl sulfide (DMS)-producers influence the atmospheric sulfur cycle by producing dimethysulfoniopropionate (DMSP) and convert it into DMS using an extracellular enzyme (DMSP-lyase) (Stefels et al., 1995). *Phaeocystisantarctica* is especially important in the Southern Ocean, where it has been observed to dominate the community during blooms (Alvain et al., 2008). The high prevalence in *Phaeocystis* blooms make it an important contributor to primary production and biogeochemical cycles where it occurs. The explicit incorporation of gelatinous zooplankton, such as *Cnidarian* jellyfish and *salps*, could improve the representation of top-down control on prey and the representation of carbon transfer efficiency to depth (Luo et al., 2020). The ability of multiphagous gelatinous zooplankton to feed across a wide spectrum of size classes would provide an indirect route of carbon flux by which even small phytoplankton biomass can be transferred to the deep ocean (Luo et al., 2022).

Third, our model does not include zooplankton vertical migration, the active transport of organic carbon to depth by zooplankton consuming organic particles at the surface during the night and respiring the inorganic nutrients below the mixed layer during the day (Steinberg et al., 2000; Longhurst and Harrison, 1988). While the global inventory of carbon export is constrained in models by ocean circulation and the upward flux of nutrients driving new production, zooplankton diel vertical migration could be an important component in mesopelagic zones, contributing significantly to oxygen consumption, particularly at oxygen minimum zones, and carbon export into the ocean interior (Bianchi et al., 2013; Aumont et al., 2018).

Fourth, MARBL-SPECTRA does not represent mixotrophy. Mixotrophs combine the autotrophic use of light and inorganic resources with the heterotrophic ingestion of prey. The incorporation of mixotrophy in ecological models enhances the transfer of biomass to larger organisms at higher trophic levels, which in turn increases the efficiency of oceanic carbon storage through the production of larger, faster-sinking, and carbon-enriched organic detritus (Ward and Follows, 2016). The exclusion of mixotrophy decreases the production of larger phytoplankton, because the nutrient affinity of plankton decreases with increasing organism size (Fig. 2). The highly efficient uptake of the small phytoplankton leaves insufficient nutrients to support photosynthesis in the larger groups, especially the mixed phytoplankton group.

Fifth, MARBL-SPECTRA does not include an explicit representation of higher trophic levels (fish, carnivorous jellies, etc.). Zooplankton losses to consumption by higher predators are implicitly modeled using a squared mortality term, which has a tendency to stabilize food webs (Edwards, 2001). Feedbacks between the higher trophic level predator and zooplankton are not resolved. One implication of this simplification is that the ecosystem effects of fishing, for example, cannot be resolved directly by MARBL-SPECTRA.

Lastly, bacterial activity is not explicitly modeled in our ecosystem model. The effect of the microbial loop is included through constant degradation rates of bacterial remineralization. Mortality and exudation losses are recycled to inorganic nutrients via constant rate degradation of several pools of organic matter (dissolved and particulate) for each essential element. Modeling bacterial activity explicitly would increase the model’s realism at capturing the microbial food web dynamics, but it should not significantly change our results because bacterial abundances are generally more stable than phytoplankton abundances seasonally in open-ocean waters (Spitz et al., 2001).

### 4.3. Outlook

Plankton community models embedded in ocean biogeochemical and circulation models are powerful tools for examining how organism traits shape species biogeography, interactions within plankton communities, the impacts of environmental changes on marine ecosystems, and feedbacks between ecosystems and biogeochemical cycles (e.g., Kwiatkowski et al., 2020; Follows et al., 2007; Ward et al., 2012; Dutkiewicz et al., 2015b). Here, we have developed and evaluated MARBL-SPECTRA, a trait-based plankton community model that resolves nine phytoplankton sizes classes across four functional groups and six zooplankton size classes, allowing for an enhanced understanding of the underlying mechanisms regulating marine plankton biogeography, and the community’s role in biogeochemical cycles. Future increases in ocean temperatures and other environmental properties are expected to modify phytoplankton community diversity and distribution through a range of direct and indirect pathways and mechanisms, many of which are simulated in MARBL-SPECTRA. The future incorporation of MARBL-SPECTRA in a fully coupled climate model would allow for the projection of model organism fitness into the future to better predict changes in plankton communities structure, biogeochemical cycles, food web dynamics, and air-sea fluxes of climate-active gases.

## 5. Acknowledgements

We acknowledge high-performance computing support from Cheyenne (doi:10.5065/D6RX99HX) provided by NCAR’s Computational and Information Systems Laboratory, sponsored by the National Science Foundation (NSF). This manuscript is based upon work supported by the National Center for Atmospheric Research, which is a major facility sponsored by the National Science Foundation under Cooperative Agreement No. 1852977, and the National Science Foundation Graduate Research Fellowship under Grant No. (DGE-2038238 and DGE-1650112). Any opinions, findings, and conclusions or recommendations expressed in this material are those of the author(s) and do not necessarily reflect the views of the National Science Foundation. In addition to NSF funds to NCAR, MARBL development has been supported by the Office of Biological & Environmental Research (BER) within the Department of Energy (DOE) (DE-SC0012603). We also gratefully acknowledge comments from John Dunne and Charles Stock on this manuscript. Code for generating the namelist parameters for MARBL-SPECTRA are available at: https://github.com/marbl-ecosys/spectra-config. Data from the MARBL-SPECTRA simulations performed for this study are available at: https://doi.org/10.6075/J0NK3F6D. All codes used for analysis are available at: https://github.com/gabyneg/SPECTRA.analysis/.

## 6. Supplementary Information

**Figure S1:**
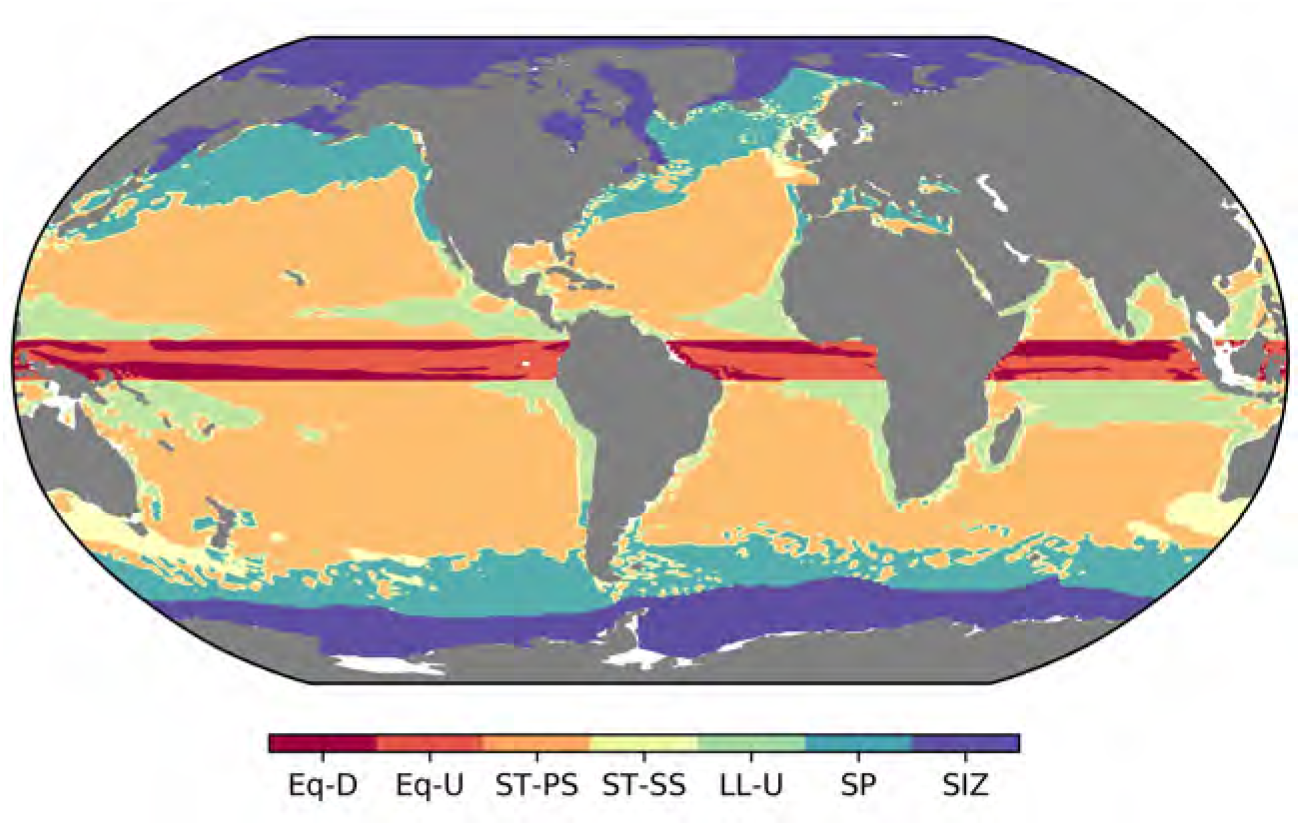
Designated oceanic biomes following Sarmiento et al. (2004): Equatorial downwelling (Eq-D), Equatorial upwelling (Eq-U), Subtropical permanently stratified (ST-PS), Subtropical seasonally stratified (ST-SS), Lower latitude upwelling (LL-U), Sub-polar seasonally stratified (SP), Seasonal ice-covered zone (SIZ). In the analyses, the northern and Southern Hemisphere biomes are separated.

**Figure S2:**
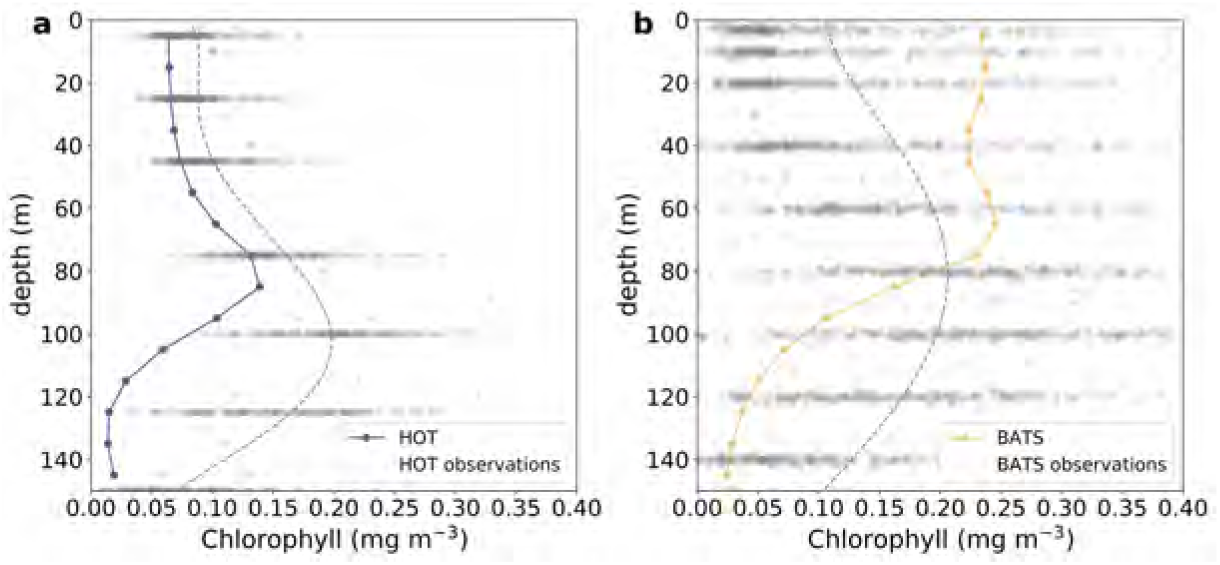
Modeled annual average chlorophyll (mg m^−3^) vertical profiles at the (a) BATS (purple) and (b) HOT (orange) stations over the period 1990 and 2009 compared with observations over the same time period (gray dots and dashed lines) The dashed gray lines represents a line of best fit to observational data, estimated using a fifth order polynomial. BATS chlorophyll data are found at http://batsftp.bios.edu/BATS/bottle/bats_pigments.txt. HOT chlorophyll data are found at https://hahana.soest.hawaii.edu/FTP/hot/primary_production.

**Table S1:** Remineralization length scales (in meters) for sinking particulate matter as a function of depth. The 100 m value is also used above that depth, and the 1000 m value is also used at deeper depths; for all values in between, the length scale is linearly interpolated from the values in the table.

| depth (m) | POC | SiO<sub>2</sub> | CaCO<sub>3</sub> |
| --- | --- | --- | --- |
| 100 | 100 | 770 | 500 |
| 250 | 300 | 2310 | 1500 |
| 500 | 450 | 3465 | 2250 |
| 1000 | 550 | 4235 | 2750 |

**Figure S3:**
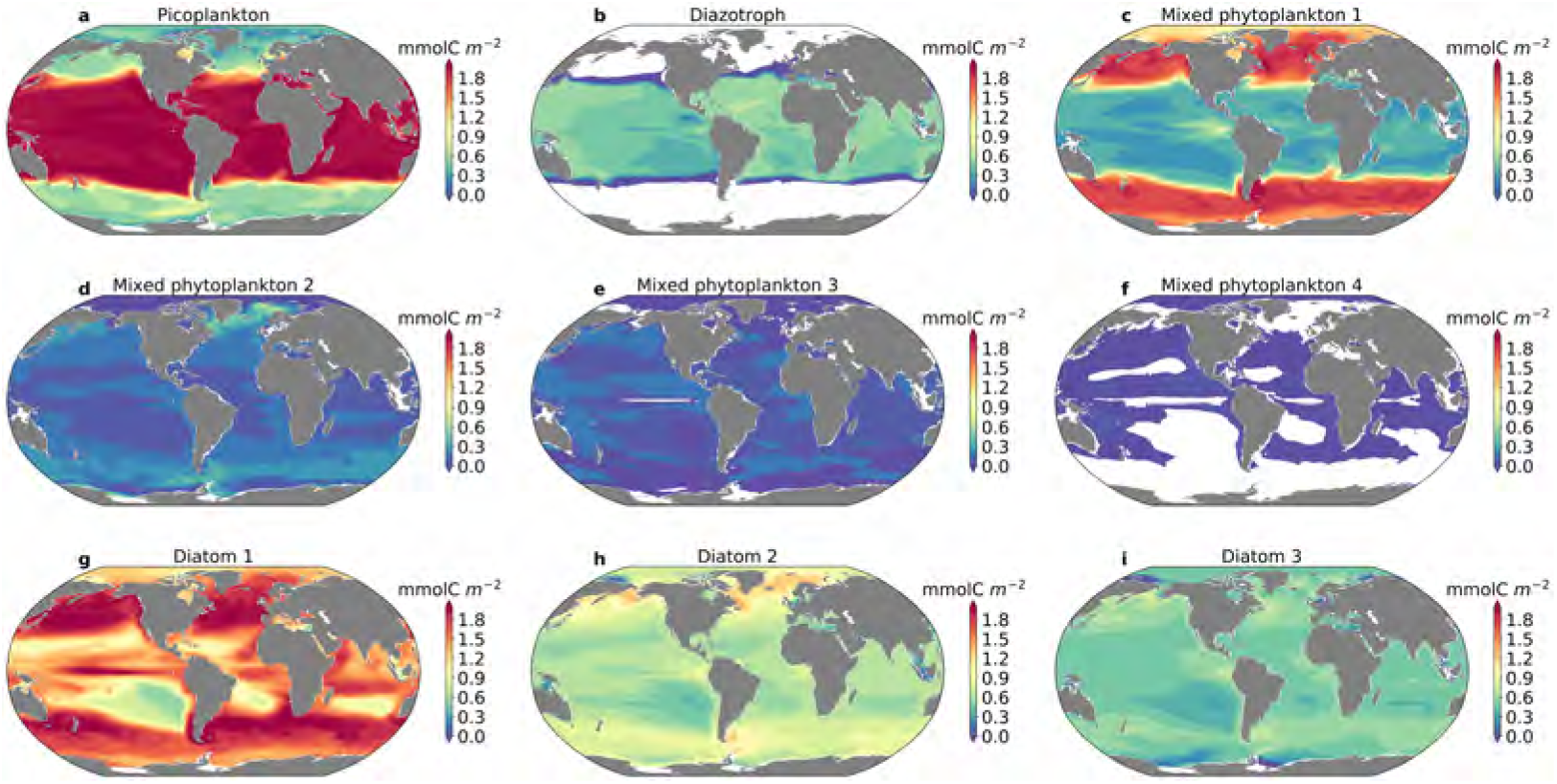
Depth integrated average annual phytoplankton biomass for each phytoplankton type: (a) picoplankton (pp) (b) diazotrophs (diaz), (c) smallest mixed phytoplankton (mp1), (d) second smallest mixed phytoplankton (mp2), (e) second largest mixed phytoplankton (mp3), (f) largest mixed phytoplankton (mp4), (g) smallest diatom (diat1), (h) medium diatom (diat2), and (i) largest diatom (diat3).

**Figure S4:**
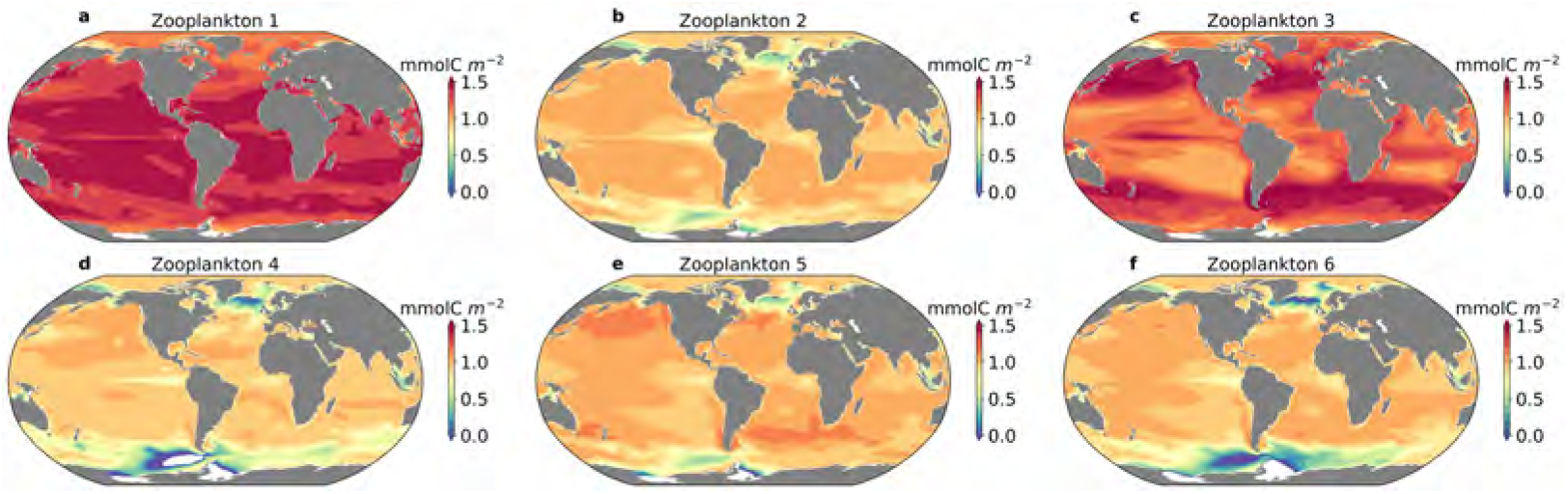
Depth integrated average annual zooplankton biomass for each zooplankton type: (a) smallest microzooplankton (zoo1) (b) largest microzooplankton (zoo2), (c) smallest mesozooplankton (zoo3), (d) second smallest mesozooplankton (zoo4), (e) medium mesozooplankton (zoo5), and (f) largest mesozooplankton (zoo6).

**Figure S5:**
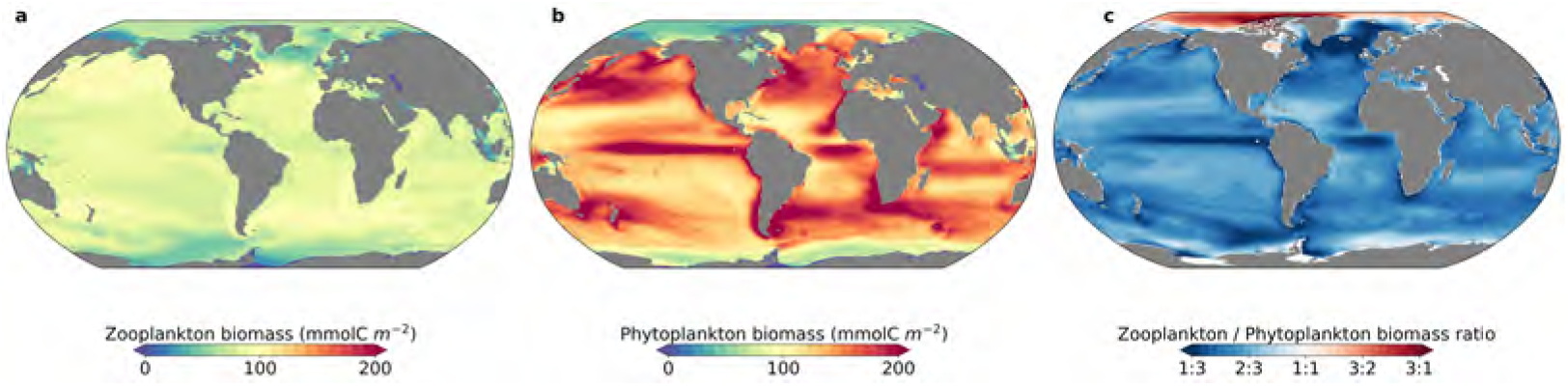
Depth integrated carbon biomass for (a) zooplankton (mmol C m^−2^) and (a) phytoplankton (mmol C m^−2^); (c) the ratio of depth-integrated zooplankton to phytoplankton biomass.

**Figure S6:**
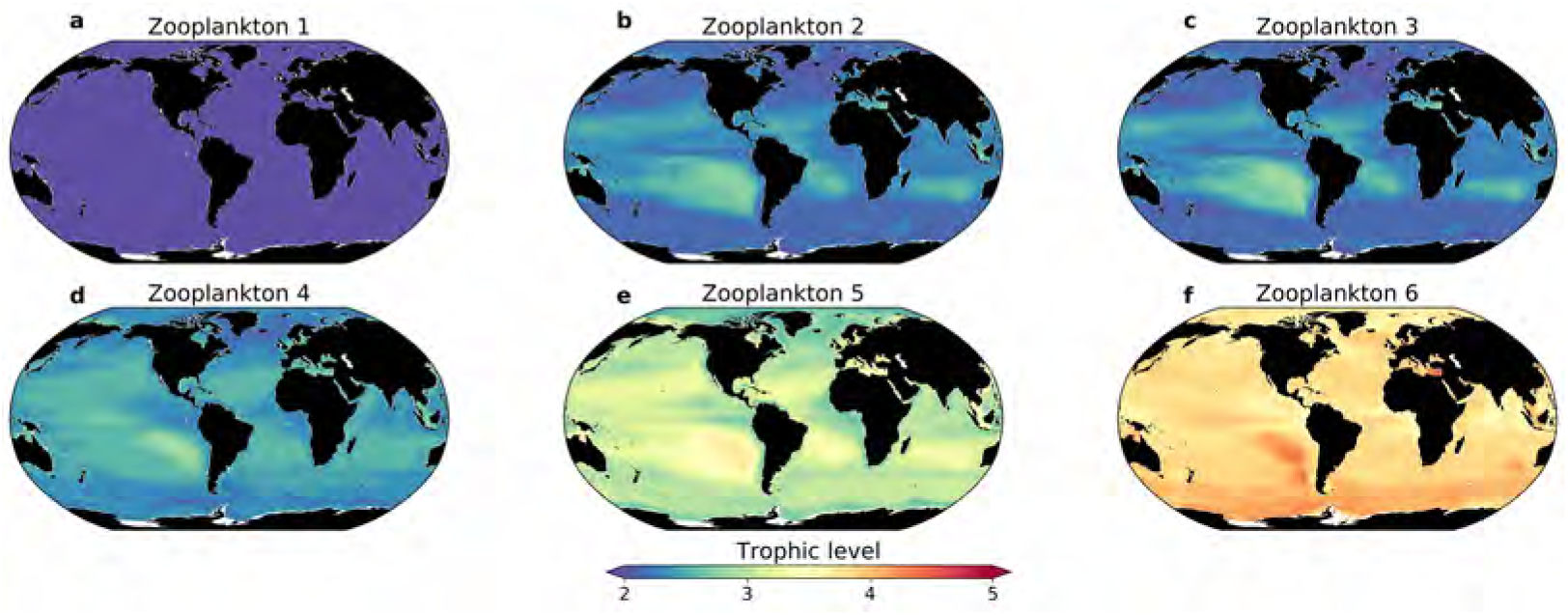
Zooplankton annual mean trophic level over the top 150 m between 1990 and 2009. (a) smallest microzooplankton (zoo1) (b) largest microzooplankton (zoo2), (c) smallest mesozooplankton (zoo3), (d) second smallest mesozooplankton (zoo4), (e) medium mesozooplankton (zoo5), and (f) largest mesozooplankton (zoo6).

**Figure S7:**
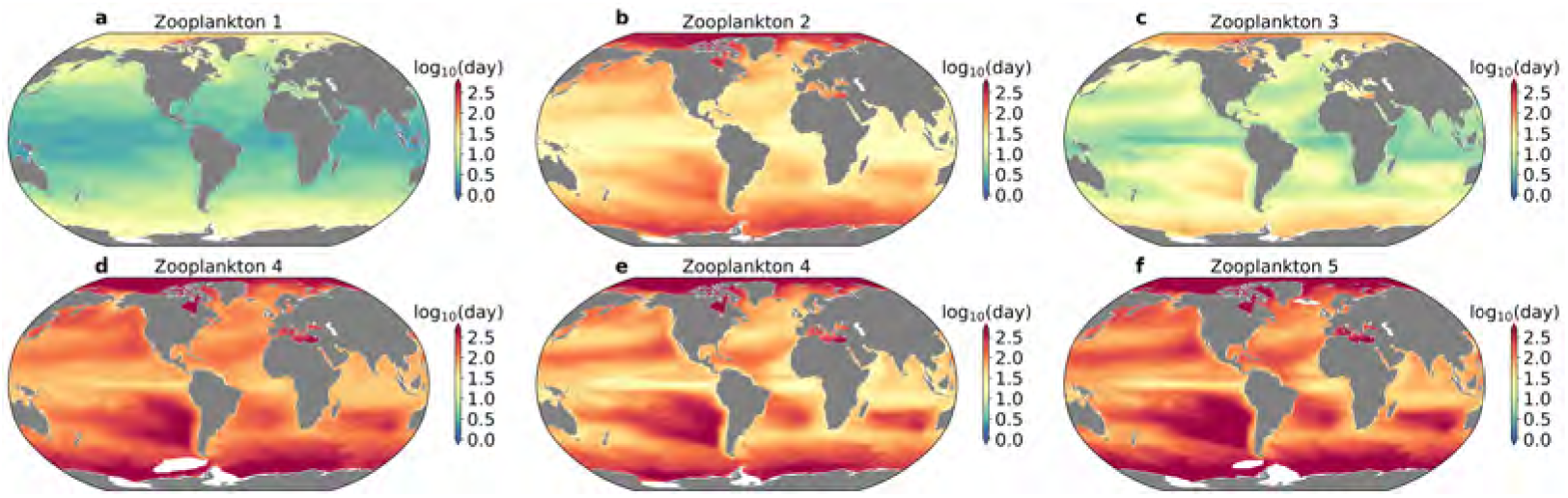
Zooplankton average annual generation time over the top 150m for each zooplankton type: (a) smallest microzooplankton (zoo1) (b) largest microzooplankton (zoo2), (c) smallest mesozooplankton (zoo3), (d) second smallest mesozooplankton (zoo4), (e) medium mesozooplankton (zoo5), and (f) largest mesozooplankton (zoo6). Values are shown in log_10_ days.

**Figure S8:**
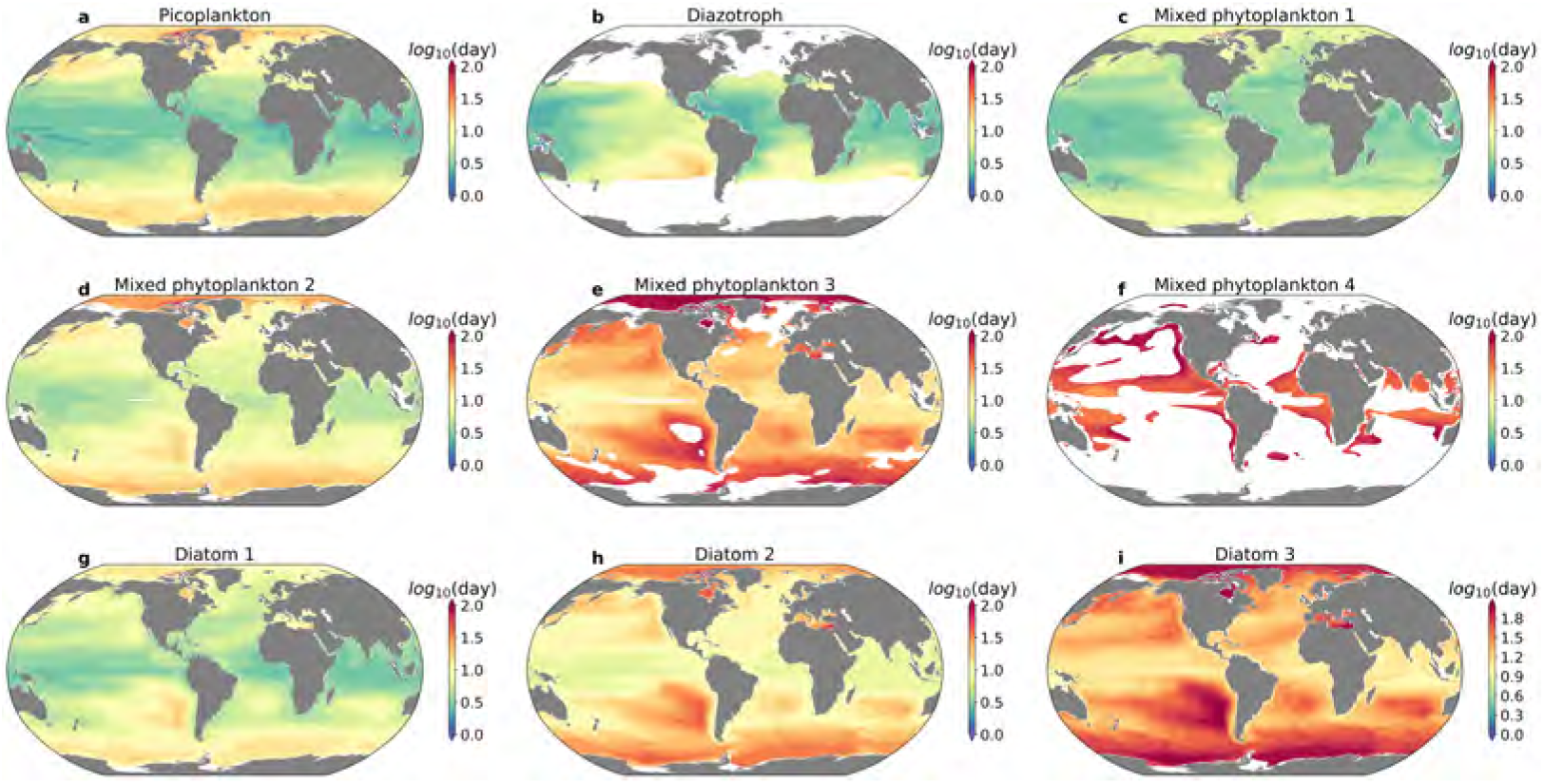
Phytoplankton average annual generation time over the top 150m for each phytoplankton type: (a) picoplankton (pp) (b) diazotrophs (diaz), (c) smallest mixed phytoplankton (mp1), (d) second smallest mixed phytoplankton (mp2), (e) second largest mixed phytoplankton (mp3), (f) largest mixed phytoplankton (mp4), (g) smallest diatom (diat1), (h) medium diatom (diat2), and (i) largest diatom (diat3).

**Figure S9:**
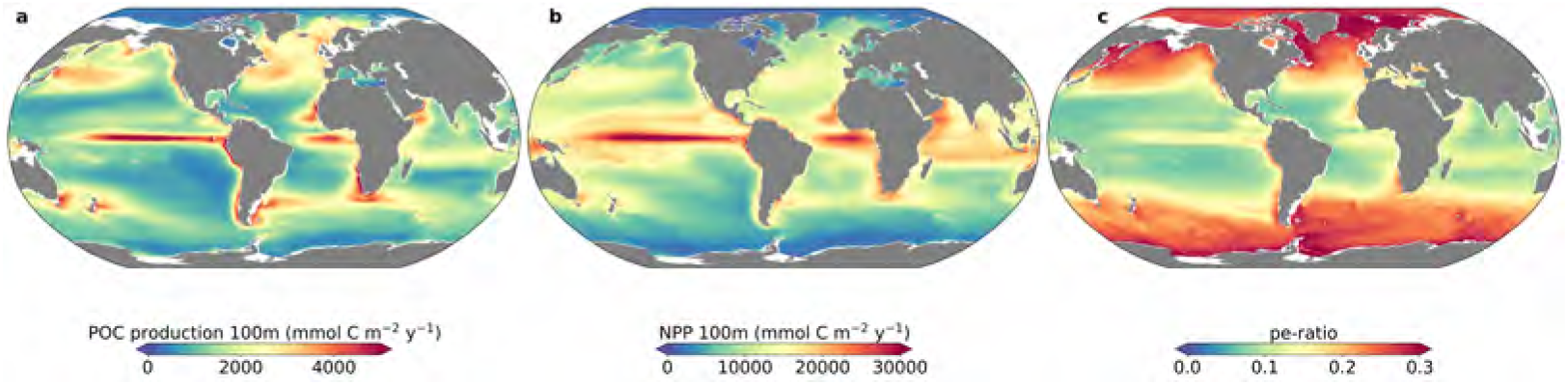
(a) Annual mean particulate organic carbon (POC) production at 100m, (b) annual mean NPP over the top 100m, and (c) pe-ratio (defined as the fraction of depth-integrated NPP exported as sinking particles at 100m) at the top 100m between 1990 and 2009.

**Figure S10:**
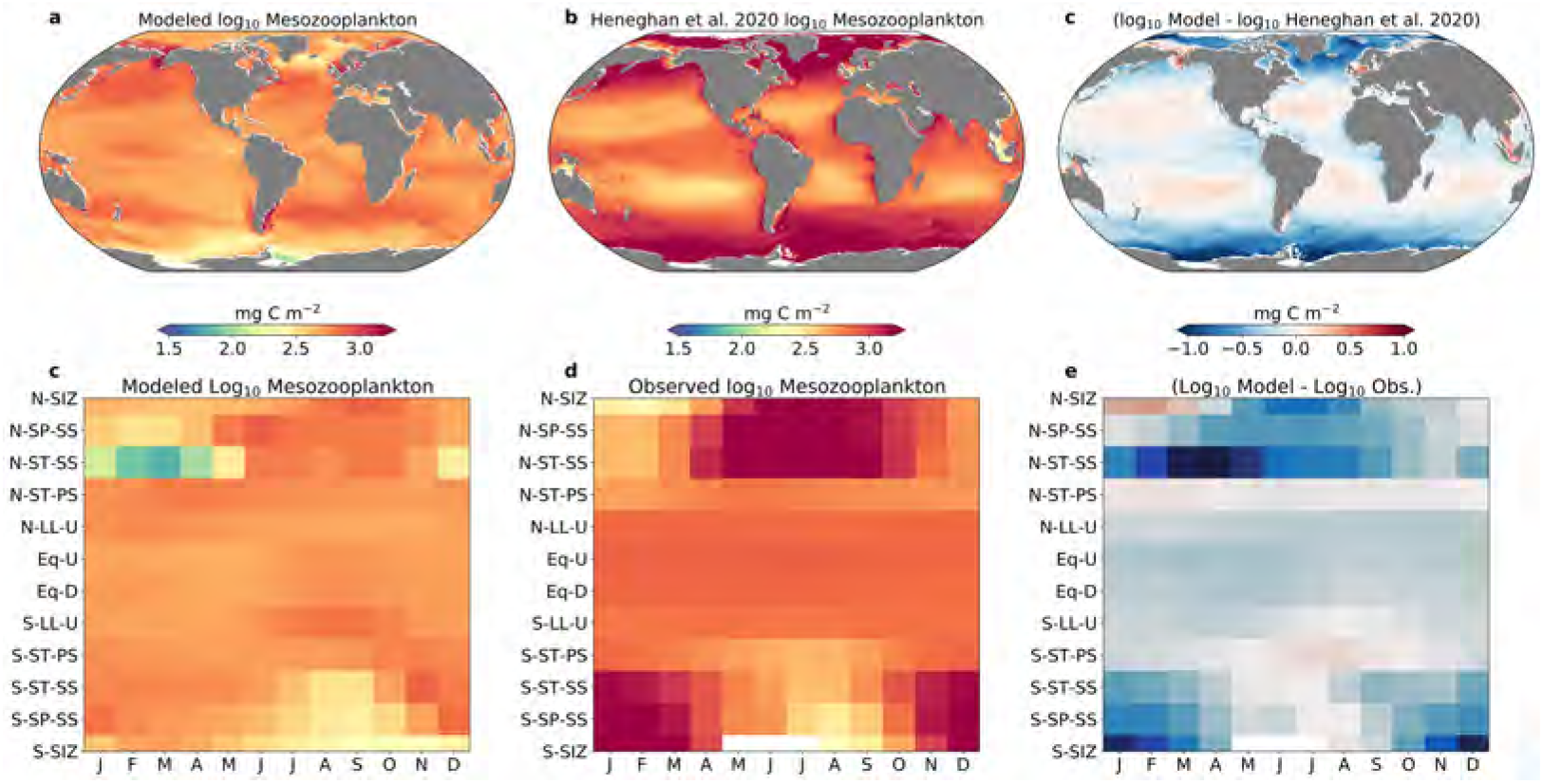
(a) Modeled annual mean mesozooplankton biomass depth-integrated (mg C m^−2^) over the top 150 m, (b) Annual average mesozooplankton biomass (mg C m^−2^) integrated over top 200m from the empirical model of Heneghan et al. (2020), and (c) difference between model and empirical model. (d) Mean monthly modeled surface mesozooplankton biomass by biomes; (e) mean monthly Heneghan et al. (2020) mesozooplankton biomass by biomes; and (f) difference between model and empirical model.

**Figure S11:**
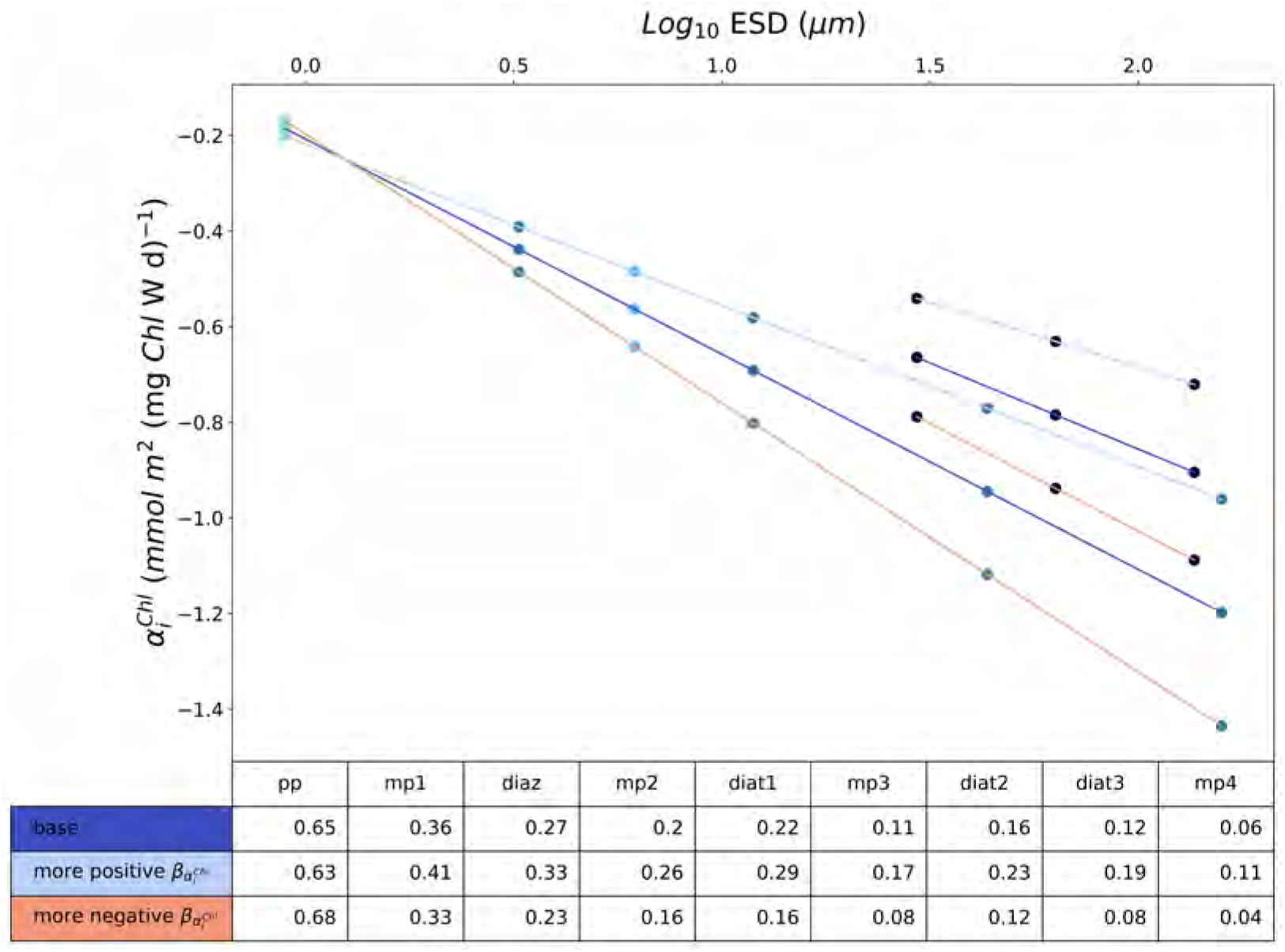
Sensitivity analysis of altering the allometric scaling exponent of the initial slope of the photosynthesis-irradiance curve (*β_α^Chl^_*). The allometric relationship of *α^Chl^* in MARBL-SPECTRA (dark blue) is compared with a more positive (light blue) and more negative *β_α^Chl^_* (light red). The more positive scaling exponent (light blue) increases the *α^Chl^* for all phytoplankton except for picoplankton. The opposite occurs with a more negative scaling exponent (light red), increasing *α^Chl^* for picoplankton and decreasing it for all other phytoplankton. The table shows the exact *α^Chl^* values for each sensitivity test. The different colored dots represent the different functional types of phytoplankton as in Figure 1. The following abbreviations apply for Figs. S11, S13, S15, S17, and S19: pp=picoplankton, mp[1-4]=mixed phytoplankton groups (1=smallest, 4=largest), diaz=diazotrophs, diat[1-3]=diatom groups (1=smallest, 3=largest)

**Figure S12:**
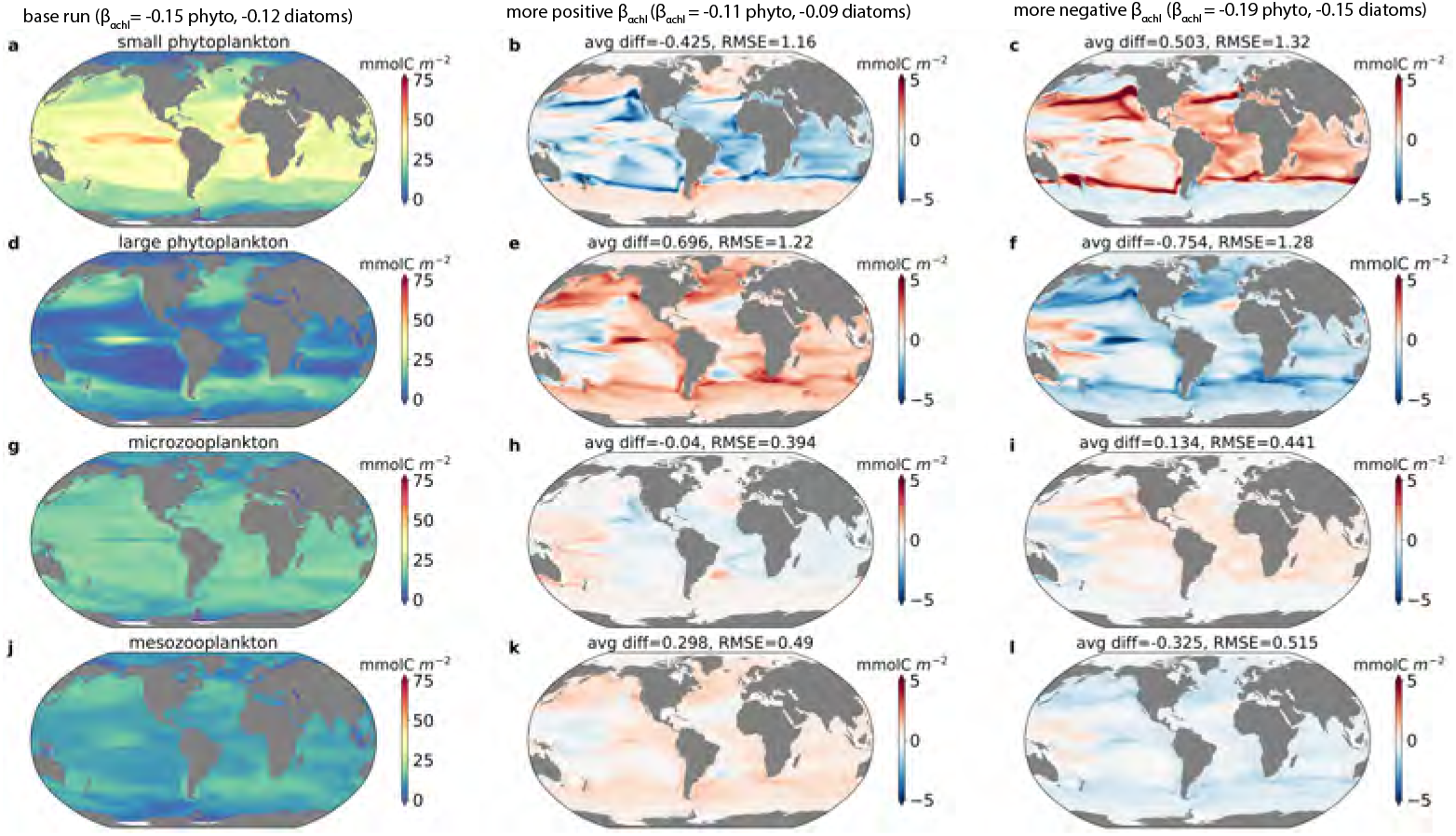
Depth-integrated average annual biomass for a) small phytoplankton (picoplankton, small mixed phytoplankton group, and diazotrophs), d) large phytoplankton (mixed phytoplankton 2, 3, and 4, and all diatoms), g) microzooplankton (smallest two zooplankton) and j) mesozooplankton (largest four zooplankton) in the base MARBL-SPECTRA run. b,e,h, and k) show the biomass differences between the base run minus the more positive slope run. c,f,i, and l) show the biomass differences between the base run minus the more negative slope run. In the base run (a,d,g,j), *β_α^Chl^_* =-0.15 for all phytoplankton and −0.12 for diatoms. In the more positive run, *β_α^Chl^_*)=-0.11 for all phytoplankton and −0.09 for diatoms. In the more negative run, *β_α_PI__* =-0.19 for all phytoplankton and −0.15 for diatoms. Avg diff is the average global difference in biomass and RMSE represents the root mean square error between the base run and each sensitivity run.

**Figure S13:**
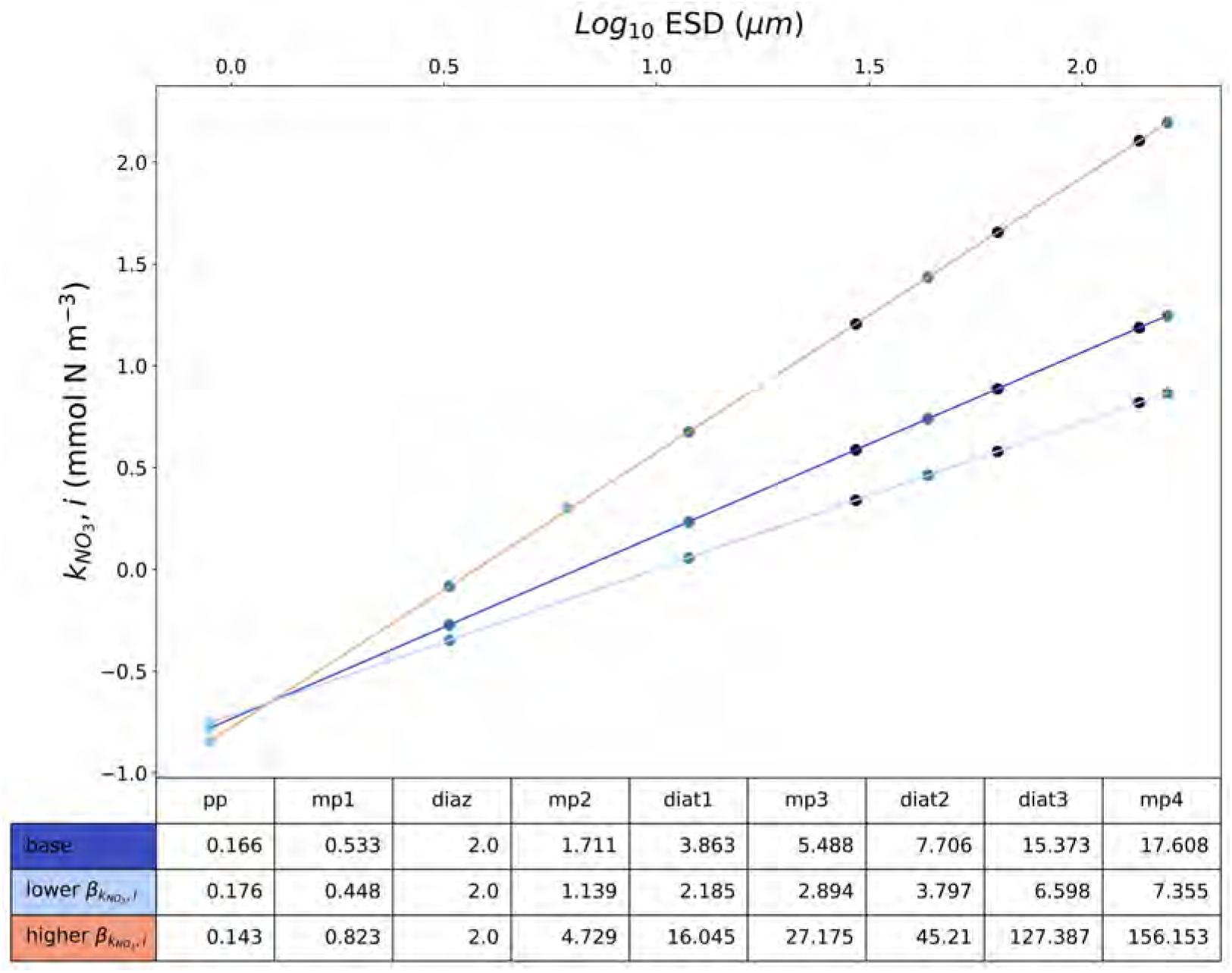
Sensitivity analysis of altering the allometric scaling exponent of all half-saturation constants for nutrient uptake (*β_k_N__*); only 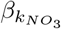 is shown here). The allometric relationship of *k*_*NO*_3__ in MARBL-SPECTRA (dark blue) is compared with a lower (light blue) and a higher 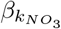 (light red). The lower scaling exponent (light blue) decreases the *k*_*NO*_3__ for all phytoplankton except for picoplankton. The opposite occurs with a higher scaling exponent (light red), which increases *k*_*NO*_3__ for all phytoplankton except for picoplankton. The table shows the exact *k*_*NO*_3__ values for each sensitivity test. The different colored dots represent the different functional types of phytoplankton as in Figure 1.

**Figure S14:**
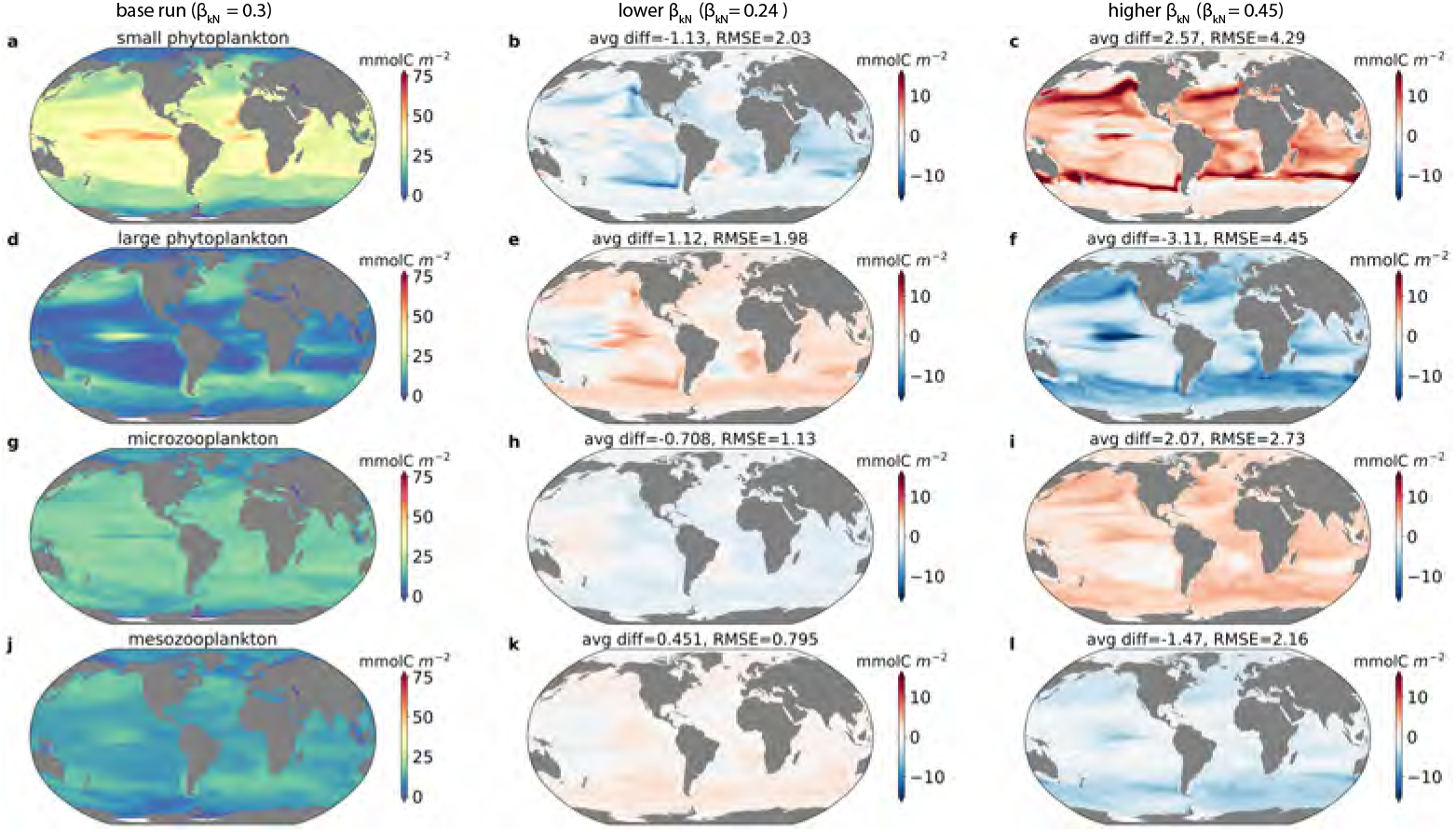
Depth-integrated average annual biomass for a) small phytoplankton (picoplankton, small mixed phytoplankton group, and diazotrophs), d) large phytoplankton (mixed phytoplankton 2, 3 and 4, and all diatoms), g) microzooplankton (smallest two zooplankton) and j) mesozooplankton (largest four zooplankton) in the base MARBL-SPECTRA run. b,e,h, and k) show the biomass differences between the base run minus the lower slope run. c,f,i, and l) show the biomass differences between the base run minus the higher slope run. In the base run (a,d,g,j), *β_k_N__* =0.3 for all phytoplankton. In the lower slope run, *β_kN_* = 0.24 and in the higher slope run *β_k_N__* =0.45 for all phytoplankton. Avg diff is the average global difference in biomass and RMSE represents the root mean square error between the base run and each sensitivity run.

**Figure S15:**
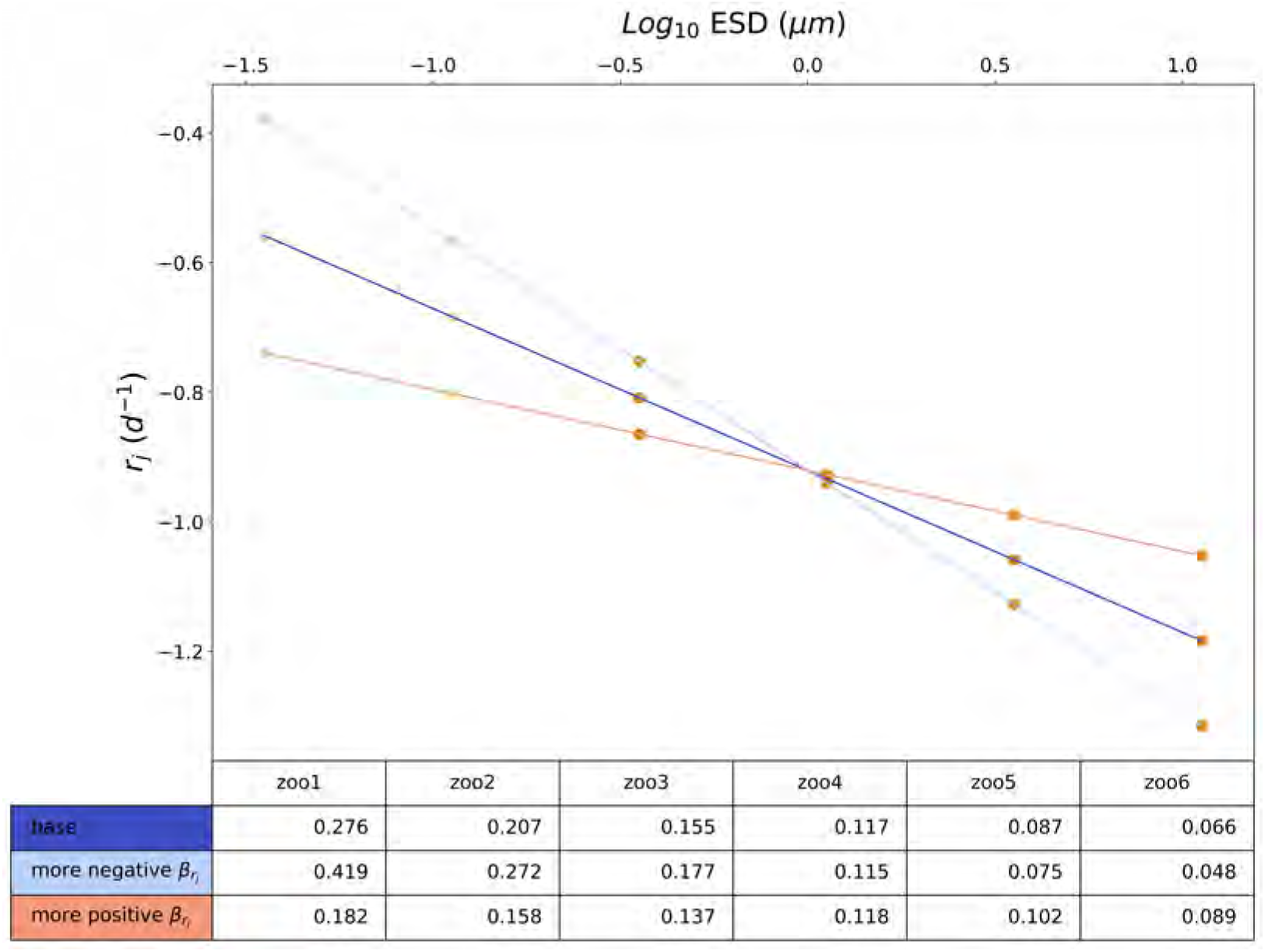
Sensitivity analysis of altering the allometric scaling exponent of the zooplankton basal respiration (*β_r_j__*). Deviations in *β_r_j__* fall ± 50%, within ranges of Hansen et al. (1997) and Kiørboe and Hirst (2014). The allometric relationship of *β_r_j__* in MARBL-SPECTRA (dark blue) is compared with a more negative (light blue) and a more positive *β_r_j__* (light red). The more negative scaling exponent (light blue) leads to higher basal respiration rates for the microzooplankton and the small mesozooplankton, and lower basal respiration rates for the largest three mesozooplankton. The more positive scaling exponent (light red) leads to lower basal respiration rates for the microzooplankton and the small mesozooplankton, and higher basal respiration rates for the largest three mesozooplankton. The table shows the exact *r_j_* values for each sensitivity test. The different colored dots represent the different functional types of zooplankton as in Figure 1.

**Figure S16:**
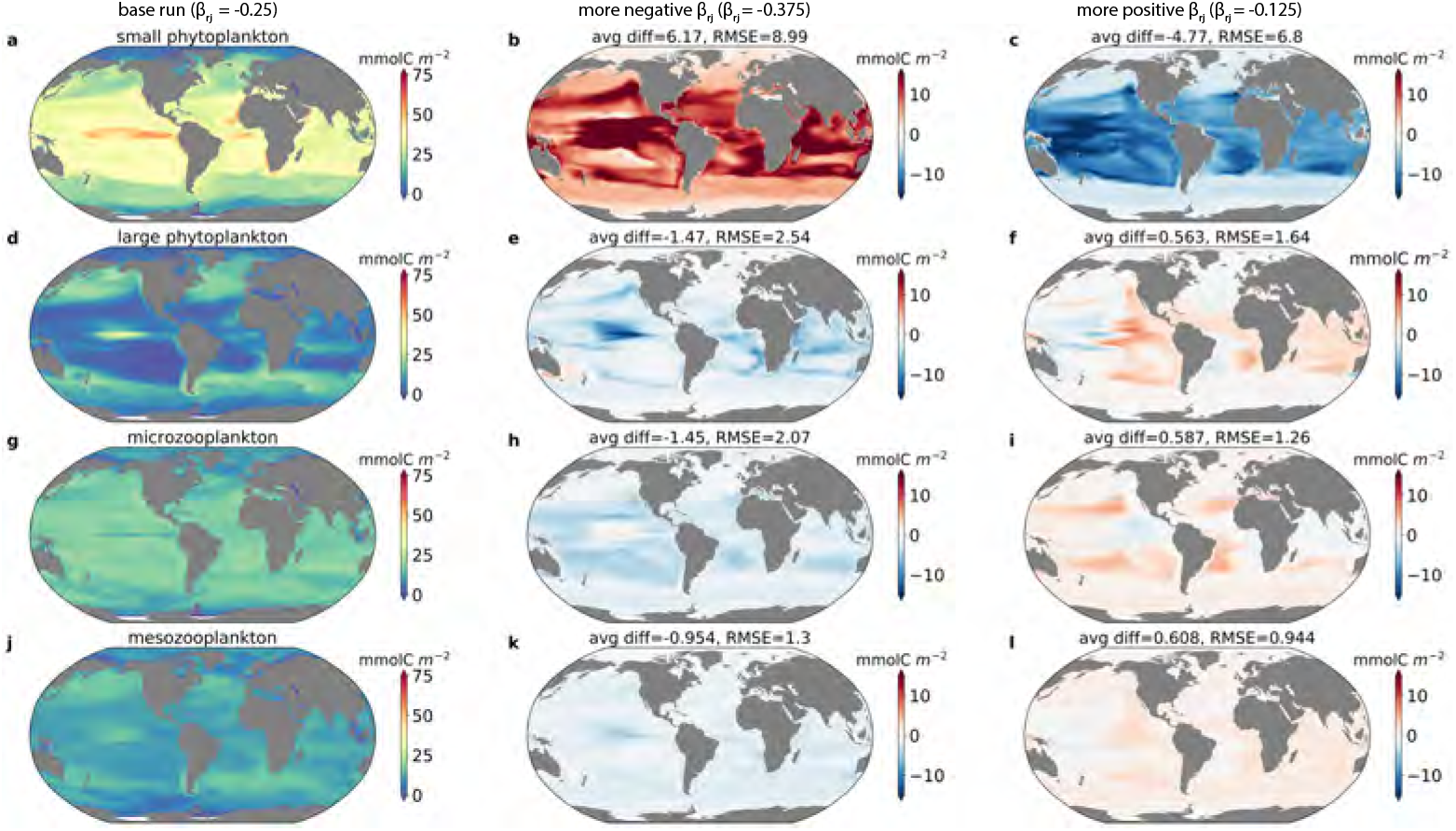
Depth-integrated average annual biomass for a) small phytoplankton (picoplankton, small mixed phytoplankton group, and diazotrophs), d) large phytoplankton (mixed phytoplankton 2, 3 and 4, and all diatoms), g) microzooplankton (smallest two zooplankton) and j) mesozooplankton (largest four zooplankton) in the base MARBL-SPECTRA run. b,e,h, and k) show the biomass differences between the base run minus the more negative slope run. c,f,i, and l) show the biomass differences between the base run minus the more positive slope run. In the base run (a,d,g,j), *β_r_j__* =-0.25 for all zooplankton. In the more negative slope run, *β_r_j__* =-0.375 and in the more positive slope run, *β_r_j__*)=-0.125 for all zooplankton. Avg diff is the average global difference in biomass and RMSE represents the root mean square error between the base run and each sensitivity run.

**Figure S17:**
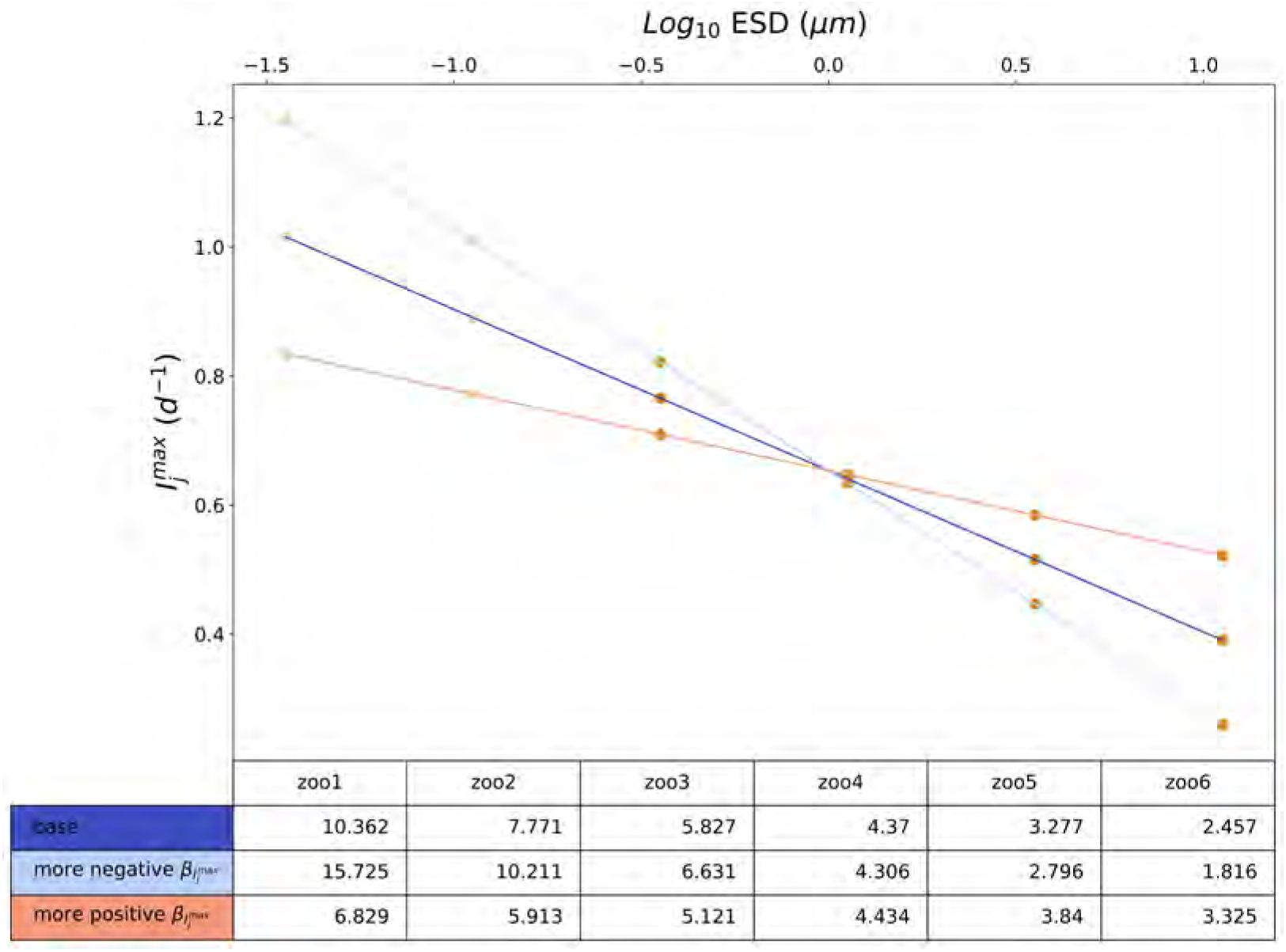
Sensitivity analysis of altering the allometric scaling exponent of the zooplankton maximum ingestion rate 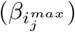. Deviations in 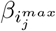 fall ± 50%, within ranges of Hansen et al. (1997) and Kiørboe and Hirst (2014). The allometric relationship of 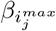 in MARBL-SPECTRA (dark blue) is compared with a more negative (light blue) and a more positive 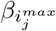 (light red). The more negative scaling exponent (light blue) leads to higher maximum ingestion rates for the microzooplankton and the small mesozooplankton, and lower maximum ingestion rates for the largest three mesozooplankton. The more positive scaling exponent (light red) leads to lower maximum ingestion rates for the microzooplankton and the small mesozooplankton, and higher maximum ingestion rates for the largest three mesozooplankton. The table shows the exact 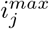 values for each sensitivity test. The different colored dots represent the different functional types of zooplankton as in Figure 1.

**Figure S18:**
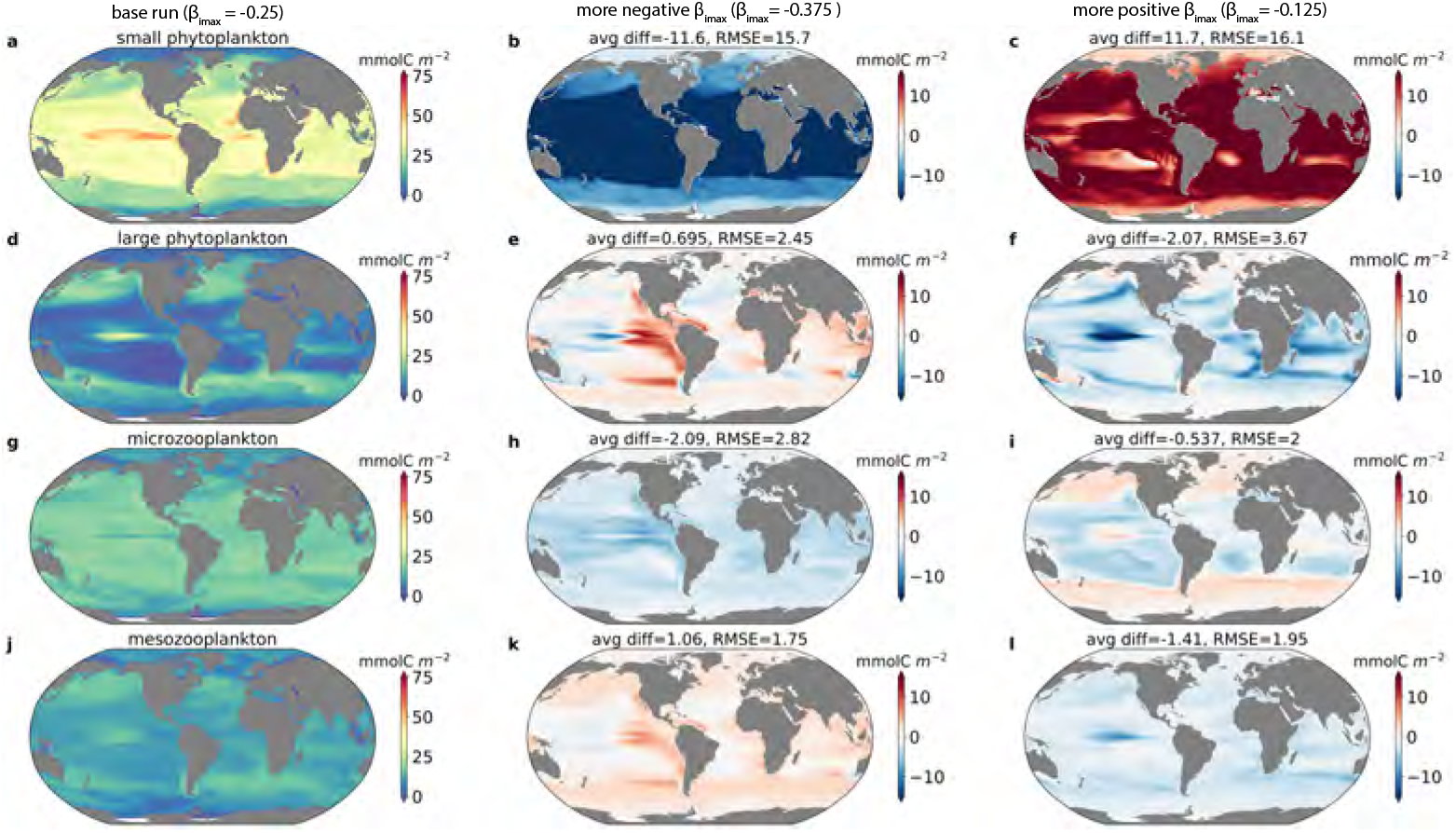
Depth-integrated average annual biomass for a) small phytoplankton (picoplankton, small mixed phytoplankton group, and diazotrophs), d) large phytoplankton (mixed phytoplankton 2, 3 and 4, and all diatoms), g) microzooplankton (smallest two zooplankton) and j) mesozooplankton (largest four zooplankton) in the base MARBL-SPECTRA run. b,e,h, and k) show the biomass differences between the base run minus the more negative slope run. c,f,i, and l) show the biomass differences between the base run minus the more positive slope run. In the base run (a,d,g,j), 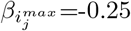 for all zooplankton. In the more negative slope run, 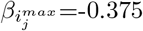, and in the more positive slope run 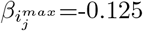 for all zooplankton. Avg diff is the average global difference in biomass and RMSE represents the root mean square error between the base run and each sensitivity run.

**Figure S19:**
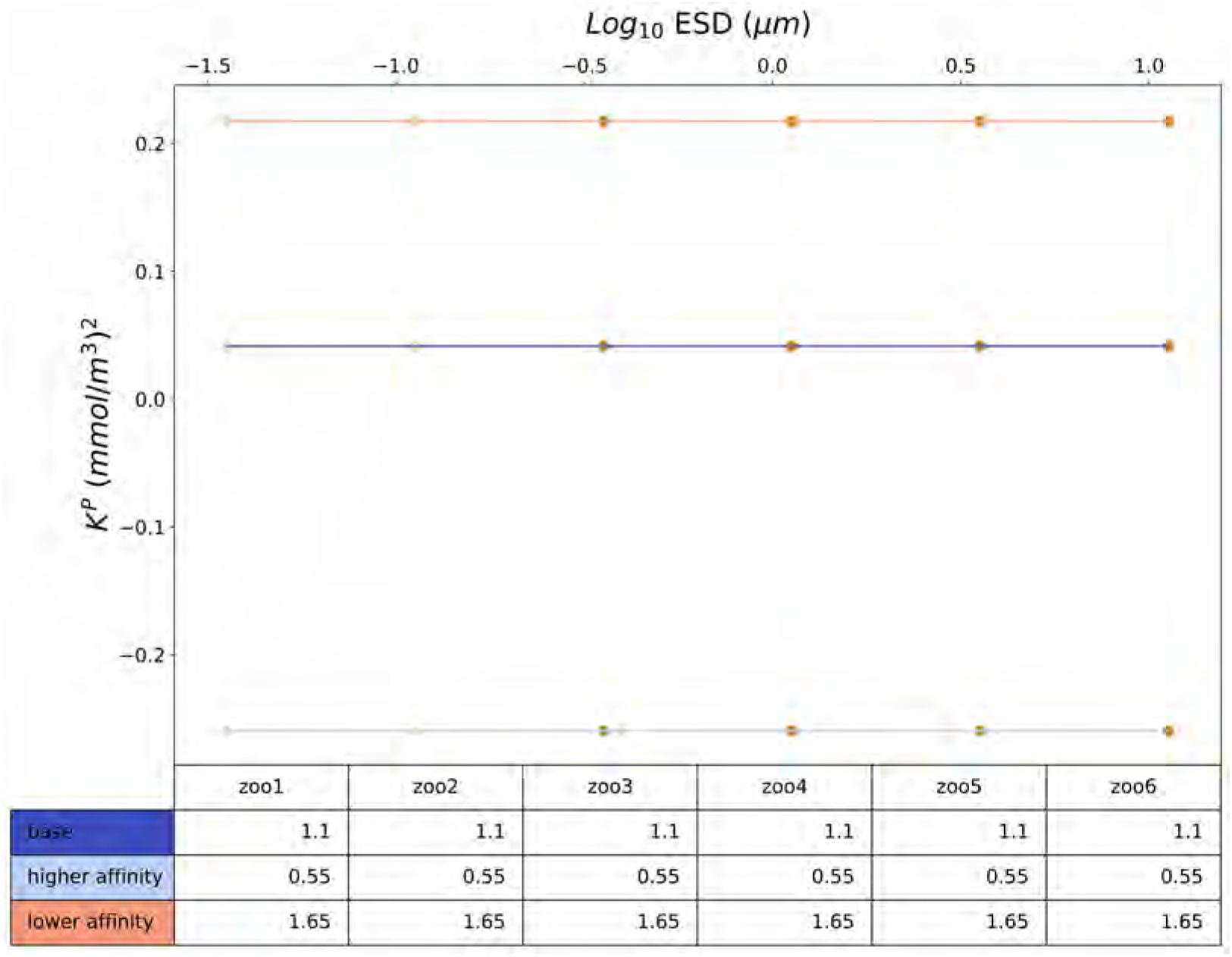
Sensitivity analysis of altering the grazing half saturation constant (*K^P^*). Deviations in *K^P^* fall ± 50%, within ranges of Hansen et al. (1997). *K^P^* in MARBL-SPECTRA (dark blue) is compared with a lower (light blue), and higher *K^P^* (light red). The lower *K^P^* (light blue) represents to higher grazing affinity for all zooplankton, compared to the higher *K^P^* (light red) representing lower grazing affinity. The table shows the exact *K^P^* values for each sensitivity test. The different colored dots represent the different functional types of zooplankton as in Figure 1.

**Figure S20:**
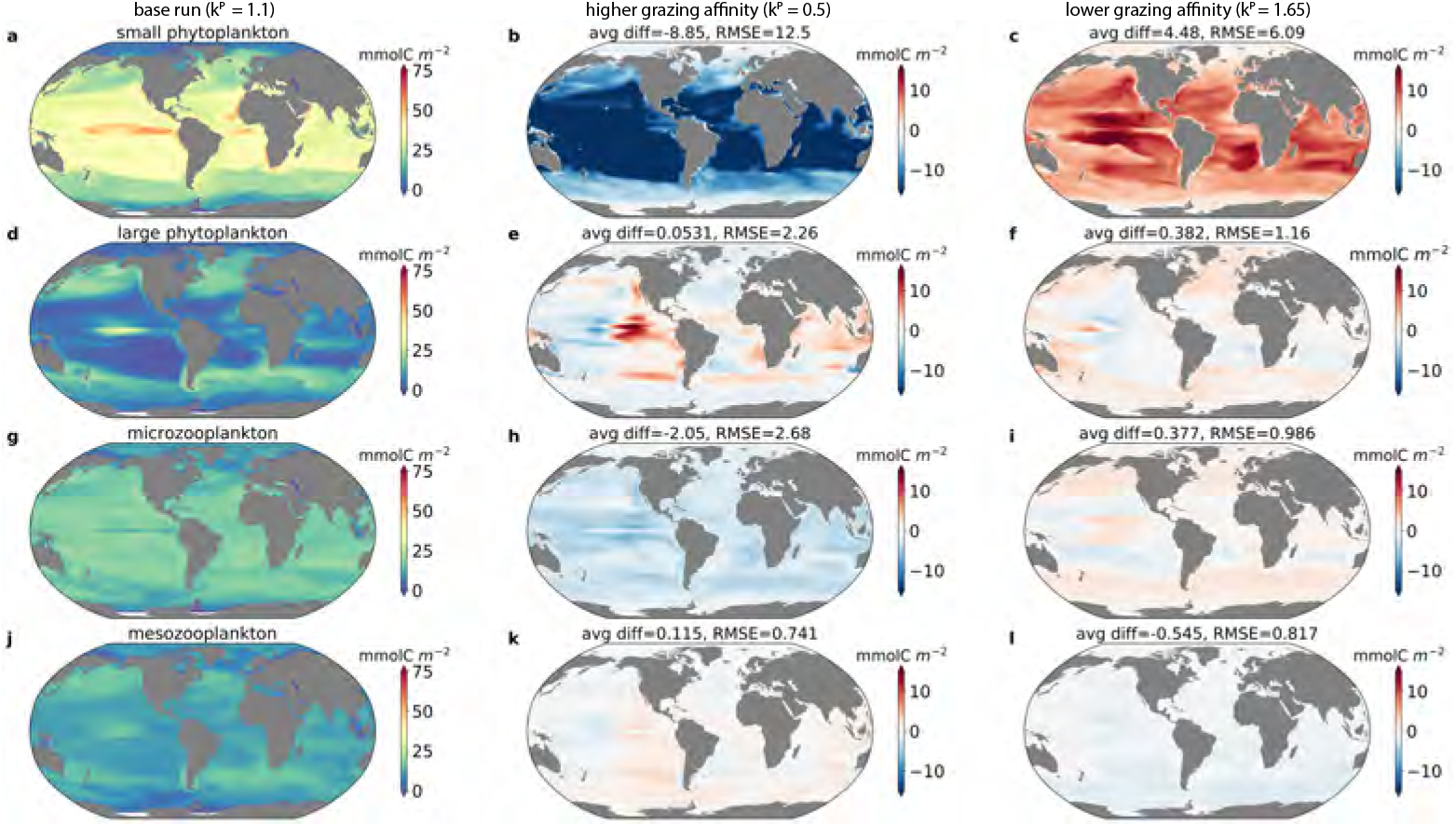
Depth-integrated average annual biomass for a) small phytoplankton (picoplankton, small mixed phytoplankton group, and diazotrophs), d) large phytoplankton (mixed phytoplankton 2, 3 and 4, and all diatoms), g) microzooplankton (smallest two zooplankton) and j) mesozooplankton (largest four zooplankton) in the base MARBL-SPECTRA run. b,e,h, and k) show the biomass differences between the base run minus the higher grazing affinity. c,f,i, and l) show the biomass differences between the base run minus the lower grazing affinity. In the base run (a,d,g,j) *K^P^*=1.1 for all zooplankton. In the higher grazing affinity run, *K^P^*=0.5, and in the lower grazing affinity *K^P^*=1.65 for all zooplankton. Avg diff is the average global difference in biomass and RMSE represents the root mean square error between the base run and each sensitivity run.

